# IMMUNE AND MOLECULAR CORRELATES OF RESPONSE TO IMMUNOTHERAPY REVEALED BY BRAIN-METASTATIC MELANOMA MODELS

**DOI:** 10.1101/2024.08.26.609785

**Authors:** Amélie Daugherty-Lopès, Eva Pérez-Guijarro, Vishaka Gopalan, Jessica Rappaport, Quanyi Chen, April Huang, Khiem C. Lam, Sung Chin, Jessica Ebersole, Emily Wu, Gabriel A. Needle, Isabella Church, George Kyriakopoulos, Shaojun Xie, Yongmei Zhao, Charli Gruen, Antonella Sassano, Romina E. Araya, Andres Thorkelsson, Cari Smith, Maxwell P. Lee, Sridhar Hannenhalli, Chi-Ping Day, Glenn Merlino, Romina S. Goldszmid

**Author notes:** Institute for Biomedical Research Sols-Morreale (IIBM), Spanish National Research Council-Universidad Autónoma de Madrid (IIBM, CSIC-UAM), Madrid, Spain. These authors contributed equally (Eva Pérez-Guijarro, Vishaka Gopalan).

## Abstract

Despite the promising results of immune checkpoint blockade (ICB) therapy, outcomes for patients with brain metastasis (BrM) remain poor. Identifying resistance mechanisms has been hindered by limited access to patient samples and relevant preclinical models. Here, we developed two mouse melanoma BrM models that recapitulate the disparate responses to ICB seen in patients. We demonstrate that these models capture the cellular and molecular complexity of human disease and reveal key factors shaping the tumor microenvironment and influencing ICB response. BR1-responsive tumor cells express inflammatory programs that polarize microglia into reactive states, eliciting robust T cell recruitment. In contrast, BR3-resistant melanoma cells are enriched in neurological programs and exploit tolerance mechanisms to maintain microglia homeostasis and limit T cell infiltration. In humans, BR1 and BR3 expression signatures correlate positively or negatively with T cell infiltration and BrM patient outcomes, respectively. Our study provides clinically relevant models and uncovers mechanistic insights into BrM ICB responses, offering potential biomarkers and therapeutic targets to improve therapy efficacy.

## INTRODUCTION

Cancer treatment has seen substantial improvement in the last decade. However, therapeutic options for patients with brain metastasis (BrM) are still limited and frequently ineffective, leading to a dismal prognosis^1,2^. Immune checkpoint blockade (ICB) has shown promising results in clinical trials, however, only a limited subset of patients respond to this approach^3–7^. While ICB combination or in association with radiotherapy, in particular stereotactic radiation therapy, can enhance efficacy, the use of steroids – often prescribed to BrM patients to reduced inflammation – has proven to be detrimental for ICB response^6^. Therefore, there is an urgent need to identify optimal combinations and regimens to improve patient outcomes. Moreover, we still lack a clear understanding of the specific tumor microenvironment (TME) responses in BrMs that could lead to the identification of biomarkers strongly correlating with the clinical benefit of ICB.

The brain tumor microenvironment (BrTME) is unique, containing highly specialized resident cell types including microglia that constitute the main immune population, and non-immune cells, such as neurons, astrocytes, and oligodendrocytes^8–10^. These are combined with specific extracellular matrix (ECM) networks and neurotransmitters^8–10^, all of which are essential for maintaining brain homeostasis. The BrTME is also controlled in part by the blood-brain barrier (BBB), which regulates the access and distribution of infiltrating immune cells, tumor cells, and drugs, to the brain^8–12^. Previous reports have shown clear differences in immune composition and tumor molecular profiles between BrMs, extracranial metastases, and primary tumors in patients^13–16^. This raises the question of whether determinants and mechanisms of ICB responses at extracranial sites also apply to BrM. Recent studies have expanded our knowledge regarding the diversity of the BrTME, demonstrating that heterogeneity in immune composition within the BrTME among patients is mainly driven by tumor cells themselves, rather than the uniqueness of the brain environment^17–22^. These studies showed that the largest discrepancies among BrM subtypes were in the T cell and myeloid compartments^17–22^. Nevertheless, how tumor cells regulate the BrTME and its impact on ICB responses remain to be elucidated.

Important limitations of these retrospective studies include the relatively small cohorts, the diverse treatments patients received, and the rare availability of biopsies from ICB responder patients. Additionally, the constraints of sample collection result in heterogeneous representation of the stroma in BrM biopsies, including an under-representation of microglia. Modeling BrM in mice is challenging. Some models rely on intracranial injection of tumor cells, bypassing extravasation and disrupting the BBB, thereby inducing local inflammation^23,24^. A few orthotopic BrM models, including autochthonous ones, recapitulate the full metastatic cascade^23–26^, though they are not without limitations such as low penetrance and/or long latencies^23,24^. Other models utilize systemic inoculation of tumor cells (e.g., intracarotid or intracardiac), recreating the cascade steps needed for brain colonization while maintaining BBB integrity^23,24^. However, these approaches also have drawbacks, including the need for invasive surgery (intracarotid inoculation) and the earlier development of extracranial lesions. The latter can hinder the comparative analyses of BrM-specific ICB-responders and non-responders, thus preventing the identification of predictive biomarkers of response and the discovery of new therapeutic targets.

The aim of this study was to address this unmet need and interrogate the immune and molecular determinants of ICB response in melanoma BrMs. To achieve this, we developed two syngeneic brain-metastatic models M4-BR1 (hereafter BR1) and M4-BR3 (BR3) derived from our previously established M4-B2905 (M4) melanoma mouse cell line^27^; both models exhibit the highest incidence of metastases and larger lesions in the brain, with BR1 being virtually brain-specific. We demonstrate that these models represent the molecular and cellular complexity and diversity of ICB responses observed in patients. Through comparative studies, we identify specific tumor molecular signatures associated with immune BrTME composition, ICB responses, and patient outcomes. Therefore, this study provides a robust framework for the necessary mechanistic studies to optimize BrM therapy.

## RESULTS

### Development of metastatic melanoma models with high brain penetrance

To study BrM therapeutic response to ICB, we generated a panel of syngeneic immunocompetent preclinical models preserving BBB integrity by intracardiac (i.c.) injection of the M4 melanoma mouse cell line^27^ into syngeneic C57BL/6 mice. We have previously demonstrated that M4, generated by neonatal UV-irradiation of hepatocyte growth factor transgenic mice (Hgf^Tg^), faithfully recapitulates the etiology, histopathology, and molecular features of human cutaneous melanoma^27^. After recovering metastatic cells from the brain, we cycled them *in vivo* to enhance their brain-metastatic potential (see Methods) (Figure S1A). We obtained two polyclonal sublines, BR1 and BR3, that exhibit high penetrance to the brain and form numerous pigmented macroscopic metastases in up to 100% of injected animals on average per experiment (Figure 1A-B). We confirmed the presence of melanoma cells in the brains of mice from both models by dopachrome tautomerase (DCT)^28,29^ immunostaining at day 18-20 post-implantation (model endpoint) (Figure S1B). To assess metastatic burden while preserving tissue integrity, we developed a machine learning approach that semi-automatically detects, counts, and measures the area of metastatic foci on whole-brain stereomicroscope images^30^. Following optimization (see Methods), we achieved a comparable number of BrMs per mouse between BR1 and BR3 (Figure 1C). However, BR3 formed more extracranial metastases and displayed a lower BrM/extracranial metastases ratio compared to BR1, indicating different metastatic tropism between the two models (Figures 1C, S1C-D). Nevertheless, in both BR1 and BR3, the highest incidence of metastases and larger lesions were observed in the brain, allowing for sufficient time to study BrMs without overwhelming extracranial disease (Figures S1C-D). Thus, we successfully developed two immunocompetent BrM models with high brain penetrance while maintaining a physiological BBB, providing a valuable platform for comparative and mechanistic studies of BrM therapeutic responses.

**Figure 1.**
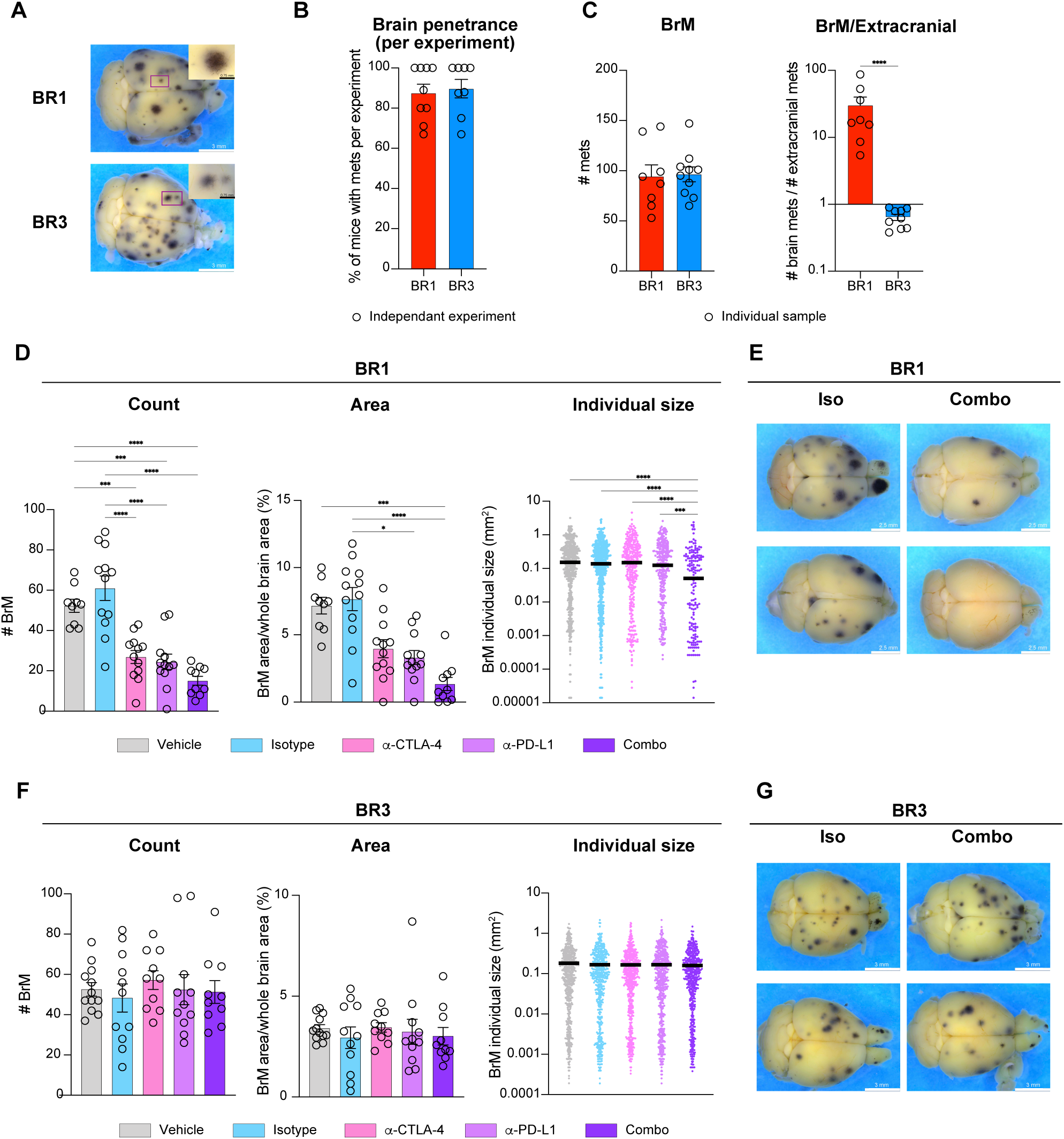
Generation of immunocompetent melanoma BrM models that exhibit ICB responses similar to brain metastases in patients. (A) Representative stereomicroscope images of whole brains (left) and zoomed-in image of BrM (inset) from BR1 and BR3 bearing animals. (B) Percentage of mice with BrM per experiment; dots depict independent experiments. (C) BrM count (left) and BrM/extracranial metastases ratio (right) per mouse; dots depict individual animals within an experiment. (D-G) BR1 or BR3 bearing animals injected with PBS (vehicle) or treated with isotype control (Iso), anti-CTLA-4 (α-CTLA-4), anti-PD-L1 (α-PD-L1), or anti-PD-L1 + anti-CTLA-4 combination therapy (Combo). (D) BR1 BrM count per individual brain (left), total brain metastatic area per individual brain (middle), and size of individual BrM lesion (right). (E) Representative brain stereomicroscope images from Iso- and Combo-treated BR1 bearing animals. (F-G) Same as D-E but for BR3 bearing mice. B-C shown as mean ± SEM, data from 9 (B) or 3 (C) experiments combined, n=3-12 mouse/group/experiment. D-G representative of 2 independent experiments for each model, data shown as mean ± SEM, n=9-12/group/experiment. *p <0.05, ***p < 0.001, ****p < 0.0001. See also Figure S1 and Table S1.

### BR1 and BR3 models exhibit mutational landscapes representative of human BrM melanomas

To confirm that our models are translationally relevant, we interrogated their molecular resemblance with human disease. We characterized the mutational landscape of cultured BR1 and BR3 cell lines in comparison to the M4 parental line using whole exome sequencing analysis. We found more mutated genes in the brain-metastatic lines compared with the parental M4 cells (432, 403, and 327 genes in BR1, BR3 and M4 parental, respectively; Figure S1E). Not surprisingly, 90% and 86% of the genes mutated in the M4 parental line were also detected in BR1 and BR3, respectively. However, each of the brain-metastatic models contained approximately 30% unique mutated genes (Figure S1E; Table S1), potentially contributing to their distinct metastatic behavior *in vivo*. Mutations related to PTEN or PI3K/AKT pathways were present across the M4 parental line, BR1, and BR3, although the specific mutated genes varied between these models (Table S1). We next explored the mutational landscape in metastatic patients using the publicly available MSK-MET-SKCM cohort^31^ and the cBioportal platform^32–34^. We found that most of the mutated genes identified in our models that are related with these pathways were also frequently mutated in melanoma patients with BrMs (Figure S1F). In line with our observation, while no specific mutations unique to melanoma BrM have been identified so far, AKT1^E17K^ increases BrM incidence in mice, and patients with mutations in the PI3K/AKT pathway (e.g., PTEN) have a higher risk of developing central nervous system (CNS) metastases^26,35,36^. Overall, these results show that our models display molecular alterations associated with BrM development in human melanoma.

### BR1 and BR3 BrMs recapitulate the diversity of responses to ICB observed in the clinic

To evaluate the ICB therapeutic response of the models, we treated syngeneic mice that were i.c.-injected with either BR1 or BR3 using anti-CTLA-4 or anti-PD-L1 monotherapies, or combination therapy (combo, anti-CTLA-4 + anti-PD-L1), dual isotype or vehicle controls. We started dosing at day 3 or 4 with each antibody, after verifying metastatic brain seeding on day 4 through microscopy and *ex vivo* culture of cells from the implanted brains (Figures S1G-H). When control groups reached the endpoint, we assessed BrM burden using our machine-learning algorithm^30^. We observed striking response differences between the two models (Figures 1D-G). We found a significant reduction in total metastatic foci count and area in BR1 following treatment with both mono- and combo-therapies (Figures 1D-E). Combo therapy was the most effective regimen, also reducing the size of individual metastases (Figures 1D-E). In addition, we observed significantly fewer extracranial metastases in this model, which disappeared in half of the monotherapy-treated and >90% of the combo-treated animals (Figure S1I). In marked contrast, BR3 BrMs were resistant to ICB, showing no changes in the total BrM foci count, metastatic area, or individual lesion size in any therapeutic group (Figures 1F-G). We observed a modest decrease in total extracranial metastasis counts in BR3-bearing animals treated with ICB monotherapies, and a significant response with combo therapy (Figure S1J). These results align with clinical studies indicating that extracranial and BrM responses to ICB do not always correlate with each other^3^.

Even at a more advanced metastatic stage (7 days after i.c. injection), the BR1 model remained significantly responsive to combo therapy and displayed a modest response to monotherapies (Figures S1K-L). Overall, our models recapitulate the diversity of clinical responses observed in melanoma BrM patients^3–5^.

### ICB-sensitive and resistant models display distinct BrTME and remodeling upon treatment

To identify potential immune correlates of BrM ICB responses, we performed multiparametric spectral flow cytometry on brain leukocytes from both combo-treated, which yielded the strongest response, and untreated BR1 and BR3 models. Consistent with their ICB responsiveness, we observed a significantly higher number of total leukocytes in BR1 compared to BR3, independent of their treatment status (Figure 2A). To gain more granularity, we performed unsupervised clustering of immune cells followed by supervised cell annotation (Figures 2B, S2A-B; Table S2). Uniform Manifold Approximation and Projection (UMAP) visualization and Principal Component Analysis (PCA) using the frequency of the identified leukocyte populations revealed different BrTME compositions between the two models in both treated and untreated groups (Figures 2C-E).

**Figure 2.**
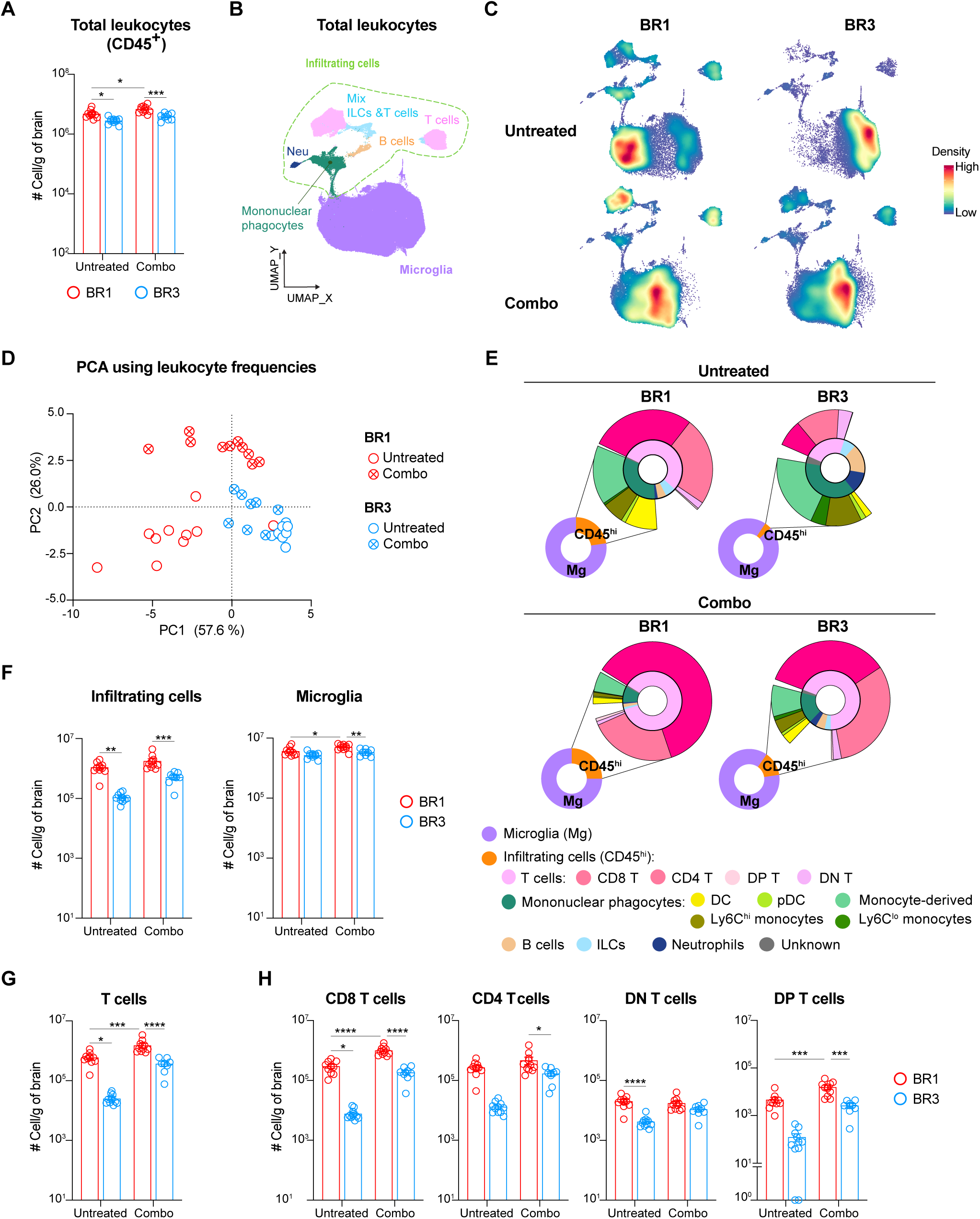
Distinct BrTME composition is associated with different responses to ICB. High-parametric spectral flow cytometry analysis of brain leukocytes from BR1 and BR3 bearing animals untreated or combo treated. (A) Absolute number of leukocytes (live CD45^+^) per gram of brain tissue. (B) UMAP projection of total BrM leukocytes. Main populations indicated: resident microglia, infiltrating mononuclear phagocytes, neutrophils (Neu), B cells, innate lymphoid cells (ILCs), and T cells. (C) Density plots showing cell distribution in untreated (top) and treated (bottom) mice from BR1 (left) and BR3 (right) models. (D) Principal Component Analysis (PCA) of main immune cell population frequencies in total leukocytes for each sample. (E) Proportion of microglia (Mg; purple) and infiltrating cells (CD45^hi^; orange) in total leukocytes. Proportion of indicated populations within total infiltrating cells (inset). (F) Absolute number of infiltrating cells (left) and microglia (right) per gram of brain. (G) Absolute number of total T cells per gram of brain. (H) Absolute number of indicated T cell subsets per gram of brain. A, F, G, and H data shown as mean ± SEM, n = 8-10/group/experiment. *p <0.05, **p < 0.01, ***p < 0.001, ****p < 0.0001. See also Figure S2 and Table S2.

To determine the specific immune populations driving the differences between untreated BR1 and BR3, we analyzed the PC1 loadings, which segregate these two groups (Figures 2D and S2C-D). These loadings revealed that BR1 samples had a higher frequency of various infiltrating cells, including T cells, DCs, natural killer (NK) cells, and inflammatory monocytes (Ly6C^hi^), which are known to favor ICB responses^37–39^ (Figures 2E and S2D-F). The absolute number of these cells was also higher in BR1 compared to BR3 (Figures 2F-H and S2G). This indicates a stronger immune cell recruitment in untreated BR1 BrMs compared to untreated BR3. In contrast, BR3 samples had a higher frequency of neutrophils, microglia, and Ly6C^lo^ monocytes (Figures 2E and S2D). Both the proportion of microglia (>90% of total leukocytes) and the infiltrating cell populations in the BR3 ICB-resistant BrTME resembled that of non-tumor-bearing brains (PBS) (Figures 2E and S2H).

We observed a significant therapy-induced increase of total leukocytes along with an enrichment of T cells in BR1 and a trend toward an increase of infiltrating T cells in BR3 (Figures 2A, 2E-H, S2E-F). This aligns with recently published clinical studies showing ICB-driven increase T cell infiltration^21,40,41^. Nevertheless, TME dissimilarities between BR1 and BR3 were still observed after combo therapy, as evidenced by their segregation along PC2 (Figures 2D, S2D and S2I). BR1 continued to display higher frequency and number of infiltrating cells compared to BR3, which after combo therapy was specifically due to a surge in CD8^+^ T cells (Figures 2E-H, S2F-G). Additionally, a higher number of microglia was observed in treated BR1 compared to treated BR3 (Figure 2F). Altogether, these observations indicate BrTME immune compositional differences between ICB-sensitive and -resistant BrMs, with some differences, such as those in T cells and microglia, persisting after therapy.

### A distinct T cell landscape is associated with ICB response

The quantity and characteristics of infiltrating T cells play an important role in the response to ICB^42–45^. Therefore, we studied the specific T cell characteristics associated with ICB-responses in our BrM models by performing single-cell RNA sequencing (scRNAseq) of whole brains from PBS and untreated BR1 and BR3 samples (Figures S3A-B). Differential gene expression analysis of total CD8^+^ T cells between ICB-sensitive BR1 and resistant BR3 revealed higher expression of activation/effector function-associated genes (*Nkg7*, *Pdcd1*, *Ifng*, *Gzmb*) in BR1, whereas the BR3 CD8^+^ T cell pool displayed a naïve signature (*Sell*, *Lef1*, *Tcf7*, *Foxp1*) (Figure 3A; Table S3). UMAP projection and unsupervised clustering of the T cells revealed six CD8^+^ T cell clusters (CD8.sc1-6) and six CD4^+^ T cell clusters (CD4.sc1-6) with different abundance between the two models (Figures 3B-C, S3C). Clusters CD8.sc1 and CD4.sc1, defined by a naïve gene signature, were enriched in PBS and BR3 samples, accounting for over 60% of their total CD8^+^ and CD4^+^ T cell populations, whereas they made up only 5-20% of T cells in BR1 (Figures 3B-D, S3C-E; Tables S3-S4). Projection of our T cell clusters onto a single-cell reference atlas for T cell states^42^ confirmed that CD8.sc1 and CD4.sc1 correspond to the naïve compartments (Figures 3E, S3F). BR3 also presented a higher proportion of cluster CD8.sc3, which mapped mostly onto CD8^+^ effector memory T cells on the reference atlas (Figures 3B-E, S3D; Table S3).

**Figure 3.**
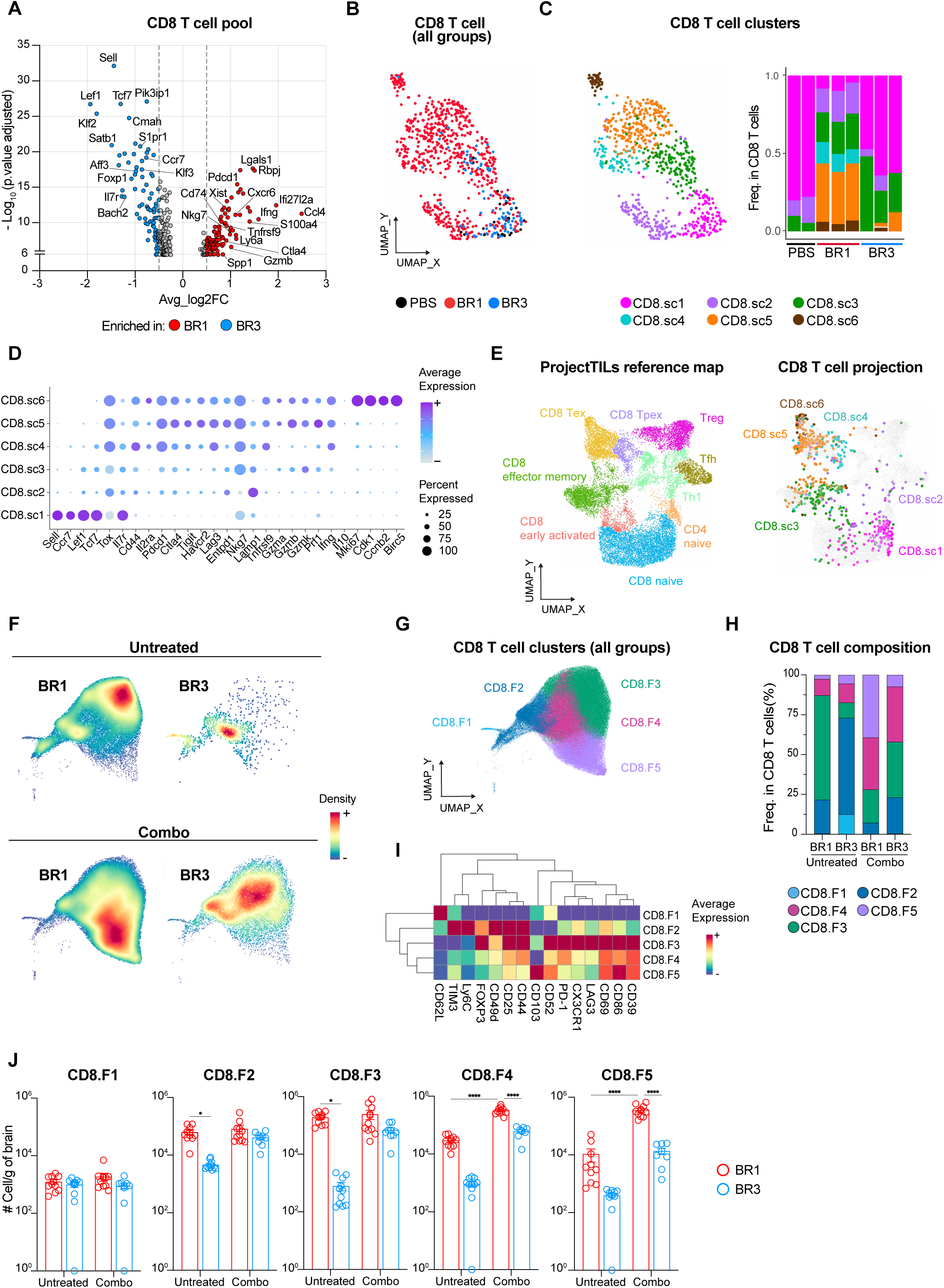
Distinct CD8^+^ T cell subsets infiltrate ICB-responder compared to ICB-resistant BrMs. (A-E) scRNA-seq analysis of CD8^+^ T cells from untreated BR1, BR3, and PBS brains. (A) Volcano plot of BR1 versus BR3 differentially expressed genes (DEGs). Significant DEGs are indicated (BR1: red, BR3: blue) (B) UMAP projection showing cell distribution in each group. (C) CD8^+^ T cell clustering. UMAP projection (left) and proportion of each cluster per sample (right). (D) Dot plot showing the expression of selected genes among clusters. (E) Projection of CD8^+^ T cell clusters (right) onto the single-cell reference atlas (left) for T cell states ProjecTILs. (F-J) High-parametric spectral flow cytometry analysis of CD8^+^ T cells from indicated samples. (F) Density plots showing cell distribution in untreated (top) and treated (bottom) BR1 (left) and BR3 (right) models. (G) UMAP projection and clustering of indicated CD8^+^ T cell clusters among total CD8^+^ T cells. (H) Proportion of indicated CD8^+^ T cell clusters in total CD8^+^ T cells. (I) Heatmap depicting expression level of indicated surface markers (scaled by marker) in each CD8^+^ T cell cluster. (J) Absolute number of indicated T cell clusters per gram of brain. A-E n = 2-3/group. F-J n = 8-10/group/experiment. J data shown as mean ± SEM, *p <0.05, ****p < 0.0001. See also Figure S3 and Tables S3-S4.

In contrast, BR1 was characterized by a higher diversity of T cells, including various CD8^+^ and CD4^+^ T cell subsets expressing different levels of genes encoding T cell degranulation/lysosomal activity (*Lamp1*, *Ctsl, Cd63;* CD8.sc2), activation/dysfunction (*Cd81*, *Ckb*, *Nr3c1;* CD8.sc2)^46,47^, regulatory function (*Foxp3, Il2ra, Ikzf2;* CD4.sc5-6), and immune checkpoint receptors previously described as activation and/or exhaustion markers (*Lag3*, *Pdcd1*, *Havcr2;* CD8.sc4-6 and CD4.sc3-4)^42,45,46^ (Figures 3B-E, S3C-E; Tables S3-S4). Among the latter, CD8.sc5, the most enriched cluster in BR1, displayed the highest level of genes encoding cytotoxic molecules (e.g., *Ifng*, *Prf1*, *Gzmb*) (Figures 3B-D, S3D; Table S3). Similarly, the BR1-enriched CD4-sc3 exhibited the highest expression of *Ifng* and *Tbx21* (T-bet), consistent with a Th1-like signature (Figures S3C, S3E; Table S4). Notably, CD8.sc4, exclusively found in the BR1 BrTME, expressed the transcription factor TCF-1 (*Tcf-7*), the nuclear protein TOX (Tox), and low levels of genes encoding cytotoxic cytokines and immune checkpoint receptors (except *Havcr2*) (Figures 3B-E; Table S3). This expression profile resembles that of the so-called “exhausted stem cell-like CD8 T cells”, previously identified as essential for ICB response and associated with better clinical outcomes in melanoma patients^43,45^. Finally, the BR1 enriched clusters CD8.sc6 and CD4.sc6 displayed a proliferative profile (*Mki67*, *Cdk1*, *Ccnb2*), suggesting that local expansion contributes to the higher T cell number observed in this model (Figures 3B-D, S3C-E; Tables S3-S4). We confirmed these T cell compositional differences between untreated BR1 and BR3 BrTME by spectral flow cytometry and showed higher number of activated T cells in BR1 (Figures 3F-J, S3G-I).

To determine if the T cell compositional differences between the two models are sustained after treatment, we analyzed the BrTME of combo-treated BR1 and BR3 (Figures 3F-J, S3G-I). BR1 showed a specific 20-fold increase in CD8.F5 and a 10-fold increase in CD8.F4, both subsets marked by activation (Figures 3H-J). Interestingly, CD8.F5 resembles tissue-resident memory T cells (T_RM_), which have been associated with better ICB efficacy and improved patient outcomes for non-brain tumors^48–50^ (Figure 3I). In addition, combo therapy in BR1 led to a significant increase in the CD4^+^ T cell subset CD4.F5 displaying activation/exhaustion markers (Figures S3H-I). On the other hand, most CD8^+^ and CD4^+^ T cell subsets slightly increased in BR3 after therapy (Figures 3J, S3I). Together, these results demonstrate that the quantity and the quality of the T cells differed between the ICB-responder and resistant BrM models, highlighting the importance of the intrinsic BrTME T cell landscape in ICB response and aligning with immune correlates of response described for tumors at extracranial sites^43–45^.

### Unique reactive microglia signatures are found in ICB-responder BrMs

Microglia, the predominant immune cell type in the CNS, are crucial in regulating brain homeostasis, response to tissue injury, and pathogens^51–53^. While different studies have suggested their dual role in BrM development^10,54–56^, their contribution to ICB response is unclear. As expected, microglia represented 80-95% of the immune BrTME in both of our models (Figures 2E, S2E). We then inquired whether different microglia transcriptional states might be associated with BrM response to ICB. To do so, we extracted microglia from our whole-brain scRNA-seq dataset (Figure S3B) and performed differential gene expression and Gene Set Enrichment Analysis (GSEA). Compared to BR3, BR1 microglia upregulated genes and pathways associated with MHC class I and II antigen presentation (MHC-I/-II) (*H2-Aa*, *Cd74*, *B2m*), response to type I and II interferons (IFN-I/ -II) (*Ifitm3*, *Ifi27l2a*, *Cxcl10*), and inflammatory proteins including T cell-attracting chemokines (*Cxcl10*, *Cxcl16*, *Cxcl9*) (Figures 4A-B), indicating a reactive microglia transcriptional signature. Similar pathways were enriched in BR1 compared to PBS (Figure S4A). Conversely, BR3 microglia showed enrichment of homeostatic genes and pathways such as neuronal development and synaptic remodeling^51–53^, displaying a transcriptional profile similar to PBS (Figures 4A-B, S4B). TNFα signaling via NF-κB was enriched in BR3, but not BR1, microglia compared to PBS (Figure S4A).

**Figure 4.**
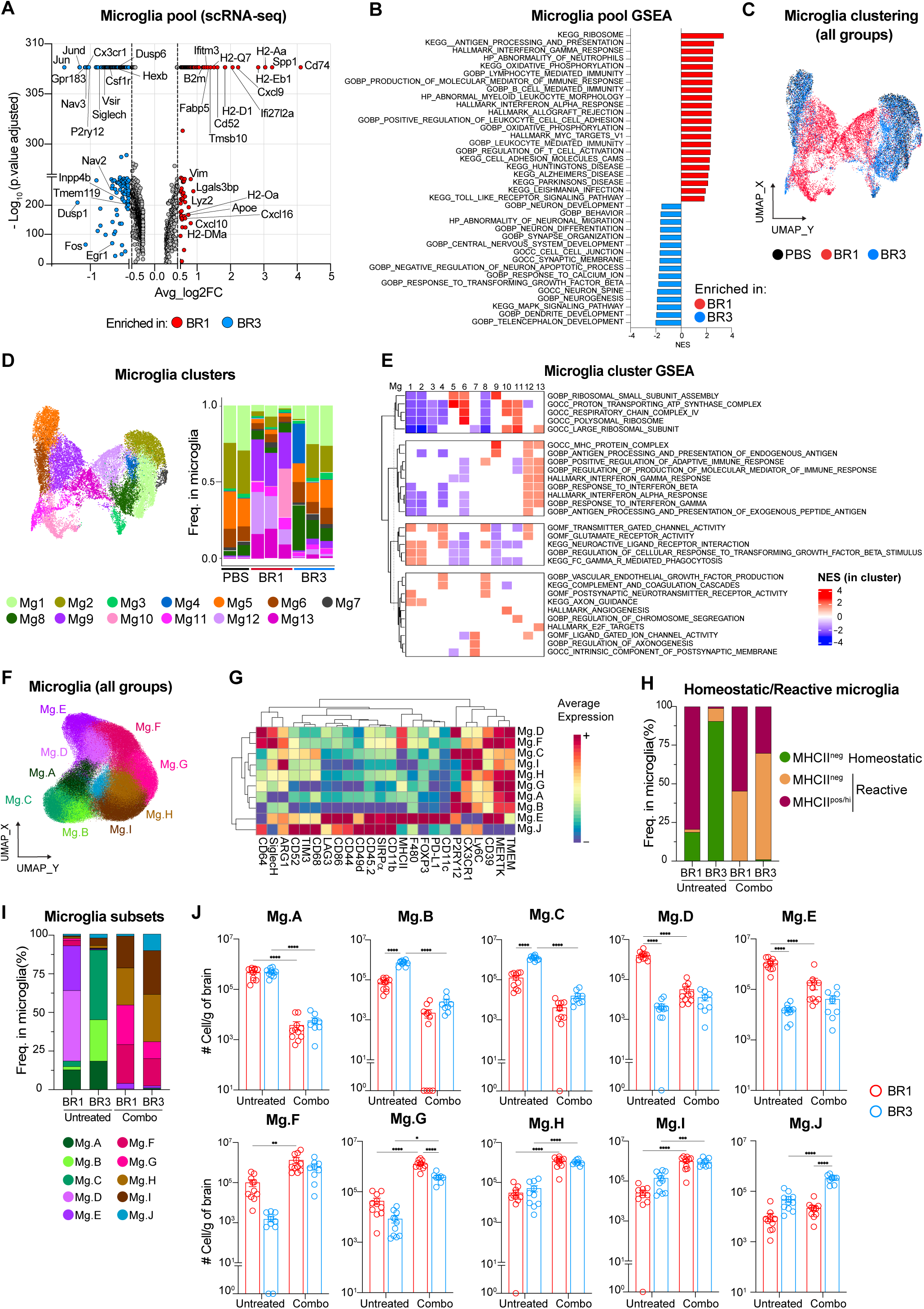
Unique reactive microglia signatures are found in ICB-responder BrMs. (A-E) scRNA-seq analysis of microglia from untreated brains. (A) Volcano plot of BR1 (red) versus BR3 (blue) DEGs. (B) Gene set enrichment analysis (GSEA) showing selected pathways among the top 10 Hallmark (HP), Gene Ontology (GO), and Kyoto Encyclopedia of Genes and Genomes (KEGG) pathways enriched in BR1 (red) microglia versus BR3 (blue) (see Methods). NES: Normalized Enrichment Score. (C) UMAP projection of total microglia showing cell distribution among groups. (D) UMAP projection clustering (left) and proportion per sample (right) of indicated microglia clusters in total microglia. (E) GSEA showing selected HP, GO and KEGG top pathways enriched in each microglia cluster. NES: Normalized Enrichment Score in each cluster. (F-J) High-parametric spectral flow cytometry analysis of microglia from untreated or combo-treated BR1 and BR3 brains. (F) UMAP projection and unsupervised clustering. (G) Heatmap representing expression level of indicated surface markers (scaled by marker) for each cluster with hierarchical clustering indicated on left and markers on top. (H) Proportion of indicated microglia states in total microglia per group. (I) Proportion of indicated microglia clusters in total microglia per group. (J) Absolute number of indicated microglia clusters per gram of brain. A-E n = 2-3/group/experiment. F-J n = 8-10/group/experiment. J data shown as mean ± SEM, n = 8-10/group/experiment. *p <0.05, **p < 0.01, ***p < 0.001, ****p < 0.0001. See also Figure S4.

Microglia are heterogeneous, particularly in disease conditions^51–53^. To investigate microglia heterogeneity in our BrM models, we performed unsupervised clustering and identified thirteen different microglia clusters (Figures 4C-D). Clusters Mg1-4 and Mg8, characterized by the highest level of homeostatic microglia genes and pathways (e.g., phagocytosis, regulation of cellular response to TGFβ, and neuronal function), represented the predominant microglia states in BR3 and PBS (Figures 4C-E, S4C). Clusters Mg5-6, also highly abundant in these groups, were enriched in pathways related to ribosomal and mitochondrial function, indicating a transcriptionally active state (Figures 4C-E). In contrast, the overall reactive signature of BR1 microglia was driven by the combination of five different clusters, Mg9-Mg13, defined by reduced expression of homeostatic genes and pathways (Mg9-11 and Mg13), upregulation of IFN response genes (Mg12-13), antigen presentation (Mg9, Mg12-13), and pathways involved in T cell recruitment and regulation of adaptive immune responses (Mg12-13) (Figures 4C-E, S4C). These clusters constituted 70-80% of the total BR1 microglia, while representing less than 15% of BR3 and minimally present in PBS (Figure 4D). Clusters Mg11 and Mg13, nearly absent in BR3 and PBS, expressed mitotic and proliferative genes (Figures 4C-E, S4C), possibly explaining the higher number of microglia found in BR1 (Figure 2F). Using high-parametric spectral flow cytometry, we confirmed the higher proportion and total number of microglia with various reactive states in BR1 (Mg.D-E) as compared to BR3 (Figures 4F-J, S4D). Taken together, our data suggest that reactive microglia, characterized by high expression of MHC-I/II and T cell-attracting chemokines, are associated with response to ICB, while resistant BrMs maintain homeostatic microglia states.

### Systemic ICB alters microglia functional state

It is unclear whether systemic ICB treatment can impact microglia states. To address this question, we assessed the combo-treated BR1 and BR3 microglia phenotype using spectral flow cytometry. Surprisingly, therapy led to the disappearance of almost all homeostatic microglia states in both models (Mg.A-Mg.C; Figures 4F-J, S4D). Instead, they were replaced by various non-homeostatic states characterized by the absence of P2RY12 expression (Mg.F-Mg.J; Figures 4F-J, S4D), the loss of which has been previously linked with neuroinflammation^52,57^. While the decrease of P2RY12 was a common therapy-induced feature, some phenotypic differences were still observed between BR1 and BR3 (Figures 4G-I, S4D). BR1 microglia maintained the expression of MHC-II, while reducing markers associated with macrophage pro-tumorigenic function such as ARG1, SIRPα^58–61^ and the T cell inhibitory receptors PD-L1, TIM3 and LAG3^62–64^ (clusters Mg.F-G; Figures 4G). In contrast, the microglia state increased in BR3, but not in BR1, after therapy (Mg.J) expressed the highest levels of TIM3, LAG3, and SIRPα (Figures 4G, 4I-J, S4D). We cannot rule out that the low levels of PD-L1 detected after combo treatment stem from a masking effect of anti-PD-L1 antibody. Combined, these results indicate that systemic ICB therapy alters microglia functional states. The key differences between microglia in ICB-responder and non-responder BrMs, regardless of treatment status, lie in the expression of genes, pathways, and proteins that can modulate T cell activities.

### BR1 and BR3 exhibit distinct tumor cell-intrinsic molecular programs that correlate with those present in human BrMs

We next sought to identify tumor cell-intrinsic molecular features that may be driving the distinct BrTME observed in our models. To do so, we compared BR1 and BR3 melanoma cell transcriptomes from our whole brain scRNA-seq data set. Model-based analysis of single-cell transcriptomics (MAST)^65,66^ followed by GSEA revealed that overall BR1 cells were enriched in MHC-I/II antigen presentation (e.g., *H2-K1/D1, B2m, Tap1, H2-Aa/Ab1/Eb1, Cd74,*), immune response and IFN signaling (e.g., *Ifng, Irf1, Stat1, Ifi203, Cxcl9*), cell cycle, and DNA replication and repair (e.g., *Mki67, Aurka/b, Plk1, Top2a, Pdgfa, Chek1*) (Figures 5A, S5A). On the other hand, BR3 cells expressed higher levels of genes and pathways related to melanin biosynthesis (e.g., *Tyr, Dct, Slc45a2*), neuronal function (e.g., *Sema3a, Unc5c, Slc6a11, Pde1c/3a/4b, Cacna1d, Selenop*), and metabolic pathways such as glutathione transferase activity (e.g., *Gsta4, Gstm1*), oxidative phosphorylation (OXPHOS, e.g., *Atp6v0d2, Cox7a1*), and lipid metabolism (e.g., *Apoe*) (Figures 5A, S5A).

**Figure 5.**
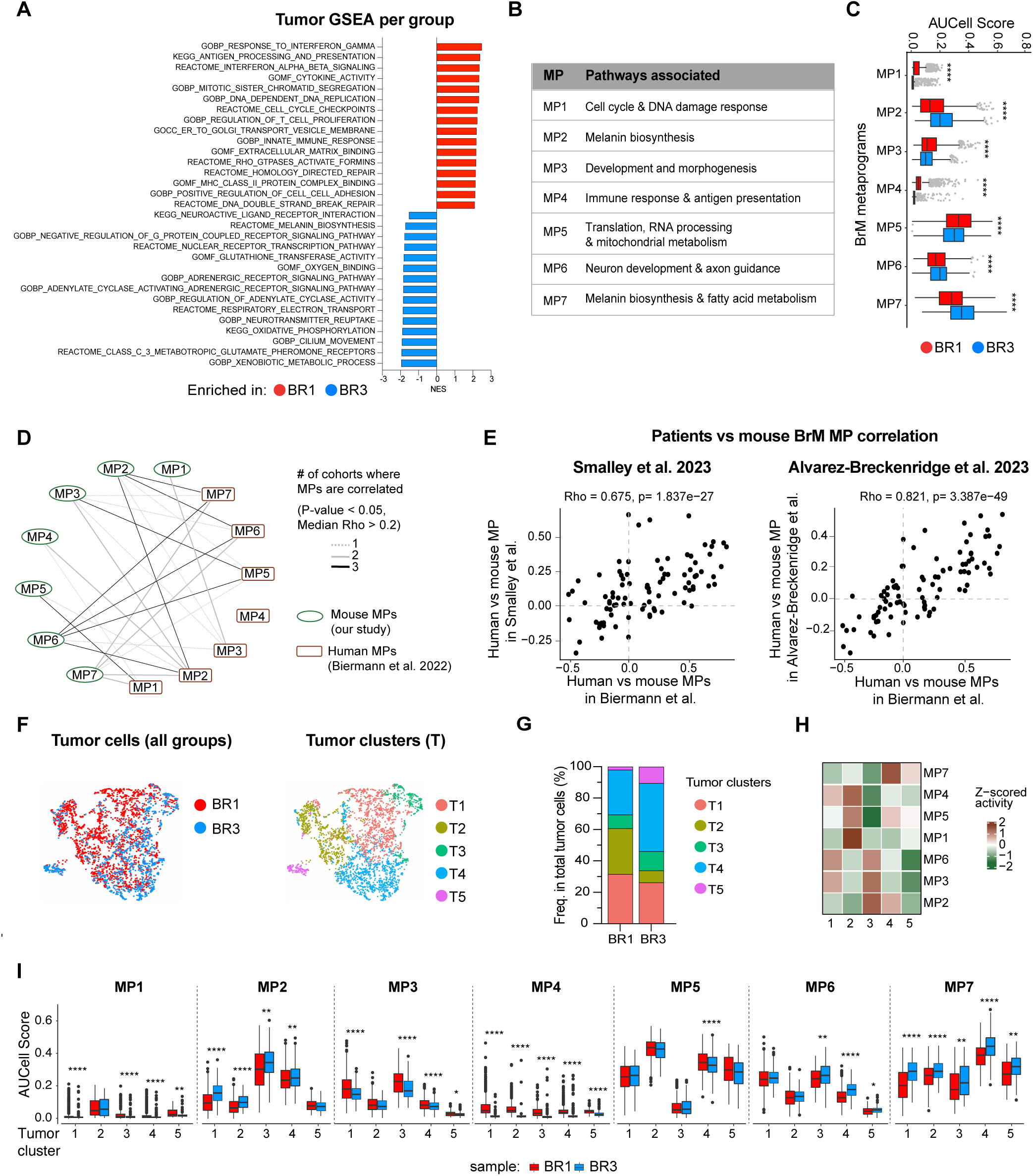
BR1 and BR3 tumor cells differentially express molecular programs that are present in human BrM tumor cells. scRNA-seq analysis of tumor cells from untreated BR1 and BR3 brains. (A) GSEA showing selected GO, KEGG and REACTOME pathways enriched among the BR1-specific and BR3-specific genes (see Methods). (B) Metascape-associated pathways for BrM metaprograms (MP) identified in BR1 and BR3 tumor cells. (C) Per cell basis analysis of MP activity per group. (D) Correlation-based network analysis between mouse MPs derived in our study (green circles) and human MPs from Biermann et al. 2022 (brown rectangles), using three patient cohorts. Edge thickness and color represent the number of cohorts in which each MP pair is correlated. (E) Spearman correlation between mouse versus human MPs activities in the indicated data sets. x- and y-axis depict spearman correlation coefficient values (F) UMAP projection of tumor cells based on Z-scores of MP activities showing BR1 (red) and BR3 (blue) cell distribution (left) and tumor clusters distribution (right). (G) Proportion of tumor clusters per group. (H) Heatmap depicting Z-score of MP mean activity across malignant cells in each cluster (columns). (I) Per cell basis analysis of MP activity within each cluster from each group. A-C and F-I n = 2-3/group, D-E n = 2-3 mouse/group (our study), n= 25 (Biermann et. al), n=14 (Smalley et. al), n=27 (Alvarez-Breckenridge et. al). See also Figure S5 and Table S5. *p <0.05, **p < 0.01, ***p < 0.001, ****p < 0.0001.

To determine the key molecular programs (referred to as metaprograms [MPs]) that characterize our models and compare them with available clinical data, we applied a Non-negative Matrix Factorization (NMF)^67^ approach (Figure S5B; see Methods). NMF-based methods are able to identify gene expression patterns that highlight differences within tumors, as well as variation between different tumors^68–70^. A similar approach has been successfully used in BrM patients to discover tumor MPs^13^. We identified seven different MPs (MP1-MP7) and characterized them using Metascape pathway analysis^71^ (Figure 5B; Table S5). Melanoma cells within BR1 tumors were enriched in MPs associated with cell cycle and DNA damage response pathways (MP1), tissue development and morphogenesis (MP3), immune response and antigen presentation (MP4), and translation and mitochondrial metabolism (MP5) (Figure 5C). Melanoma cells in BR3 tumors displayed higher activity of MPs related to melanin biosynthesis (MP2 and MP7) and neuron development and axon guidance (MP6) (Figure 5C). To validate that our mouse BrM MPs are representative of human disease, we computed correlations between these mouse MPs and human MPs from three melanoma patient cohorts^13,40,72^ (Figure 5D-E; see Methods). Each mouse MP identified in our study correlated with one or more patient BrM MPs involved in similar functions from at least one cohort (Figure 5D). We verified that the correlations found between pairs of human- and mouse-MPs were consistent across all melanoma BrM human datasets^13,40,72^ (Figure 5E). We did not observe significant correlations between mouse MPs and human MPs exclusively expressed in extracranial metastases (i.e., human-MP4)^13^ (Figure 5D).

Given that the BR1 and BR3 cell lines are polyclonal, we investigated whether the transcriptional signatures distinguishing these two models were broadly expressed across all cells within each model or specific to certain cell subpopulations. We performed unsupervised clustering of tumor cells based on their expression of MPs, identifying 5 clusters (T1-T5) with distinct MP activity that were differentially distributed between the two models (Figures 5F-I, S5C). The main differences between BR1 and BR3 were the higher proportion of T2 in BR1, which displayed the highest activity for MP1, MP4 and, MP5, whereas BR3 had higher frequency of T3, T4, and T5, expressing MP2, MP3, MP6, and MP7 (Figures 5G-I). Analysis of MP activity on a per-cell basis revealed that in some cases, MP expression varied between groups even within the same cluster, for example, MP6 was expressed at a higher level in BR3 cells in clusters 3, 4, and 5 (Figure 5I). These results suggest that the difference between BR1 and BR3 tumor cells and their overall MP expression can be attributed to variation in cluster abundance, the specific MP activity within these clusters, and their activity at the per-cell level. In summary, we identified molecular programs distinguishing ICB-responder and ICB-resistant BrM models that may contribute to their distinct BrTME and therapy responses. Furthermore, we demonstrated that these MPs are also present in human melanoma BrMs, highlighting the translational relevance of our models.

### BR1 and BR3 tumor cell-intrinsic features shape the microglia functional states that drive T cell phenotypes

Cell-cell interactions are essential to shape the TME and drive antitumor immunity. Based on the distinct BR1 and BR3 BrTME and cell-intrinsic programs, we hypothesized that tumor cells skew microglia functional state and downstream interaction with T cells. To test this, we performed pairwise interactome analysis of tumor cell – microglia – T cells in our scRNAseq dataset to infer ligand-receptor (L-R) pairs implicated in their crosstalk (Figures 6A-C; S6A-B). We observed the highest predicted interaction number and strength coming from tumor cells to microglia in both models, with no significant differences in the number of interactions between BR1 and BR3 (Figures 6A, S6A). This crosstalk occurred among distinct microglia and tumor states and involved different types of interactions in BR1 compared to BR3 (Figures 6B-C). Key differences involved enhanced communication in BR1 from the tumor cell cluster enriched in immune and cell-cycle-MPs (T2) to reactive microglia states (Mg9, Mg12-13), including upregulation of the microglia-activating MIF_CD74_CD44 axis^73–75^ (Figures 6B-C). In BR1, we also observed enrichment of the LGALS9 axis, known to induce MHC-II expression on macrophages and DCs and strong anti-tumor responses in murine melanoma models^76,77^. BR3, on the other hand, showed stronger interactions than BR1 between tumor clusters displaying melanin-, development-, and neuro-MP activities (T3-T5) and homeostatic microglia (Mg1-2, Mg4, and Mg8; Figure 6B). Among the most upregulated interactions in BR3 were L-R pairs previously associated with microglia anti-inflammatory pathways (e.g., CD200_CD200R1, GAS6_MERTK), amyloid precursor response (e.g., APP_SORL1/TREM2), as well as those involved in maintenance of microglia homeostatic function, including phagocytosis (e.g., APOE_TREM2_TYROBP) and neuro-function (e.g., PTPRM-PTPRM) (Figure 6C). The predominant predicted tumor-to-microglia interactions in this model were driven by tumor junctional adhesion molecule (JAM) and extracellular matrix (ECM) ligands, such as collagens (e.g., COL4A2, COL6A3) and laminins (e.g., LAMA3, LAMC1; Figure 6C).

**Figure 6:**
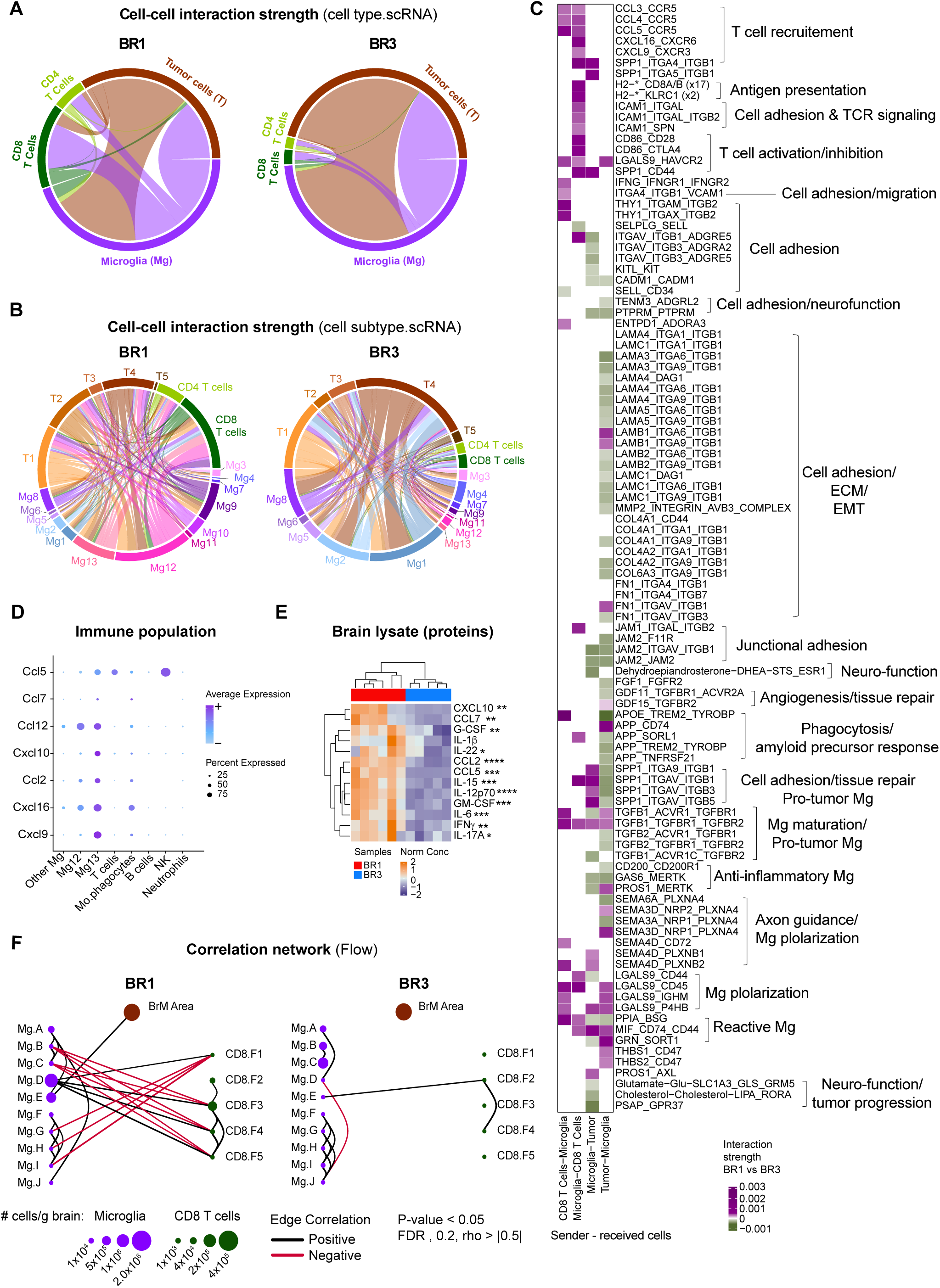
BrM tumor cells dictate the TME differences between ICB-responder BR1 and ICB-resistant BR3. (A-C) Cell-cell interaction analysis from scRNA-seq data of untreated BR1 and BR3 brains. (A-B) Circos plot showing the overall predicted interaction strength between indicated cell types (A) and indicated clusters within each cell type (B). The outside border color of the circos represent the sender and receiver cell populations. The width of the lines inside the circos plot indicates the interaction strength. The color of the lines inside the circos plot indicates the sender cell population. (C) Heatmap showing the top and bottom 10% most differential interaction (by strength) of ligand-receptor (L-R) pairs between models for indicated cell type pairs (see methods). Interactions stronger in BR1 versus BR3 are depicted in purple, interactions stronger in BR3 depicted in green. Numbers in parentheses indicate the number of different L-R pairs. No number indicates that only one L-R pair is present. (D) Dot plot showing the expression of selected chemokines in each immune cell population. (E) Cytokine/chemokine protein profile (normalized measurement) from untreated BR1 and BR3 whole brain lysates. (F) Correlation-based network analysis of total brain metastatic area (BrM area; determined by machine learning algorithm), microglia (Mg.A – Mg.J) and CD8^+^ T cell (CD8.F1 – CD8.F5) cluster cell count per gram of brain (quantified by spectral high-parametric flow cytometry analysis) from untreated BR1 and BR3 brains. Lines represent spearman correlations. A-D n = 2-3/group, E n = 5-6/group, K n = 8-10/group/experiment. *p <0.05, **p < 0.01, ***p < 0.001, ****p < 0.0001. See also Figure S6.

We then investigated the downstream microglia – T cell crosstalk. We found significantly stronger interactions from microglia to T cells, specifically CD8^+^ T cells, in BR1 compared to BR3 (Figures 6A, S6A). The reactive microglia states, which receive signals from BR1 tumor cells, were the ones predicted to preferentially interact with T cells in BR1 (Figure 6B). This crosstalk involved all the necessary signals to efficiently activate T cell responses, such as adhesion molecules, antigen presentation, TCR and co-stimulatory signaling (Figures 6C, S6B), explaining the T cell landscape observed in BR1. The inferred communications from microglia to CD8^+^ T cells were also characterized by T cell-attracting chemokines (Figure 6C)^78–80^, further suggesting that BR1 microglia are responsible for CD8^+^ T cell recruitment. We confirmed that these chemokine-encoding genes were predominantly expressed by BR1 microglia, with significantly higher protein levels found in BR1 compared to BR3 whole-brain lysates (Figures 6D-E, S6C). In marked contrast to BR1, the few microglia – T cell interactions predicted in BR3 involved only cell-cell adhesion processes between microglia homeostatic states (Mg1, Mg2, and Mg8) and naïve T cells (SELL) (Figures 6B-C, S6B).

To confirm these predicted interactions using an independent approach, we performed a correlation analysis of cell populations identified by flow cytometry. In agreement with the interactome analysis, we observed positive correlations between BR1-reactive microglia (Mg.D) and CD8^+^ T cells expressing activation markers (CD8.F3-F5) (Figures 6F). No correlations were observed between the BR3-associated microglia (Mg.A-C) and CD8^+^ T cell (Figures 6F). Altogether, these findings support our hypothesis that BR1 and BR3 tumor cell-intrinsic programs dictate microglia functional states, which in turn shape T cell phenotypes.

### The molecular features identified in our preclinical models predict T cell infiltration and BrM patient survival

To further validate the clinical relevance of our findings, we interrogated the potential associations of our ICB-responder and ICB-resistant signatures with BrTME composition and patient survival in publicly available datasets^13,21^. We first focused on the association between BR1 and BR3 MPs and CD8^+^ T cell infiltration in untreated melanoma patients^13^. To do so, we extracted CD8^+^ T cells from their scRNA-seq dataset and re-analyzed them using unsupervised clustering (Figure 7A). Consistent with our preclinical data, we found that the frequency of total CD8^+^ T cells within the biopsies positively correlated with BR1 immune-MP4 activity and negatively correlated with BR3 neuro-MP6 (Figure 7B). Out of the nine CD8^+^ T cell clusters identified in the BrM patients, eight shared similarities with the subsets found in our BrM models (clusters 0, 1, 3-8) (Figure 7A, S6D). All BR3 enriched MPs (MP2, MP6, MP7) showed a trend toward positive correlation with the CD8^+^ T cluster 5 resembling BR3 CD8^+^ T cells (Figures 7C, S6D). Conversely, CD8^+^ T cell cluster 8 resembling BR1 proliferative/cytotoxic CD8^+^ T cells showed a trend of positive correlation with BR1 MPs and negative with BR3 MPs (Figures 7C, S6D).

**Figure 7.**
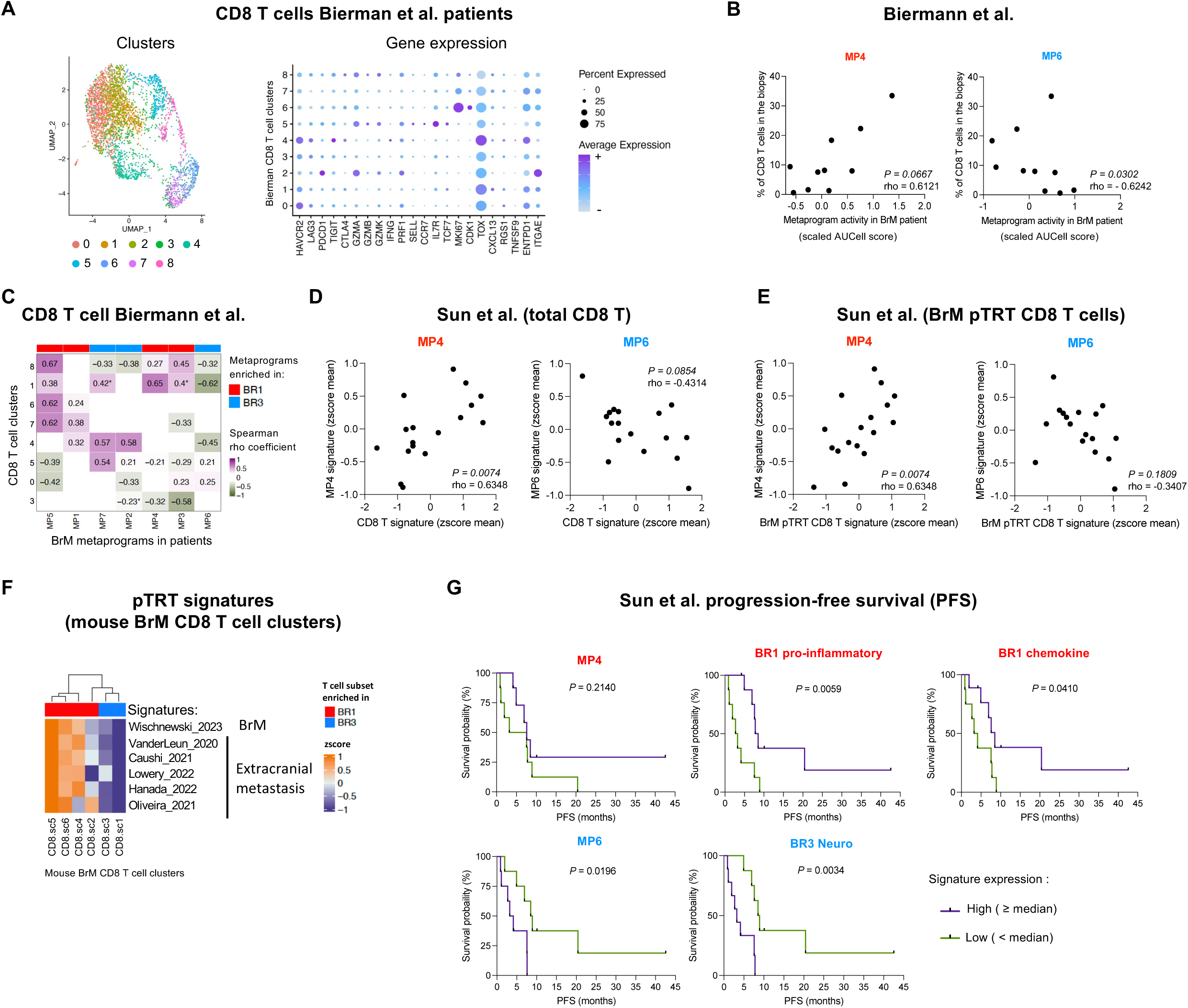
BR1 and BR3 signatures are associated with BrM patient CD8^+^ T cell infiltration and survival. (A-C) scRNA-seq analysis of CD8^+^ T cells from Biermann et al. 2022 untreated melanoma BrM patient biopsies. (A) UMAP projection and clustering (left). Dot plot showing the expression of selected differentially expressed T cell markers (right). (B) Spearman correlation of MP4 (BR1-MP) or MP6 (BR3-MP) activity and CD8^+^ T cell percentage among total cells per sample within each biopsy. (C) Heatmap representing spearman correlations between proportion (% of total cells) of individual CD8^+^ T cell clusters (row) and the indicated MP activity (column) within each biopsy. (D-E) Pseudobulk RNA-seq analysis of untreated and treated patient pan-cancer BrM biopsies from Sun et al. 2023. Spearman correlations of the indicated signatures’ expression (see methods). (F) Expression of “tumor-reactive” human CD8 T cell signatures defined by the indicated studies (row) within each mouse CD8^+^ T cell cluster from our study (column). Heatmap depicts Z-score expression across rows. (G) Progression-free survival of BrM patients stratified by median expression of the indicated signatures. BR1 and BR3 signatures are indicated in red and blue respectively. A-C n= 10 patients. D-E, G n=17 patients. F n = 2-3/group. See also Figure S6 and Tables S5-S6.

We corroborated our observations using pseudobulk RNA-seq analysis of data from a larger patient cohort of pan-cancer BrM, composed of individuals untreated or treated with various therapies, including ICB^21^. Total CD8^+^ T cell signature showed a significant positive correlation with BR1 immune-MP4 and a trend to negatively correlate with BR3 neuro-MP6 (Figures 7D, S6E; Table S6). Additionally, molecular signatures recently reported in patients as “potential tumor reactive T cell” (pTRT), both in BrM (BrM pTRT)^19^ and extracranial tumors^81–85^ showed a similar correlation pattern (Figures 7E, S6E; Table S6). These signatures were also expressed at a higher level in BR1 CD8^+^ T cells compared to BR3 (Figure 7F, S6F). Furthermore, we found a trend toward better progression-free survival (PFS) with high expression of the BR1 MP4 signature, while the BR3 neuro-MP6 was significantly associated with worse outcomes (Figure 7G, Table S6). We generated three additional signatures based on interactome and DEGs between BR1 and BR3: “BR1 pro-inflammatory”, “BR1 chemokines”, and “BR3 Neuro” (Table S6; see Methods). These signatures were confirmed to be more highly expressed in the corresponding groups using pseudobulk RNAseq analysis of our mouse BrM data set (Figure S6G). We found that the “BR1-proinflammatory” and “BR1 chemokine” signatures significantly correlated with better patient outcomes, whereas the “BR3 Neuro” signature was significantly associated with poorer PFS (Figure 7G). Taken together, our data demonstrate that the molecular signatures characteristic of our preclinical mouse models are also expressed in patient BrMs, correlate with their BrTME composition, and can predict patient outcomes.

## DISCUSSION

BrM remains a critical challenge in cancer treatment with most cases being resistant to ICB^3–5,7,86,87^. Unfortunately, the limited availability of patient biopsies and sparsity of immunocompetent preclinical models representing diverse ICB responses^23–25,88^ has hampered efforts to identify biomarkers and determinants of efficacy. Here, we have developed two translationally relevant melanoma BrM mouse models that overcome these limitations, enabling us to identify tumor molecular signatures that influence the BrTME and therapy responses. We show that ICB-responder BrMs (BR1) express a molecular program of response to IFN, antigen presentation, and inflammatory pathways that polarizes microglia into reactive states. These microglia are characterized by high expression of MHC-I/II, co-stimulatory molecules, and T cell-recruiting chemokines, which enhance T cell recruitment and activation, favoring ICB efficacy. In contrast, resistant BrMs (BR3) express programs involved in neurological function and evade the immune response by preserving a homeostatic state within the BrTME. Importantly, we validated the translational relevance of our findings by demonstrating that BR1 signatures positively correlated with T cell infiltration in human BrMs, and were associated with improved patient outcomes, whereas BR3 signatures showed negative correlation with T cells and were associated with poorer patient outcomes.

We found striking differences between responder and resistant models in the BrTME T cell subsets. The presence of a proliferative T cell subset in the responder model suggests that local expansion may contribute to shaping T cell landscape in the brain. An interesting finding from our study is that therapy induces an increase in T cell infiltration in both responder and non-responder models; however, the phenotypic characteristics of these cells were markedly distinct. These observations can help explain contradictory results observed from studies showing a positive correlation between T cell infiltration and improved patient outcomes vs. others that did not^21,89–92^, stressing the importance of studying other immune populations that may influence T cell function. Indeed, our findings reveal that not only T cells but also microglia distinguish responders from non-responders. We show that specific reactive microglia states are significantly enriched in ICB responders and positively correlate with CD8^+^ T cell infiltration. Although the relationship between microglia – T cell crosstalk and therapy responses has not been explored in patients, recent studies have reported positive correlations between MHC-II expression in microglia and CD8^+^ T cells in BrM patients^19,89^. Supporting our results, previous data have shown that microglia sharing some of the BR1 signatures can attract or activate T cells in other contexts, such as CNS infection^93,94^, Alzheimer’s disease^95^, or glioblastoma (GBM) treated with ICB^96^. Of note, CXCL9 alone or MHC-II^+^ reactive microglia were recently found to be required for GBM response to PD-1 or CTLA-4 blockade, respectively^96,97^.

Recent studies using breast and lung BrM preclinical models have shown that microglia with some BR1-reactive characteristics could suppress T cell activity and support BrM development, while others have demonstrated that microglia can recruit T cells and impair BrM progression^54,56,98,99^. These dual microglia functions are not mutually exclusive and suggest a context-dependent role. We observed that BR1 microglia states express different T cell immune checkpoints, which can paradoxically both induce and limit T cell activation in the absence of treatment. When these inhibitor axes are targeted by ICB, they unleash the T cells’ anti-tumor potential. Thus, our findings indicate that microglia can play context-dependent roles in BrM based on the brain tumor subtype and the therapy utilized. Moreover, our observations can help explain the inconsistent pro- and anti-tumor impact of microglia depletion observed across independent preclinical studies^10,54,56,98^. While depleting microglia has been suggested as a potential therapeutic approach for brain tumors, our data indicate that such a strategy could be more harmful than beneficial depending on the context. Instead, modulating the balance between pro- and anti-tumor microglia states might be a more effective alternative. Here we reveal four major types of interactions used preferentially by the ICB-resistant BR3 tumors to shape microglia states: (1) ligands known to inhibit microglia reactivity such as CD200 and Gas6^100–103^, (2) response to amyloid precursor, (3) components of the extra-cellular matrix such as collagens and laminins, and (4) ligands implicated in neurological processes. In the CNS, these axes mediate crosstalk involving microglia and other cells (e.g., neurons, astrocytes, and endothelial cells) and are crucial to maintaining brain homeostasis and resolve neuroinflammation^100,104–112^. Others have described the pro-tumoral effect of CD200-CD200R and Gas6-MertK axes in primary brain tumors^102^, and amyloid beta in melanoma BrM^113^. A recent report implicated collagens in murine breast and lung BrM growth^98^. The latter study demonstrated that tumor-derived collagen polarizes pro-tumor microglia through TNFα-NF-κB signaling, suppressing T cell infiltration and activation^98^. We found this pathway differentially upregulated in BR3 microglia, suggesting that similar interactions might contribute to the BR3 BrTME profile and its ICB-resistance. Additionally, while no direct link has been established between neurotransmitters and BrM response to therapy, independent studies have shown that glutamate can support GBM growth^114,115^ and decrease microglia reactive states^108^. Thus, our findings suggest that the ICB-resistant BR3 BrM model hijacks natural CNS tolerance systems to support homeostatic microglia, to inhibit reactive states, and to promote pro-tumoral polarization. The molecular axes that we identified not only deepen our understanding of how BrM tumors shape the BrTME but also provide potential targets for overcoming therapy resistance^114,115^.

We further show that BrM tumor cells with proliferative and inflammatory signatures (BR1) benefit from ICB therapies, whereas BrM tumor cells enriched in neurological features, such as axon guidance (BR3), do not and may require alternative or additional strategies, such as targeting myeloid cells or tumor ligands. We extended our findings to a pan-cancer BrM cohort, confirming that BR1 and BR3 signatures correlate with CD8^+^ T cell infiltration and patient outcomes. These observations provide potential predictive biomarkers and valuable insights for treatment decision-making.

An important hurdle to improving BrM therapy is the difficulty in accessing patient samples, underscoring the importance of relevant preclinical models to elucidate mechanisms of resistance. Through the generation of new models that recapitulate the complexity and clinical behavior of human BrMs, we identified key molecular features associated with ICB response. Therefore, the BR1 and BR3 models constitute a valuable resource to perform mechanistic studies. Our comparative and integrative analyses of mouse and human data have provided critical insights into the mechanisms governing treatment response, contributing toward the development of more effective therapeutic options for BrM patients.

### Limitations of the study

Here we focused on the immune and the tumor cell compartments, but we cannot rule out the role of other important BrTME actors, such as astrocytes and endothelial cells. Further studies should investigate how these cell types can influence ICB responses. An important limitation in the field, that persists in our study, is the dearth of available human BrM samples, which limits the possibility to directly associate ICB response status and different BrM features in patients. Although we demonstrated that BR1 and BR3 signatures are associated with the patients’ BrTME and survival in cohorts that included ICB treatment, it will be important to test our findings when larger cohorts of treated patients become available. A remaining unmet need is the ability to identify valuable predictors of response employing less invasive approaches, such as monitoring blood or cerebrospinal fluid. Despite these constraints, our report significantly enhances our understanding of the mechanisms by which BrM tumor cells can impact BrTME and ICB responses. We anticipate that our findings will facilitate the development of new therapeutic approaches and help inform clinical trials for melanoma patients with brain metastases.

## ACKNOWLEDGMENTS

We thank Patricia Steeg’s laboratory, particularly Alex Wu, for their advice and help with the BrM model development. We are grateful to the following NCI facilities: CCR Single Cell Analysis Facility, SpiTR Nanoscale Protein Analysis Section, LGI Flow Cytometry Core, CCR Genomics Technology Laboratory, NIH Animal Facility Bethesda and Frederick, and NCI LASP Small Animal Imaging Program (SAIP). We also thank Jennifer Dwyer (LCBG) and Katerina Grafanaki (University of Patras, School of Medicine) for technical help, Christophe Cataisson (LCBG) and Grégoire Atlan-Bonnet (LICI) for access to equipment, and members of LICI and LCBG for their helpful discussion. This research was supported in full by the Intramural Research Program of the NIH (CCR-NCI), including CCR FLEX Synergy Award, and a CCR Excellence in Postdoctoral Research Transition Award (2020) to EP-G. GK was supported by a Fulbright Foundation Visiting Research Fellowship from Greece (2023–2024). EP-G was supported in part by the Spanish *Agencia Estatal de Investigación* (RYC2021-034893-I and PID2022-141113OA-I00) and a Young Investigator Award from Melanoma Research Alliance (Ref: #1037420). This work utilized the computational resources of the NIH HPC Biowulf cluster (https://hpc.nih.gov).

## INCLUSION AND DIVERSITY

One or more of the authors of this paper self-identifies as an underrepresented ethnic minority in science. One or more of the authors of this paper self-identifies as a member of the LGBTQ+ community

---

**Figure S1.**
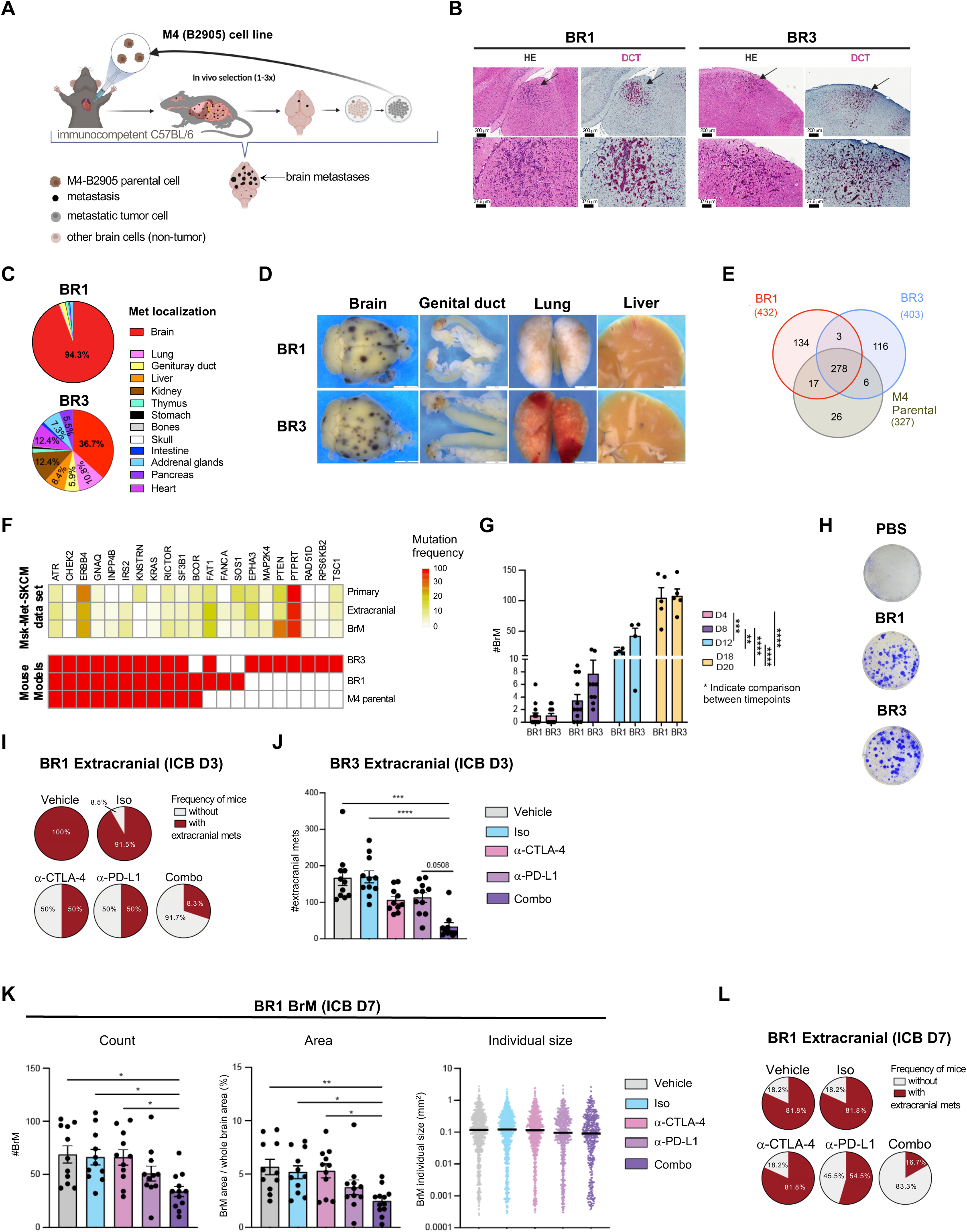
Characterization of the metastatic potential, mutational landscapes, and ICB responses of BR1 and BR3 models, related to Figure 1. (A) Experimental design for the development of BrM melanoma models. (B) Hematoxylin / eosin (HE) and dopachrome tautomerase (DCT; pink) immunostaining images of brain sagittal sections showing BR1 (left panels) and BR3 (right panels) BrMs. Magnification 15x (top) and 40x (bottom). (C) Percentage of metastasis per organ in BR1 (top) and BR3 (bottom) models. (D) Representative stereomicroscope images of the indicated organs from BR1 and BR3 BrM-bearing animals. (E) Venn diagram showing the number of genes mutated and their overlap between the indicated cell lines. (F) Mutation frequency of indicated genes found in the brain-metastatic cell lines (marked in red in bottom heatmap), BrMs, extracranial metastases, and primary tumors from patients with BrMs from the MSK MetTropism cutaneous melanoma (SKCM) dataset. Color scale represents the mutation frequency in the indicated sample group. (G) BrM counts at the indicated time point after intracardiac injection (i.c.) in BR1 and BR3 bearing animals. D18/D20 indicate harvest at experimental endpoint. Asterix indicated difference between indicated timepoints. (H) Macroscopic pictures of cultured cell colonies recovered from mouse brains from BR1 and BR3 bearing animals or non-BrM control (PBS). (I) Proportion of mice with or without BR1 extracranial metastases after treatment on day 3 post tumor injection with PBS (vehicle), isotype control (Iso), anti-CTLA-4 (α-CTLA-4) or anti-PD-L1 (α-PD-L1) alone, or anti-PD-L1 + anti-CTLA-4 combination therapy (Combo). (J) BR3 extracranial metastatic count per mouse in animals treated as in I. (K) BR1 BrM count (left), total brain metastatic area per individual brain (middle), and size of individual BrM lesion (right) from animals treated as in I starting treatment on day 7 post tumor injection. (L) Proportion of animals from K with or without extracranial metastases. B one representative of 2 experiments, n=2-5 mouse/group/experiment. C data from 3 experiments combined, n=2-5 mouse/group/experiment. D one representative of 3 independent experiments, n=2-5 mouse/group/experiment. G data combined from 2 independent experiments, n = 2-5 mouse/group/experiment. I-L representative of 2 independent experiments per model, n = 9-12/group/experiment. G, J, K data shown as mean ± SEM *p <0.05, **p <0.01, ***p < 0.001, ****p < 0.0001. See also Table S1

**Figure S2.**
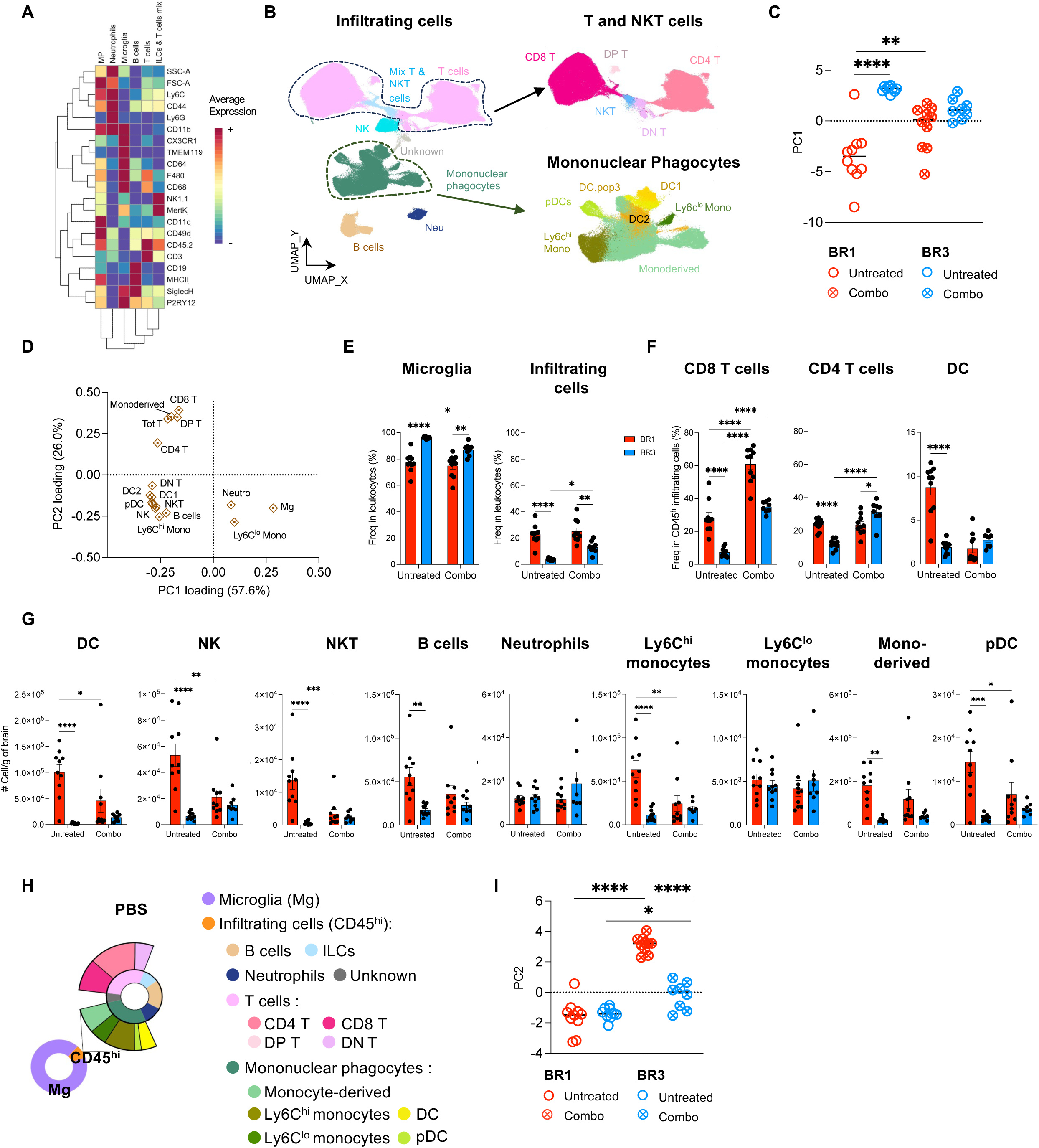
Global immune BrTME characterization of untreated and treated BR1 and BR3 bearing brains, related to Figure 2. (A) Heatmap representing expression level of indicated surface markers (scaled by marker) for each immune cluster with hierarchical clustering indicated on left and markers on top. (B) UMAP projection of indicated immune cells. NKT: natural killer T; T cell subsets: CD8^+^CD4^-^ (CD8 T), CD8^-^CD4^+^ (CD4 T), CD8^-^CD4^-^ (DN T), CD8^+^CD4^+^ (DP T); mononuclear phagocytes including dendritic cells [DC1, DC2, DC.pop3, plasmacytoid DC (pDCs)] and monocytes subsets [Ly6C^hi^ mono and Ly6C^lo^ mono]. (C) Quantification of principal component 1 (PC1) from Figure 2D. (D) PC1 and PC2 loadings from Figure 2D. (E-F) Proportion of indicated immune cells. (G) Absolute number of indicated immune cells per gram of brain. (H) Proportion of microglia (Mg; purple) and infiltrating cells (CD45^hi^; orange) within total leukocytes in control (PBS) brains. Proportion of indicated populations in total infiltrating cells (inset). (I) PC2 quantification from Figure 2D. C, E-G, and I data shown as mean, n = 8-10/group/experiment. *p <0.05, **p < 0.01, ****p < 0.0001. See also Table S2.

**Figure S3.**
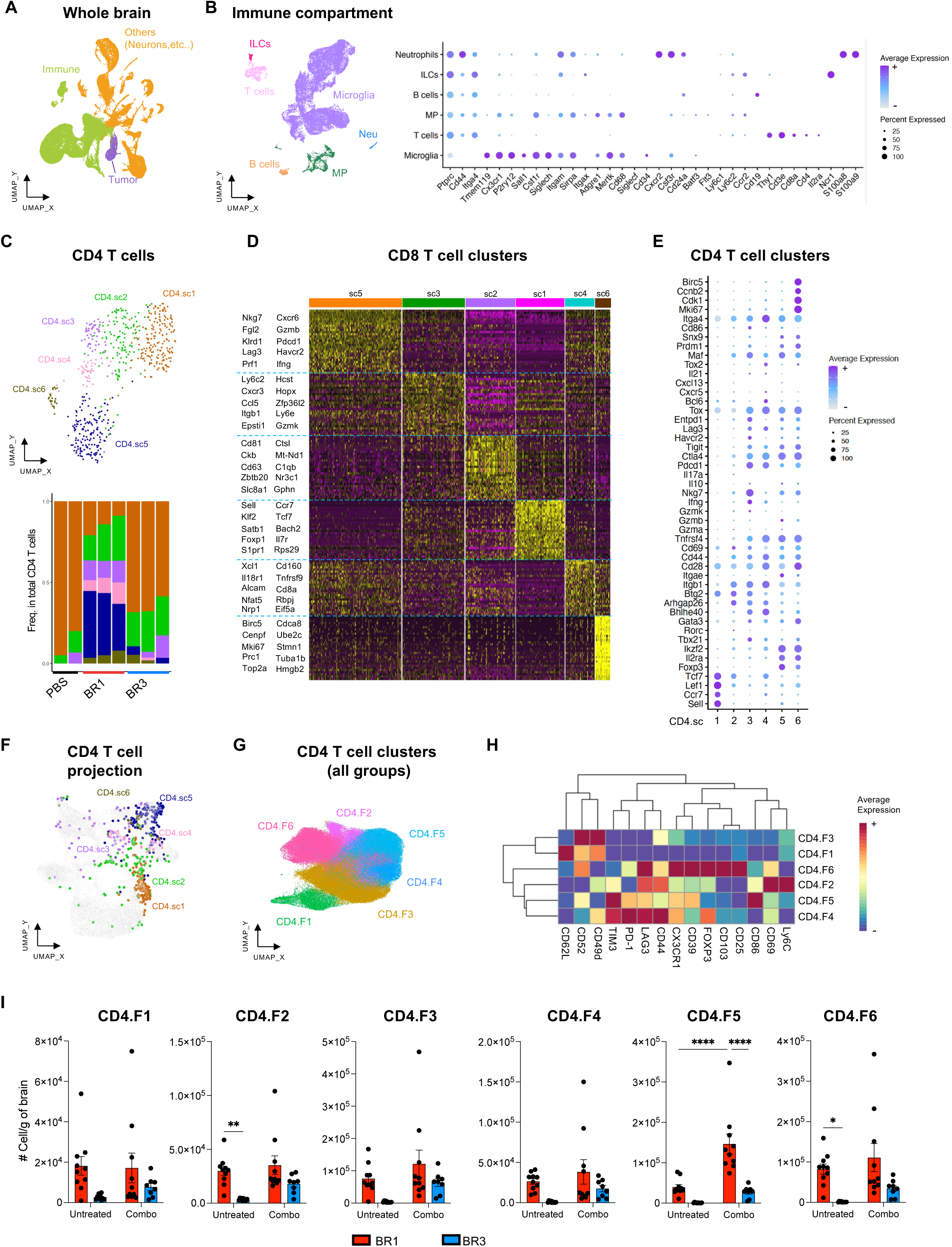
Distinct T cell characteristics found in the ICB-responder model, related to Figure 3. (A-B) ScRNA-seq analysis of untreated BR1, BR3, and PBS whole brains. (A) UMAP projection of all cells. (B) UMAP projection of immune cells (left). Dot plot showing the expression of selected genes among clusters (right). (C-F) scRNA-seq analysis of indicated brain T cell populations. (C) UMAP projection of indicated CD4^+^ T cell clusters among total CD4^+^ T cells (top). Proportion of indicated CD4^+^ T cell clusters in total CD4^+^ T cells per sample (bottom). (D) Heatmap representing expression level of the top 30 CD8^+^ T cell cluster-specific genes. (E) Dot plot showing the expression of selected genes among CD4^+^ T cell clusters. (F) Projection of our CD4^+^ T cell clusters onto the single-cell reference atlas for T cell states ProjecTILs. See figure 4E for the Reference map (left). (G-I) High-parametric flow cytometry analysis of CD4^+^ T cells from BR1 and BR3 brains untreated or combo treated. (G) UMAP projection of CD4^+^ T cell populations. (H) Heatmap representing expression level of indicated surface markers (scaled by marker) for each CD4^+^ T cell cluster with hierarchical clustering indicated on left and markers on top. (I) Absolute number of indicated CD4^+^ T cell clusters per gram of brain. A-D n = 2-3/group/experiment. F-G Data shown as mean ± SEM, n = 8-10/group/experiment. *p <0.05, ***p < 0.001, ****p < 0.0001. See also Table S3-S4.

**Figure S4.**
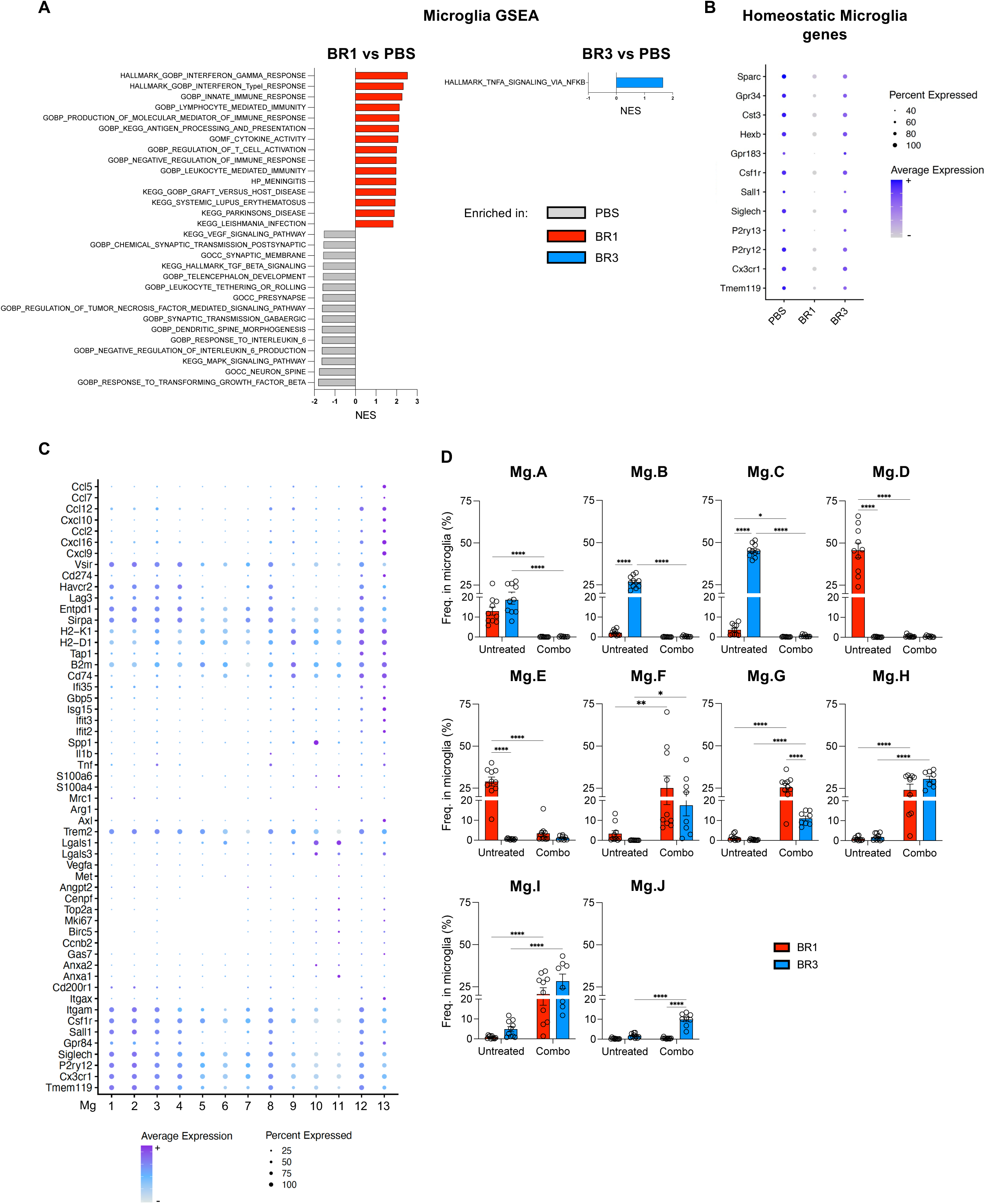
Distinct microglia are found in ICB-responder and resistant BrMs, related to Figure 4. (A-C) scRNA-seq analysis of microglia from untreated BR1, BR3, and PBS brains. (A) GSEA showing HP, GO, and KEGG pathways selected among the top 10 enriched in total BR1 microglia versus PBS (left) and in total BR3 microglia versus PBS (right) (see Methods). (B) Dot plot showing the expression of homeostatic microglia genes per group. (C) Dot plot showing expression of selected genes among microglia clusters. (D) High-parametric spectral flow cytometry analysis of microglia from BR1 and BR3 bearing animals untreated or combo-treated. Proportion of indicated microglia clusters in total microglia. A-C n = 2-3/group/experiment. D n = 8-10/group/experiment. *p <0.05, **p < 0.01, ***p < 0.001, ****p < 0.0001

**Figure S5.**
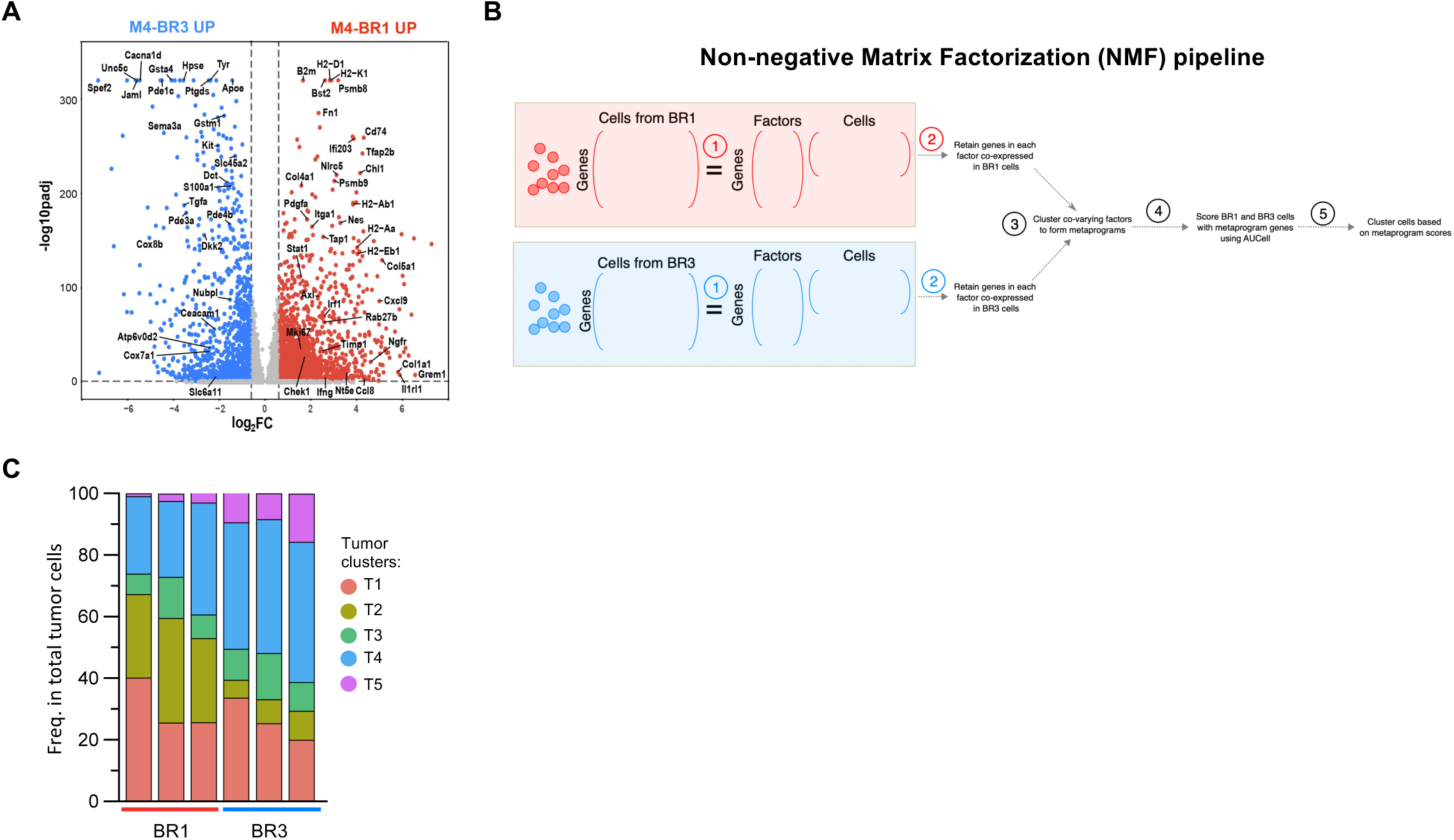
BR1 and BR3 tumor cell molecular characterization by scRNA-seq, related to Figure 5. scRNA-seq analysis of tumor cells from untreated BR1 and BR3 brains. (A) Volcano plot of BR1 versus BR3 differentially expressed genes. (B) Schematic of metaprogram discovery and their use in clustering malignant cells in BR1 and BR3 models. Step 1: NMF is carried out (k = 30 factors) separately in malignant cells from BR1 and BR3 models based on genes that are expressed in at least 1% of cells in either model. Step 2: Genes with a high weight in each factor are then filtered to retain genes that are co-expressed. Step 3: The activity of each filtered factor is computed across all malignant cells using AUCell. A correlation matrix between all pairs of factors is computed based on these activities, and after hierarchical clustering, correlated factors are grouped together. Genes occurring in at least 25% of constituent factors in a group are retained to form metaprograms. Step 4: AUCell is used to compute the activity of all metaprograms across all malignant cells. Step 5: Cluster cells using Louvain clustering based on metaprogram activity scores. (C) Proportion per sample of indicated tumor clusters. A-C n = 2-3/group. See also Table S5.

**Figure S6:**
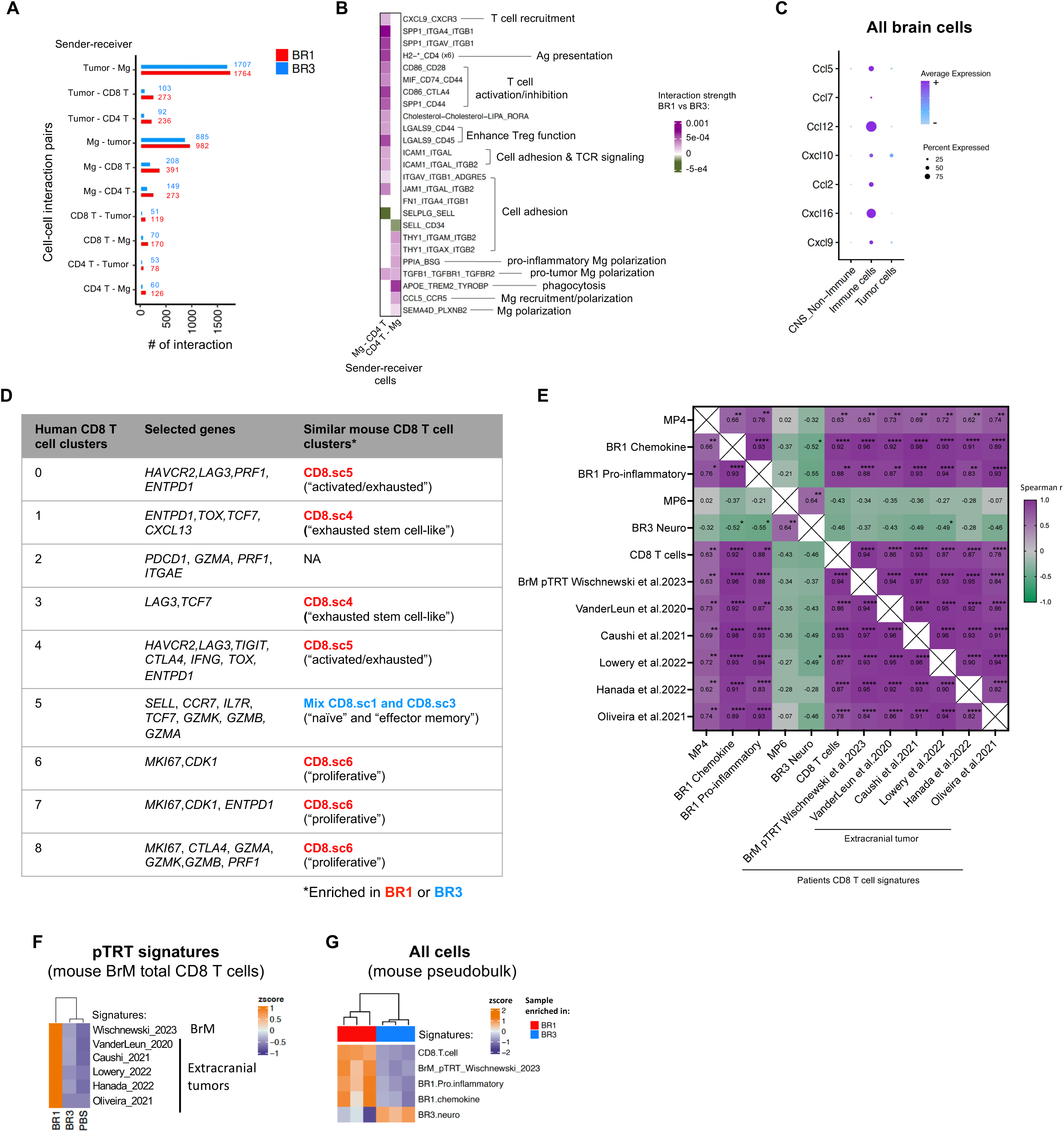
BR1 and BR3 tumor molecular features control the BrTME. (A-B) Cell-cell interaction analysis from scRNA-seq data of untreated BR1 and BR3 brains. (A) Total number of interactions between indicated cell sender-cell receiver pairs per group. (B) Heatmap showing the top and bottom 10% most differential interaction strength of ligand-receptor (L-R) pairs between BR1 versus BR3 (see methods) for each indicated cell type pair. Interactions stronger in BR1 versus BR3 are depicted in purple, interactions stronger in BR3 depicted in green. Numbers in parentheses indicate the number of different L-R pairs. No number indicates that only one L-R pair is present. (C) scRNA-seq analysis of all brain cells from untreated BR1 and BR3-bearing mice. Dot plot showing the expression of selected chemokines in each cell type. (D) Human CD8^+^ T cell clusters identified in Biermann et al. 2022 dataset with their selected expressed genes and similar mouse CD8^+^ T cell clusters from BR1 and BR3 models. (E) Pseudobulk RNA-seq analysis of Sun et al. 2023 pan-cancer BrM patient cohort. Spearman correlation matrix between the different BR1 and BR3 signatures in patient biopsies. See also Table S6. (F) scRNA-seq analysis of total CD8^+^ T cells from BR1, BR3, and non-tumor (PBS) bearing mice. Heatmap showing Z-score expression level of various published gene signatures described as “tumor-reactive” CD8 T cells in cancer patients. See supplementary table S8 for details. (G) Pseudobulk RNA-seq analysis of BR1 and BR3-bearing brains. Heatmap showing Z-score expression of CD8^+^ T cells, BR1, and BR3 signatures among total brain cells across samples. A-C, F-G n = 2-3/group. D n= 10 patients. E n=17 patients. *p <0.05, **p < 0.01, ***p < 0.001, ****p < 0.0001. See also Table S6.

## RESOURCE AVAILABILITY

### Lead contact

Further information and requests for resources and reagents should be directed to and will be fulfilled by the lead contact, Romina S. Goldszmid.

### Materials availability

Cell lines generated in this study will be made available from the lead contact (R.S.G) and/or G.M, C-P. D., E.P.-G. upon request but may require a completed NIH Materials Transfer Agreement.

### Data and code availability

Single-cell RNA-seq and whole exosome sequencing data generated by this study will be deposited at GEO and made publicly available as of the date of publication. This paper also analyzes existing, publicly available data. All accession numbers will be listed in the key resources table and method section. Any additional information required to reanalyze the data reported in this paper is available from the lead contact upon request.

## EXPERIMENTAL MODEL AND STUDY PARTICIPANT DETAILS

### Animals

All animal studies were approved by the Institutional Animal Care and Use Committee of National Cancer Institute and were conducted in accordance with the IACUC guidelines at the National Institutes of Health Guide for the Care and Use of Laboratory Animals. C57BL6/NCr female mice were obtained from Charles River Laboratories (CRL) and were group-housed with up to 5 mice per cage, in a specific pathogen-free environment in the Center for Cancer Research (CCR) animal facility at the NCI Bethesda or the NCI Frederick. In both facilities, mice were housed under standard conditions with *ad libitum* access to food and water and under a 12/12h light/dark cycle. In all studies, 6-8 weeks-old mice were used. For experiments in which brain metastatic untreated mice were compared, the mice distribution per group was randomized to obtain similar weight per group, as weight can impact the success of the tumor implantation by intracardiac injection. Additional randomization was performed for brain metastatic therapy studies after tumor implantation but before the administration of the immunotherapy, considering the order of injection, as this factor might also influence tumor implantation.

### Cell lines

M4-BR1 (BR1) and M4-BR3 (BR3) brain metastatic cell lines were generated by intracardiac (i.c.) injection of 5×10^5^ M4-B2905 (M4) cells, which originated in a hepatocyte growth factor (Hgf^tg^) transgenic C57BL/6J male mouse as described before^27^. Successive *in-vivo* selection for homing to the brain was done for one (BR1) or three (BR3) cycles. Briefly, brains from 5-10 mice i.c. implanted were cut separately in small pieces and homogenized in 0.25% trypsin (Gibco #25200056) for less than 10 min. Whole brain homogenates were plated and maintained in culture for 2 weeks or until expanded in enough cell numbers. Successful cultures were harvested and injected i.c. again to repeat the cycle.

M4 cell line was cultured in RPMI medium supplemented with 10% FBS and 1% l-Glutamine (200 mM); BR1 and BR3 melanoma cell lines were cultured in high glucose, GlutaMAX^TM^ supplement, HEPES DMEM medium (Gibco #10564011) supplemented with 10% heat inactivated FBS at 37°C, 5% CO_2_ in air atmosphere. All cells were tested and found negative for specific mouse pathogens (MTBM test) and were confirmed as Mycoplasma negative using a Mycoplasma detection kit (Lonza, LT07-418).

### Publicly available patient dataset

The single-cell read-count matrix of the single-cell RNA-sequencing (scRNA-seq) dataset from BrM patient biopsies used in this manuscript were downloaded from Gene Expression Omnibus (GEO): Smalley et al. 2021 (GSE174401) 14 melanoma patients untreated or treated with immune-checkpoint blockade (ICB) and/or targeted therapy, and/or radiotherapy (RT); Sun et al. 2023 (GSE193745), 17 pan-cancer patients untreated or treated with ICB and/or targeted therapies and/or RT. Alvarez-Breckenridge et al. 2023, was downloaded from Broad Single Cell Portal (SCP1493) and composed of 27 melanoma patients untreated or treated with ICB and/or targeted therapies and/or RT. Biermann et al. 2022 (GSE200218), single-nucleus RNA-seq from 25 untreated melanoma patients (15 BrM biopsies and 10 extra-BrM biospies) were downloaded from GEO.

### Experimental brain metastasis

C57BL/6J mice were shaved at the chest and intracardially (i.c.) injected with 5×10^5^ BR1 cells or 7.5×10^4^ BR3 cells per mouse. Animals were kept under anesthesia (2-3% isoflurane in oxygen) and continuous monitoring during the whole procedure. The endpoint of the model was set when the animal weight loss reached 5% of the initial weight and/or any behavior changes were observed such as shortness of breath or slow motion, and was reached around 18-20 days after i.c. injection. When, in some rare cases, animals did not show any brain metastasis implantation at endpoint, they were excluded from the study.

### Immune checkpoint inhibitor treatment

To test and compare ICB monotherapy and combination efficacy between the models we implanted BR1 and BR3 cells by ultrasound-guided intracardiac injection. Mice were treated with monotherapy anti-CTLA-4 (BioXcell, BP0164) and anti-gp120 IgG1 isotype control (BioXcell, BE0154), or anti-PD-L1 (Genentech, clone 6E11) and anti-IgG2b isotype control (BioXcell, BP0086), or anti-CTLA-4 and anti-PD-L1 combination therapy, or a mix of anti-gp120 IgG1 and anti-IgG2b isotype control, or with PBS as a supplementary control. Anti-CTLA-4 and anti-IgG2b antibodies were administered twice per week for a total of 4 doses. The first dose was administered intravenously (IV) at 200mg per mouse, and the following 3 doses were administered intraperitoneally (IP) at 200mg per mouse. Anti-PD-L1 and anti-gp120 IgG1 antibodies were administered twice per week for a total of 5 doses. The first dose was administered intravenously (IV) at 200mg per mouse, and the following 4 doses were administered intraperitoneally (IP) at 100mg per mouse. Depending on the studies this treatment regimen started 3 days or 7 days post-tumor implantation.

### Tissue collection

Mice were euthanized in CO_2_ chambers following the American Veterinary Medical Association (AVMA) guidelines at endpoint or as indicated for kinetic studies (4, 8 or 12 days after ic. injection). Mice were then intracardially perfused with 20 mL of ice-cold 1X PBS supplemented with 2 mM EDTA. Lungs, thymus, livers, kidneys, adrenal glands, digestive and reproductive ducts were harvested, fixed for 24h in 10% formalin at room temperature, and then kept in ethanol 70% at room temperature until metastasis quantification. Brains were harvested, immediately imaged using a stereomicroscope, and prepared for immunohistochemistry, immunophenotyping, or single-cell RNA-sequencing (scRNA-seq) experiments.

### Microscopy imaging

Macroscopic whole-brain pictures were acquired using a Leica M165 FC stereomicroscope (LEICA, USA) equipped with a Leica LED5000 ring light illumination (RL-80/40) and a digital color FLEXACAM C1 camera, as previously described^30^. Briefly, two pictures per brain were acquired: the whole ventral surface and the whole dorsal surface, at 4X magnification using the camera set on brightfield. Pictures focusing on individual brain metastases were acquired using the same microscopic platform and settings but at 12.5X magnification. All images were acquired with the Leica Application Suite X (LAS X) software, version 5.1.0.

### Metastasis burden quantification

Extracranial metastasis burden was assessed by manual counting on fresh organs using a SMZ745T Nikon (Nikon, USA) stereomicroscope trinocular system at 2-4X magnification range. Brain metastasis burden was assessed using a machine learning algorithm that we developed to detect, count and measure size of each metastatic lesions and total metastatic area on whole brain LEICA stereomicroscope pictures, as previously described^30^ using the software QuPath^116^.

### Colony formation assay

BR1, BR3-BrM bearing, and PBS (non-tumor bearing controls) brains were harvested at 4 days post-i.c. injection (as described above) and were processed using Adult Brain Dissociation Kit (Miltenyi, #130-107-677) following the manufacturer’s instruction. Each single cell suspension obtained were seeded in duplicate in 6-well plates and cultured in high glucose, GlutaMAX^TM^ supplement, HEPES DMEM medium (Gibco #10564011) supplemented with 10% heat inactivated FBS and 50 mg/mL of gentamicin, at 37°C, 5% CO_2_ in air atmosphere. Media was renewed and dead cell remove every 2 days. After 10 days, the cells were washed with 1X PBS, fixed for 5 min at room temperature in ice cold methanol (VWR Chemicals, 85800.290P), washed with 1X PBS and then stained with a 2.3% crystal violet solution (Sigma-Aldrich, HT90132). After removing the excess stain, the cells were washed in molecular biology water (Vita scientific, 351-029-101) and the well were air-dried overnight at room temperature and protected from light. The wells were then imaged using a Nikon D90 camera (Nikon, USA).

### Histology and immunohistochemistry

Formalin-fixed and paraffin-embedded 5μm sagittal or longitudinal sections of brains from BR1 or BR3 BrM-bearing mice or injected with PBS as non-BrM control were stained with hematoxylin and eosin (HE) or DCT (Pep8h58^117^) antibody using standard immunohistochemistry methods. Antigen retrieval was performed at 95°C for 20 minutes in a pressure cooker using target retrieval buffer pH6 (Dako, S1699). Protein Blocking reagent (Dako, X0909) and Bloxall blocking solution (Vector, SP-6000-100) were used to block nonspecific proteins and endogenous peroxidase and alkaline phosphatase activities following provider recommendations. DCT antibody (dilution 1:5000) was incubated overnight at 4°C in a humidity chamber. Antibody detection was performed using Impress AP Reagent anti-rabbit Ig (Vector, MP-5401) and ImPACT Vector Red (Vector, SK-5105) following provider protocol. Slides were counterstained with Mayer’s Hematoxylin (Electron Microscopy Sciences, 26043-06) and GIEMSA (New Comer Supply, 1120A) for 2-5 minutes each at. Whole slides were scanned using an AT2 Leica scanner (Leica, USA) at 20X magnification and digital image files were visualized using HALO link (Indica labs).

### Whole exome sequencing of brain metastatic cell lines

Mouse genomic DNA concentration was measured using Quant-iT PicoGreen dsDNA Assay Kit (Thermofisher scientific, P11496). 200ng of DNA was sheared by Covaris Instrument (E210) to ∼150-200bp fragments. Shearing was done in a Covaris snap cap tube (microTUBE Covaris p/n 520045) with the following parameters: duty cycle, 10%; intensity, 5; cycle burst, 200; time, 360 seconds at 4°C. Samples quality assessment/size was validated by Bioanalyzer DNA-High sensitivity kit (Agilent # 5067-4626).

Agilent SureSelectXT library prep ILM Kit (Agilent Technologies, #G9611A) was used to prepare the library for each sheared mouse DNA sample. DNA fragment’s ends were repaired, followed by Adenylation of the 3’ end and then ligation of paired-end adaptor. Adaptor-ligated libraries were then amplified (Pre-capture PCR amplification: 98°C 2 minutes, 10 cycles; 98°C 30 seconds; 65°C 30 seconds; 72°C 1 minute; then 72°C 10 minutes, 4°C hold) by Herculase II fusion enzyme (Agilent Technologies, #600679). After each step DNA was purified by Ampure XP beads (Beckmann Coulter Genomics #A63882). DNA Lo Bind Tubes, 1.5mL PCR clean (Eppendorf # 022431021) or 96 well plates were used to process the samples.

Samples were analyzed by bioanalyzer using DNA-1000 kit (Agilent #5067-1504). Concentration of each library was determined by integrating under the peak of approximately 225-275bp. Then each gDNA library (∼750-1000ng) was hybridized with biotinylated mouse specific capture RNA baits (SureSelectXT Mouse All Exon, catalog #5190-4641, 16 reactions) in the presence of blocking oligonucleotides. Hybridization was done at 65°C for 16 hours using SureSelectXT kit reagents. Bait-target hybrids were captured by streptavidin-coated magnetic beads (Dynabeads MyOne Streptavidin T1, Life Technologies, #6560) for 30 minutes at room temperature. Then after a series of washes to remove the non-specifically bound DNA, the captured library was eluted in nuclease free water and half volume was amplified with individual index (Post-capture PCR amplification: 98°C 2 minutes; 10 cycles of 98°C 30 seconds; 57°C 30 seconds; 72°C 1 minute; 72°C 10 minutes, 4°C hold). The Bioanalyzer High sensitivity kit was used to validate the size of the libraries prior to sequencing. The libraries were sequenced in a NovaSeq S2-300 chip.

### Single nucleotide variant (SNV) analysis

Fastq sequence reads were mapped to the mouse reference genome mm10 (https://www.ncbi.nlm.nih.gov/datasets/genome/GCF_000001635.20) with BWA or Bowtie. Single nucleotide variants (SNV) were identified using samtools mpileup or GATK HaplotypeCaller. Mouse germline single nucleotide polymorphisms (SNPs) were filtered out using the Sanger Mouse Genomes Project database for variants identified from whole genome sequencing of 36 mouse strains (ftp://ftp-mouse.sanger.ac.uk/current_snps/mgp.v5.merged.snps_all.dbSNP142.vcf.gz). Variants with a Phred-scaled quality score of <30 were removed. Variants that are present in normal spleen samples (in-house collection) were also removed. Variants were annotated with Annovar software to identify non-synonymous mutations. All analyses were carried out on NIH biowulf2 high performance computing environment. Statistical analysis was carried out in R environment.

For pathway analysis of the genes mutated in each cell line we used Ingenuity Pathway Analysis (QIAGEN Digital Insights). For comparison with human BrM we selected 21 genes involved in MAPK and PI3K/mTOR pathways that were mutated in at least one of the BR cell lines and retrieved mutation data from cBioportal (https://www.cbioportal.org/). We used MSK MetTropism data set^31^ selecting the samples from metastatic cutaneous melanoma patients and sorted the data in three groups: 1) brain metastases (BrM), 2) all other extracranial metastases, and 3) primary melanomas from patients with BrM. The heatmaps were generated using R package Ggplot2.

### Flow cytometry on leukocytes

For immunophenotyping experiments, freshly harvested brains were processed to obtain single-cell suspension following a previously described protocol^118^ with some adaptations. Briefly, whole brain tissues were mechanically disrupted using a dounce homogenizer in plain RPMI media. Brain leukocytes were enriched by a 70%/30% Percoll gradient centrifugation and then washed in 10 mL of 1X PBS. After centrifugation. Single cell suspensions were resuspended in 1X PBS and stained with LIVE/DEAD^TM^ Fixable Dead Cell Stain Kit (Invitrogen). FC receptors were blocked with anti-CD16/32 antibody (2.4G2, BioXcell) and the cell suspensions were successively stained in Brilliant Stain Buffer (BD Biosciences) with one or a combination of the following anti-mouse antibodies: Ly6G (1A8, BD Biosciences), P2RY12 (S16007D, Biolegend), Sirpα (P84, BD Biosciences), CD86 (GL1, BD Biosciences), CD4 (GK1.5, BD Biosciences), CD45.2 (104, BD Biosciences), CD11b (M1/70, BD Biosciences), CX3CR1 (SA011F12, Biolegend), CD8α (MCD0826, ThermoFisher scientific), Arg1 (A1exF5, ThermoFisher scientific), MHCII (M5/114.15.2, BD Biosciences), CD103 (M290, BD Biosciences), CD62L (MEL-14, Biolegend), PD-1 (J43, BD Biosciences), CD24 (M1/69, BD Biosciences), PD-L1 (MIH5, BD Biosciences), SiglecH (440C, BD Biosciences), CD64 (X54-5/7.1, BD Biosciences), TMEM119 (106-6, abcam), CD3 (17A2, Biolegend), CD68 (FA-11, Biolegend), CD49d (9C10, BD Biosciences), LAG3 (ebioC9B7W, Invitrogen), CD44 (IM7, Biolegend), FOXP3 (FJK.16s, Invitrogen), CD69 (H1.2F3, Biolegend), CD25 (PC1, Biolegend), NK1.1 (PK136, Biolegend), CD11c (N418, ThermoFisher scientific), CD39 (Duha59, Biolegend), MertK (DS5MMER, ThermoFisher scientific), Tim3 (RMT4-54, BD Biosciences), F4/80 (BM8, Invitrogen), Ly6C (AL-21, BD Biosciences), CD52 (4C8, BD Biosciences), CD19 (6D5, Biolegend). When unlabeled primary rabbit antibodies or biotinylated antibodies were used, cells were subsequently incubated with anti-rabbit IgG antibodies or fluorochrome-conjugated streptavidin. For intracellular staining, the cell suspension was fixed and permeabilized with the eBioscience^TM^ FOXP3/Transcription Factor Staining Buffer Set (eBioscience) after the cell surface staining, following the manufacturer’s instruction.

The cell suspension was then filtered through a 40 μm filter and each sample was acquired using the spectral flow cytometer Cytek^®^ Aurora (5 lasers, 5L UV-V-B-YG-R) and the SpectroFlo^®^ software (v.3.3.0) (Cytek® Biosciences). For data analysis initial gating strategy were performed as previously described^119^ with FlowJo software (v.10.10.0): (1) FSC-A versus SSC-A to exclude debris; (2) FSC-A versus FSC-H to select singlet; (3) SSC-A versus Live/Dead for dead cell exclusion; (4) SSC-A versus CD45 to select CD45^+^ leukocytes. They were then followed by unsupervised high parametric analysis using Spectre package^120^ (v.1.0.0) in R (v.4.2.1). Uniform Manifold Approximation and Projection (UMAP) and density UMAP was run with a concatenation of all samples per group simultaneously. For cluster analysis, Rphenograph was selected (k = 25-45) and the clusters were annotated based on expression level of the markers of interest. Cell populations were defined as indicated in Supplementary Table S2. Frequencies of total leukocytes and subpopulations were determined using FlowJo and Spectre respectively and total leukocytes and subpopulation absolute numbers per g of brain were calculated. Principal component analysis (PCA) was performed in R using the singular value decomposition and visualized using GraphPadPrism (v.10.2.3). Spearman correlation network between CD8 T cell subsets, microglia subsets, and brain metastatic area were calculated within each group using the R package Hmisc_5.1.0 and multiple hypothesis correction was performed using FDR. Correlations that passed p-value < 0.05, FDR < 0.2, spearman rho > |0.5| were visualized using the R package igraph_1.5.1.

### Single-cell RNA sequencing (scRNAseq) of mouse BrM

#### Brain dissociation, cell capture and sequencing

Single cell suspension from BR1 or BR3-BrM bearing or PBS (non-implanted controls) brains were obtained using Adult Brain Dissociation Kit (Miltenyi, #130-107-677) following manufacturer’s protocol. Library preparation, captures and sequencing were performed by Single Cell Analysis Facility (SCAF, CCR, NCI). The sample concentration and viability were assessed using the LunaFx7 fluorescent cell counter. The cellular samples were loaded in the lanes according to the 10X Genomics 3’ v3.1 Single Cell User Guide with a single capture lane per sample targeting recovery of up to 6,000 cells per lane. Cell partitioning completed successfully with uniform emulsion consistency and the reverse transcription PCR was run overnight. All subsequent steps and quality control was performed as described in the 10X Genomics 3’ v3.1 Single Cell User Guide. Two NovaSeq runs for GEX were performed: NovaSeq 2k S1 100-cycle and NovaSeq 6k S1 100-cycle.

#### scRNA-seq data preprocessing (10x chromium single cell 3’ v3) and analysis

10x Genomics Cell Ranger (v.7.0.1) was used to perform demultiplexing, alignment to mouse reference mm10 (refdata-gex-mm10-2020-A), gene and transcript quantification. The parameters – include-introns –force-cells were applied in Cell Ranger count step. Seurat (v.4.2.0)^121^ was used for preprocessing, normalization and clustering. Genes detected in fewer than 3 cells and cells containing less than 100 genes were removed from the data sets prior to further filtering. Doublets and background cells were filtered based on negative binomial distribution to determine the upper and lower limit of UMI count per cell (9-93,348) and gene count per cell (33-16,472). Mitochondrial percentages in cells were calculated in Seurat, we set up the filtering threshold as 15.46% (median + 4 x median absolute deviation), cells with higher mitochondrial expression than the selected threshold were filtered out. After removing unwanted cells from the dataset, the data was normalized by SCTransform from Seurat. Seurat RunPCA and ElbowPlot were used to find the optimal number of principal components (PCs), Seurat FindClusters function was called to use the first 20 PCs at the resolution of 0.6 and identified 37 distinct cell clusters. Uniform manifold approximation and (UMAP)^122^ was used for data visualization in two-dimensional space. These upstream analyses were performed by CCR-SF Bioinformatics Team (NIH-CCR).

Then all downstream analysis were performed using Seurat package (v.5.1.0)^123^ in R. Cell type identifications were performed using the reference cell type gene expression databases: Allen brain atlas (https://portal.brain-map.org/), Immgen (https://www.immgen.org/) and PanglaoDB (https://panglaodb.se/) and the automated annotation algorithm: Azimuth (https://azimuth.hubmapconsortium.org/) and SC_Type^124^.

Re-clustering on immune cell subsets was performed using FindNeighbors (dimension 1:30) and FindClusters (resolution 0.5), and visualized using RunUMAP (dimension 1:30). Cluster specific genes were identified using the FindMarkers function with a fold change of > 0.25. Re-clustering for specific downstream immune cell analysis was performed following the same approach. Re-clustering on tumor cells was performed using non-negative matrix (NMF) as described in the next section. For all immune cell analysis, the differentially expressed genes (DEGs) between groups and cluster was performed using the FindMarkers function (p_adj < 0.05 and Avg_Log2FC > 0.25). For volcano plot the thresholds of significance used were: p_adj < 0.05 and Avg_Log2FC > 0.5. For melanoma cells the DEGs between BR1 and BR3 groups were analyzed using MAST (Model-based Analysis of Single-cell Transcriptomics)^65^. Log_2_ values of BR1 vs. BR3 fold change of gene expression levels (Log_2_FC) and Log_10_ of p values adjusted of t-test (-Log_10_p-value.adj) were analyzed to make the volcano plot. The threshold of significance used is Log_2_FC > 0.6 and p < 0.05. Gene Set Enrichment Analysis (GSEA) (https://www.gsea-msigdb.org/gsea/index.jsp) were then performed and using GO, KEGG and Hallmark for microglia (group comparison and clusters), and using Gene Ontology (GO), Kyoto Encyclopedia of Genes and Genomes (KEGG) and REACTOME datasets for melanoma BrM cells. The top pathways determined by the highest Normalized Enrichment Score (NES) were used to select those represented on the figures.

### Non-negative matrix factorization and metaprogram discovery

We retained genes expressed in at least 1% of all malignant cells from both BR1 and BR3 mouse models and ran a fast NMF implementation^67^ separately on malignant cells from each model with no L1 or L2 penalty specified. We chose to obtain 30 factors for cells from each mouse model based on the change in reconstruction error of the gene expression matrix as the number of factors was increased. For a given NMF run within BR1 or BR3 models, we prioritized genes that contributed to each factor using a two-step procedure. We first z-scored the loadings of each gene across all factors and for a given factor, retained those genes with z > 1.5. Second, for a given factor, we computed the Spearman correlation coefficient of all gene pairs and used those pairs whose correlation was greater than 0.4 and constructed a gene co-expression graph. The final prioritized gene list (referred to as prioritized genes) consists of only those genes in the largest component of the co-expression graph. Any factor with less than ten prioritized genes is discarded. For each factor, we then use AUCell^125^ to score the activity of prioritized genes across malignant cells from both the mouse models and compute the Pearson correlation coefficient between all pairs of factors including both models. We hierarchically clustered the factors based on the inter-factor correlations and grouped all factors into eight clusters or metaprograms (this number maximized the Silhouette coefficient of the clustering). Within each metaprogram, we only retained those genes that were present in at least 25% of all constituent factors. The metaprograms that were active (based on the Global_k1 AUCell threshold) in less than 10% of cells from each model were discarded. This procedure yielded seven metaprograms spanning the cells in the two mouse models. Each metaprogram functional implication was characterized by pathway analysis using Metascape^71^.

Since NMF factorizations are non-unique i.e., can yield different factors when re-run, we repeated the above procedure 100 times. For each run, we trained a logistic regression classifier to distinguish BR1 and BR3 tumor cells (using five-fold cross-validation) based on the metaprograms obtained from the procedure outlined above and computed its area under the ROC (auROC) value. We chose this criterion to ensure that our pipeline to derive metaprograms from an NMF run is able to capture differences between BR1 and BR3 models. The metaprograms derived in this paper were based on the NMF run whose auROC (auROC = 0.927) was the median auROC value amongst all NMF runs.

To cluster all malignant cells, we first scored the activity of these seven metaprograms using AUCell. We then z-scored each metaprogram activity across all cells and constructed a k-nearest neighbor graph (k=30) based on the seven z-scores for each cell. The Mahalanobis distance was used to compute distances between cells for neighborhood graph construction as this distance metric takes into account potential correlations between metaprogram scores. We ran 100 parallel iterations of Louvain clustering (with a resolution of 0.4) on this graph and kept the clustering with the highest Silhouette coefficient (where cell-cell distances were computed using the Mahalanobis distance metric). For visualization, we computed UMAP coordinates of each malignant cell based on the activity across the seven metaprograms.

### Correlation analysis of mouse tumor metaprograms with tumor metaprograms derived in Biermann et. al

We used AUCell to compute the activity of our metaprograms as well as metaprograms derived by Biermann et al. 2022 in every malignant cell from Biermann et al. 2022, Smalley et al. 2023, Alvarez-Breckenridge et al. 2023. We then computed the pair-wise Spearman correlation coefficient between every mouse metaprogram and every Biermann et al. 2022 metaprogram across cells in each tumor biopsy. Within a given cohort, the median correlation coefficients computed for a given pair of metaprograms across all tumors was retained for further analyses. A metaprogram pair was considered to be significantly and positively correlated within a cohort if the set of tumor-wise correlations were significantly above 0.2 (Wilcoxon one-sided p-value < 0.05). We then counted the number of cohorts in which a metaprogram pair were significantly positively correlated and only visualized those that were correlated in at least one cohort.

### Cell-cell interaction discovery using Cellchat

We used Cellchat^126^ to infer cell-cell communications between microglia, CD4 T cells, CD8 T cells and malignant cells in BR1 and BR3 samples separately. Given the relatively lower numbers of CD4 and CD8 T cells in the data, we grouped all sub-clusters of those two cell types while keeping CD4 and CD8 T cells separated and detected interactions between them and sub-clusters of microglia and up-regulated (p_adj < 0.05) in the sender and receiver cell type, respectively and (b) the ligand and receptor were expressed in at least 10% of cells in the sender and receiver cell type, respectively. Interactions amongst the top 10% most different in strength (which is defined by Cellchat as the product of ligand and receptor expression in sender and receiver cell types and weighted by the proportion of cells in each cell type) between BR1 and BR3 was labelled as either BR1-specific or BR3-specific.

### Whole metastatic brain protein expression

Brain hemispheres were mechanically disrupted in 500 μL ProcartaPlex^TM^ cell lysis buffer (ThermoFisher scientific, EPX-99999-000) for every 100mg of tissue, using an Omni Tissue Master 125 Homogenizer (Omni international). Supernatants were extracted and total concentration of protein was measured using the Pierce^TM^ BCA Protein Assay Kit (ThermoFisher scientific, 23227). Samples were diluted to 10 mg/mL protein with PBS and cytokines concentration were measured using the ProcartaPlexTM Mouse Cytokine & Chemokine Panel 1A, 36plex (ThermoFisher scientific, EPX360-26092-901) according to manufacturer’s instructions.

### Patient single-cell transcriptomic analysis on T cells

The single-cell read-count matrix of Biermann et al. 2022 dataset were processed using Seurat (v4.3.0)^123^. We chose only brain metastatic biopsies and normalized read counts using the NormalizeData function. The 5000 most variable genes were chosen to compute 50 principal components, which were then clustered using the FindClusters function (with a resolution of 0.8). We then subset the data to those clusters with high expression of CD45 for re-analysis. Based on the expression of *CD3D*, *CD3E*, *NCR1*, *CD8A*, *CD8B*, *CD4*, *GZMA*, *GZMB*, *IFNG*, *GZMK*, *PRF1*, *ITGAM*, *ITGAX*, *CD14*, *NCAM1*, *FCGR3A*, *P2RY12*, *HBB*, we separated lymphoid cells from other CD45+ sub-populations. We then retained only those samples from BrM patients (n=15) with at least 50 lymphoid cells (resulting in a set of cells from 10 patients) and carried out batch-correction and integration in Seurat (using 30 PCs and k.weight = 50). The expression of *HAVCR2*, *LAG3*, *PDCD1*, *TIGIT*, *CTLA4*, *GZMA*, *GZMB*, *GZMK*, *IFNG*, *PRF1*, *SELL*, *CCR7*, *IL7R*, *TCF7*, *MKI67*, *CDK1*, *CD45RO*, *TOX*, *CXCL13*, *RGS1*, *TNFSF9*, *ENTPD1*, *ITGAE* was used to annotate all CD8^+^ T cell subclusters.

### Patient bulk-RNA sequencing and survival analysis

MP4 and MP6 signatures were built using the whole gene lists that define these MPs. “BR1 pro-inflammatory” and “BR1 chemokine” include pro-inflammatory and chemokine, respectively, DEGs and cell-cell interactions enriched in BR1. “BR3 neuro” includes axon guidance BR3 DEGs that were also enriched in this group the interactome analysis. To yield a single gene expression vector for each biopsy in Sun et al. 2023 dataset, bulk RNA expression was calculated per sample by summing counts for all cells using the AggregateExpression function from the R package Seurat^123^(v.5.0.1). Counts were normalized per sample using log_2_(CPM+0.001). Mouse metaprogram genes were converted to human homologs using R software and the Jackson laboratory database (http://www.informatics.jax.org/downloads/reports/HOM_MouseHumanSequence.rpT); genes without direct homologs or that were not contained in the dataset was omitted from signature calculations as well as genes not found in the Sun et al. human dataset. Z-scores were calculated for genes of interest across all samples and mean z-score for each gene signatures were performed. Spearman correlation between the different z-score signature in patients were performed using GraphPadPrism. For survival analysis: for each signature, patients were separated based on median into High (≥ median) and Low (< median) expression. Log-rank (Mantel-Cox) test was performed to calculate the correlation of the gene signature expression and the patient progression free survival after surgery, and p-value reported using GraphPadPrism.

## QUANTIFICATION AND STATISTICAL ANALYSIS

### Statistical analysis

Statistical analyses were performed using GraphPadPrism or R. Data were analyzed for normal distribution prior to comparisons. Outliers were detected using ROUT’ test and excluded from the analysis. Values are presented as the mean ± SEM; individual values (representing individual samples when non indicated or individual experiment when indicated) are presented as scatterplots with column bar graphs or box plot. Data in pie charts represent mean of samples in group. Statistical significance between two groups was determined by two-tailed unpaired Student’s t test (parametric) or Mann-Whitney (non-parametric). For the comparison of multiple groups, comparing different treatment in one unique model, one-way ANOVA followed by Tukey’s multiple comparison correction test (parametric) or Mann-Whitney (non-parametric) followed by Dunn’s multiple comparison correction test were used). For the comparison of multiple groups, comparing BR1 and BR3 in untreated and treated conditions, two-way ANOVA followed by Tukey’s multiple comparison correction test (parametric) or Mann-Whitney (non-parametric) followed by Dunn’s multiple comparison correction test were used. Gene signature z-scores were calculated for bulk/pseudobulk and single-cell RNA expression in R using scale and AddModule (Seurat) functions, respectively. Tumor metaprogram z-scores were calculated as z-score = (activity - mean(activity))/sd(activity).

## SUPPLEMENTARY TABLES

**Table S1:**
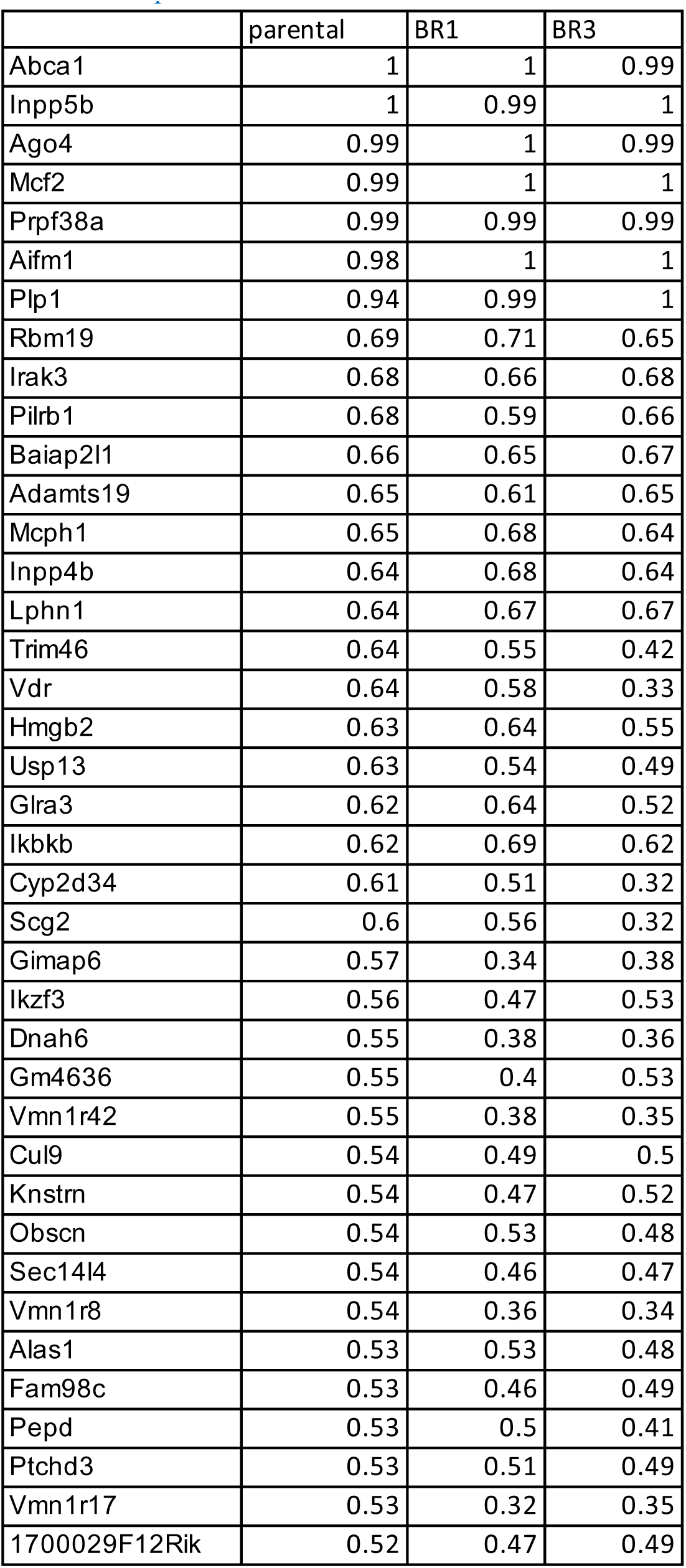
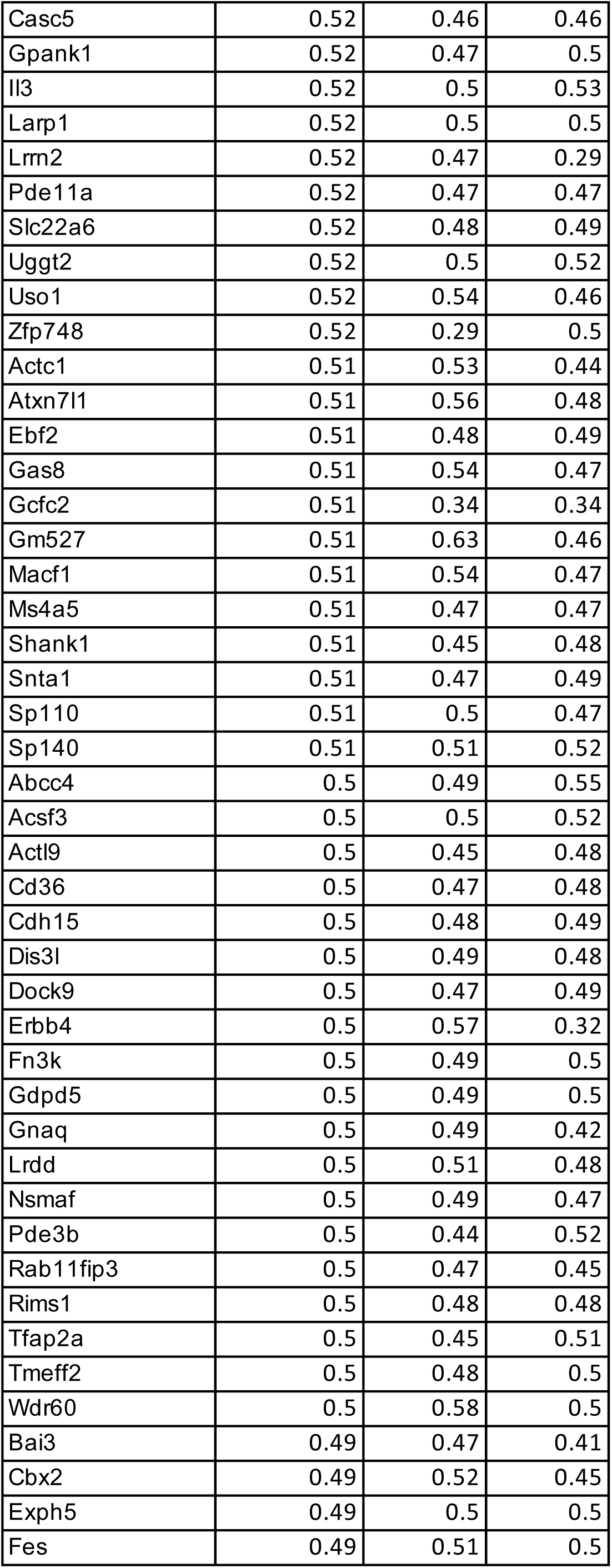
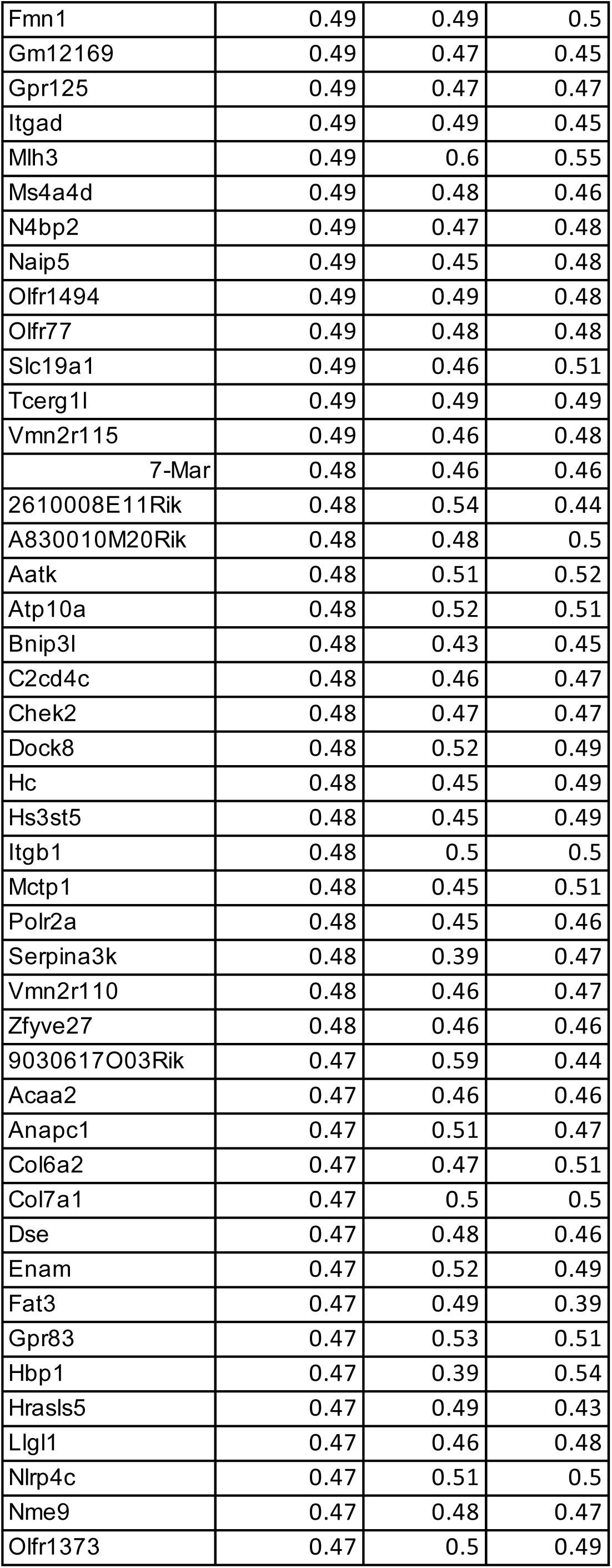
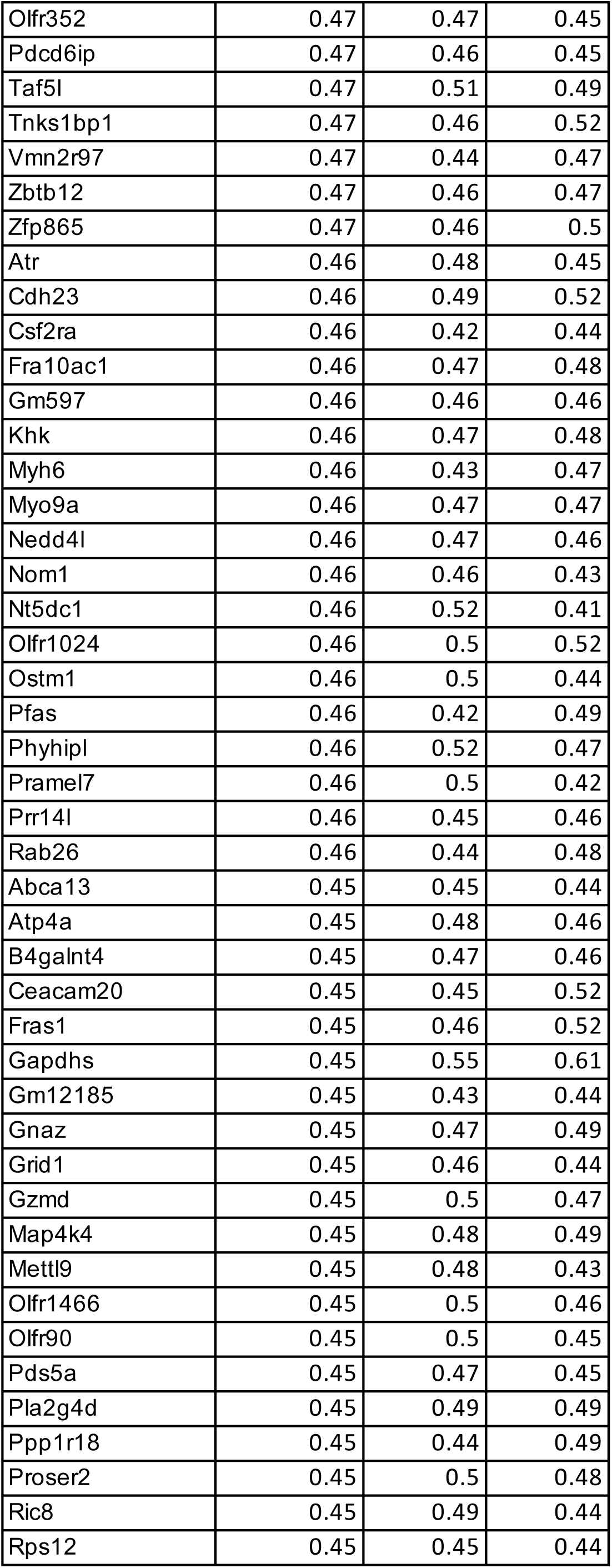
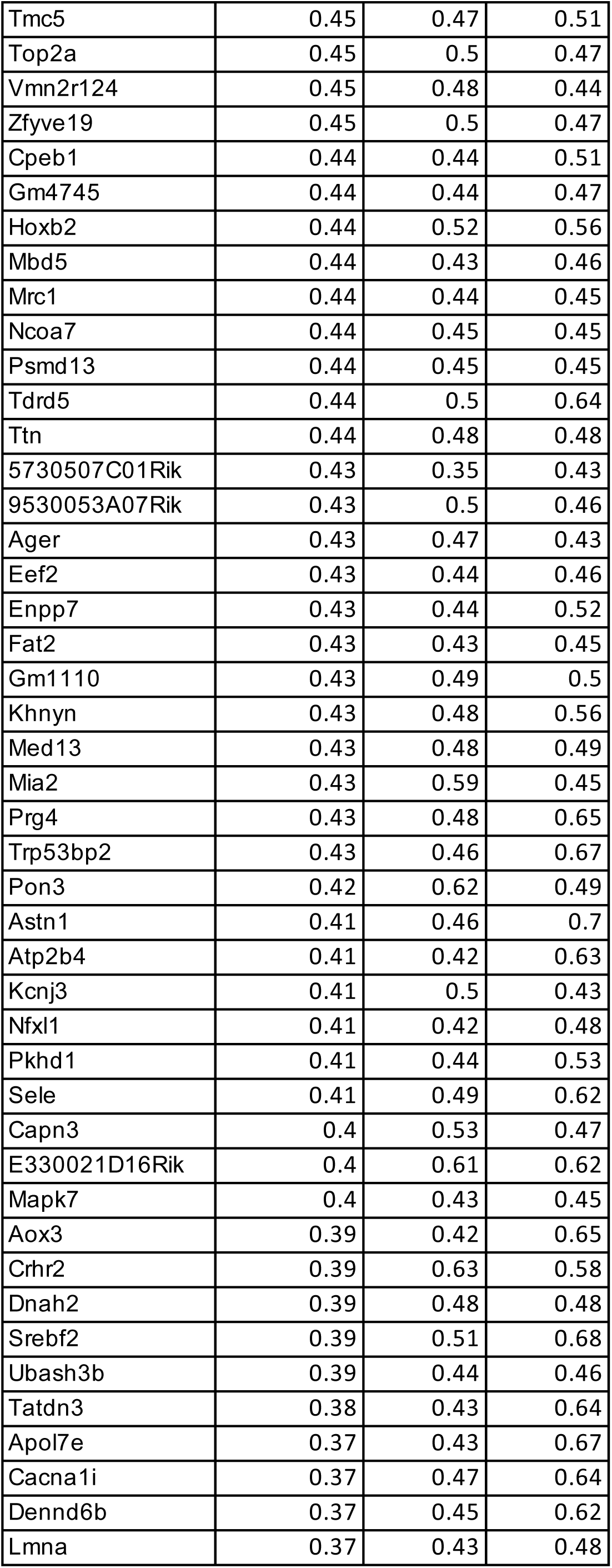
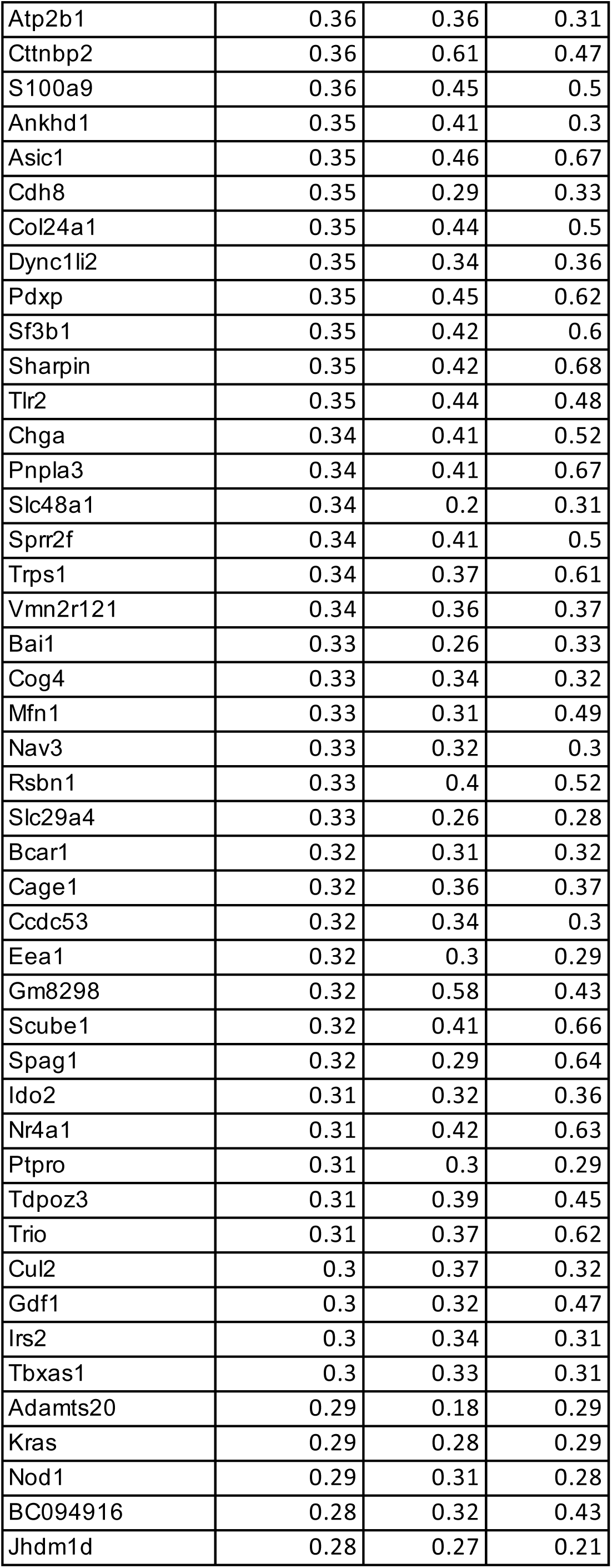
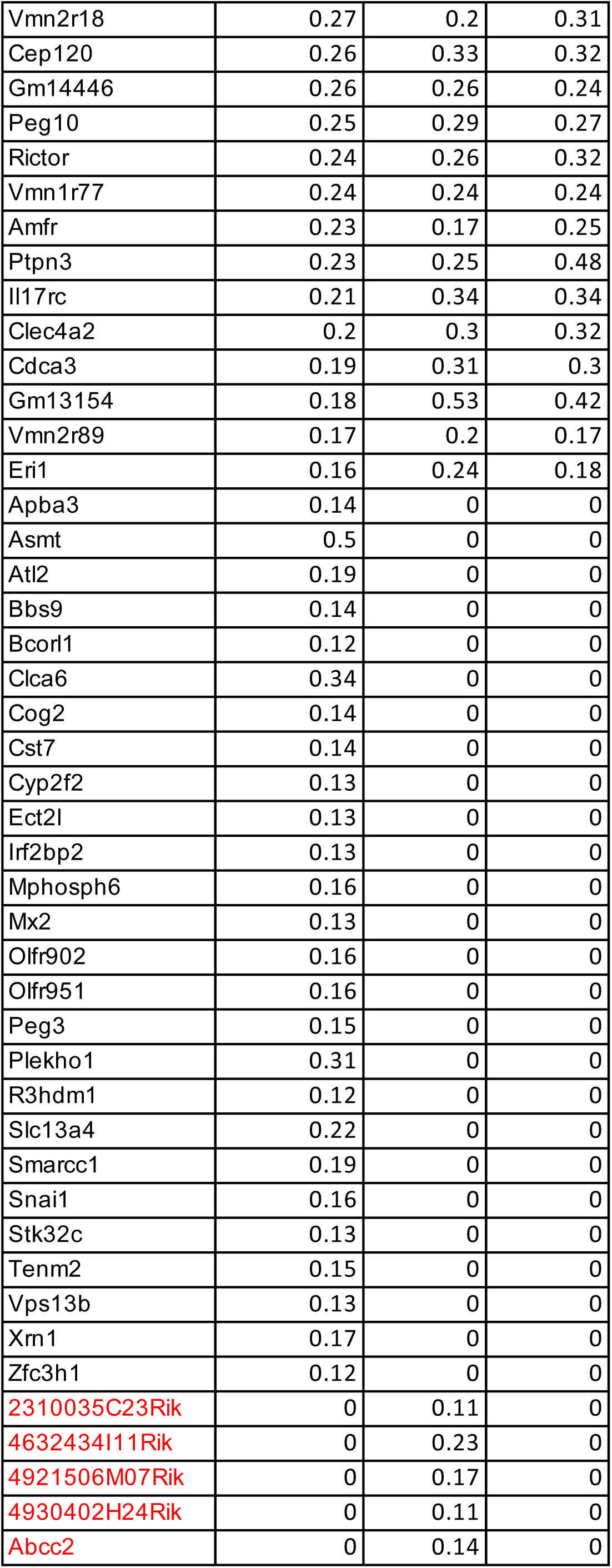
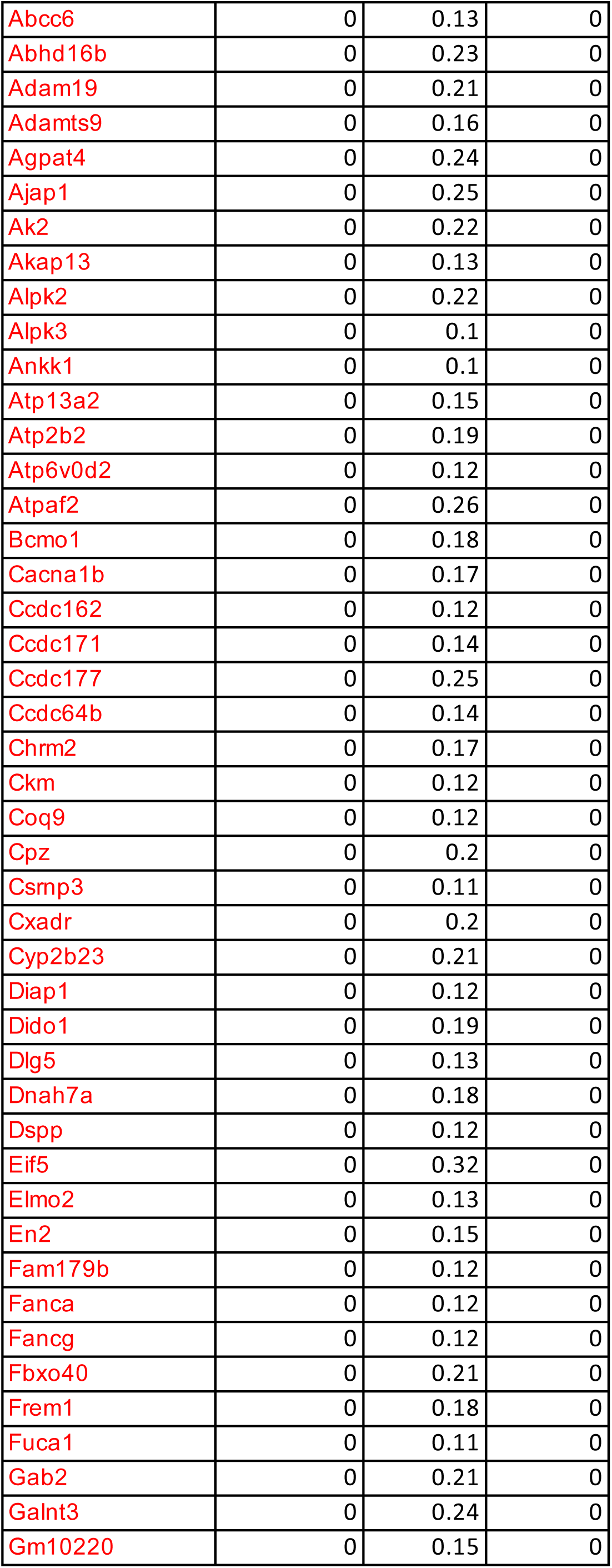
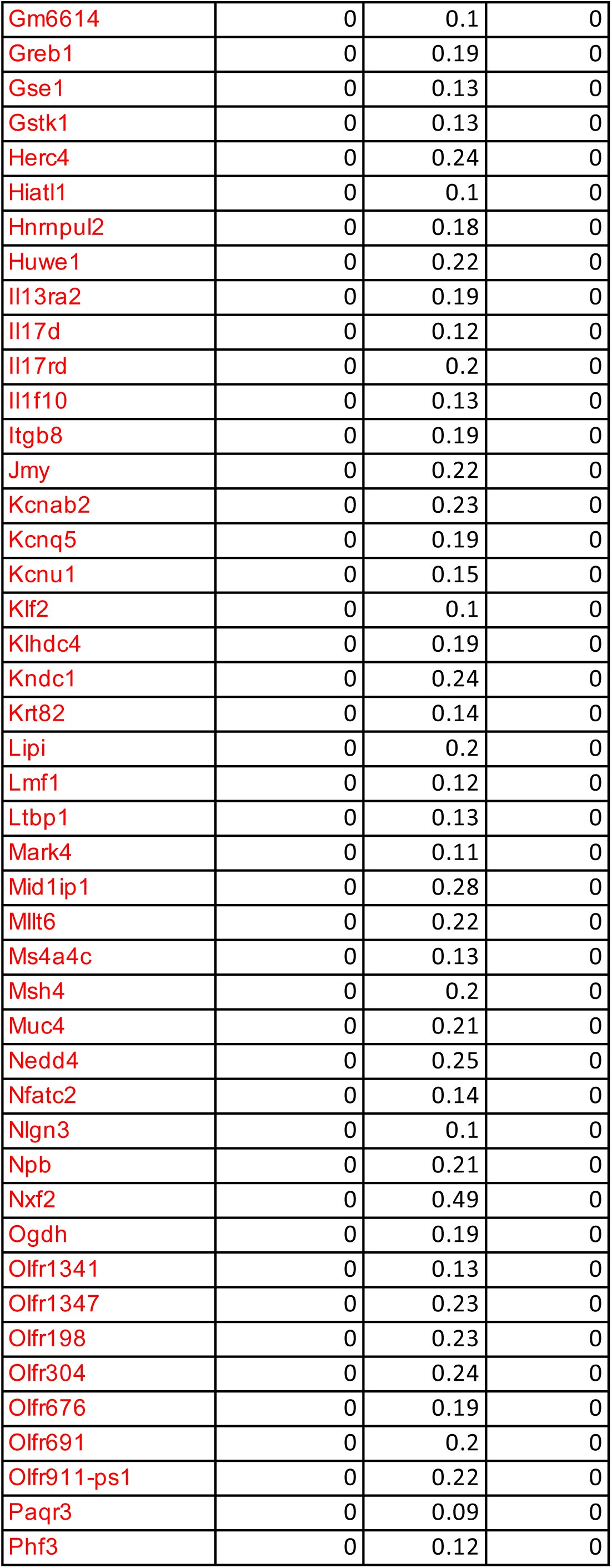
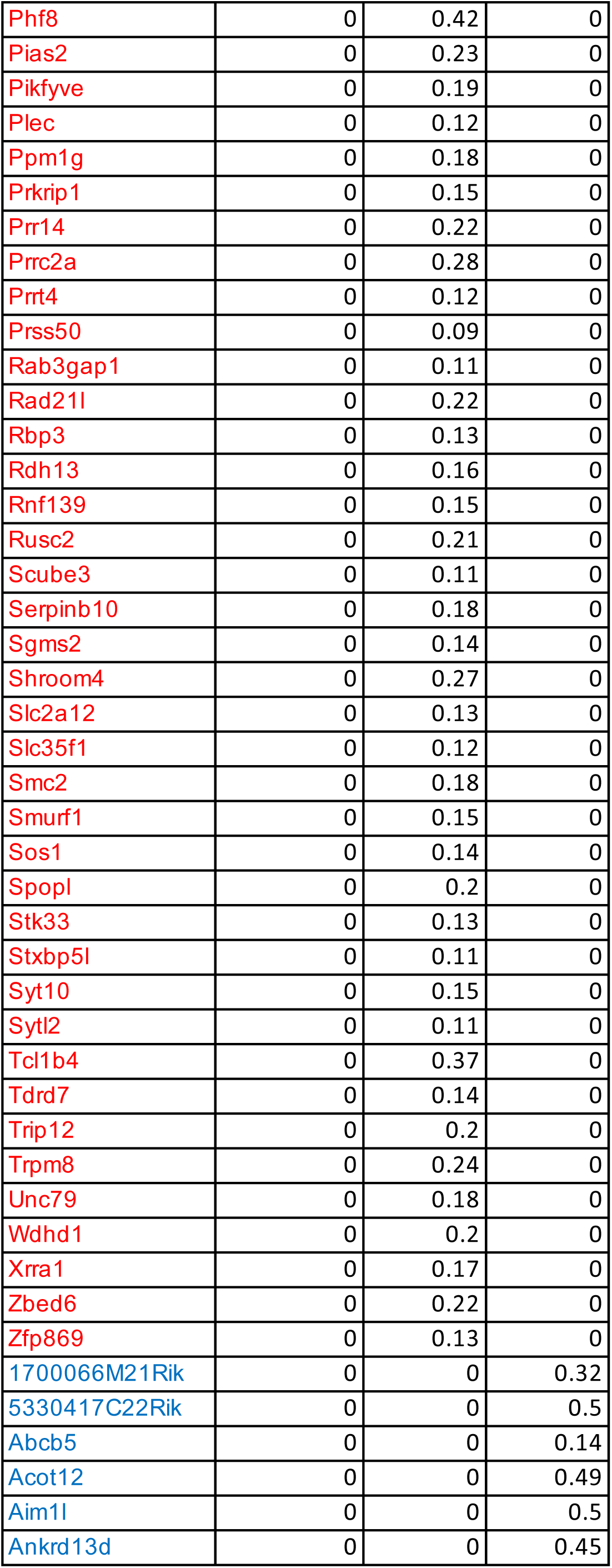
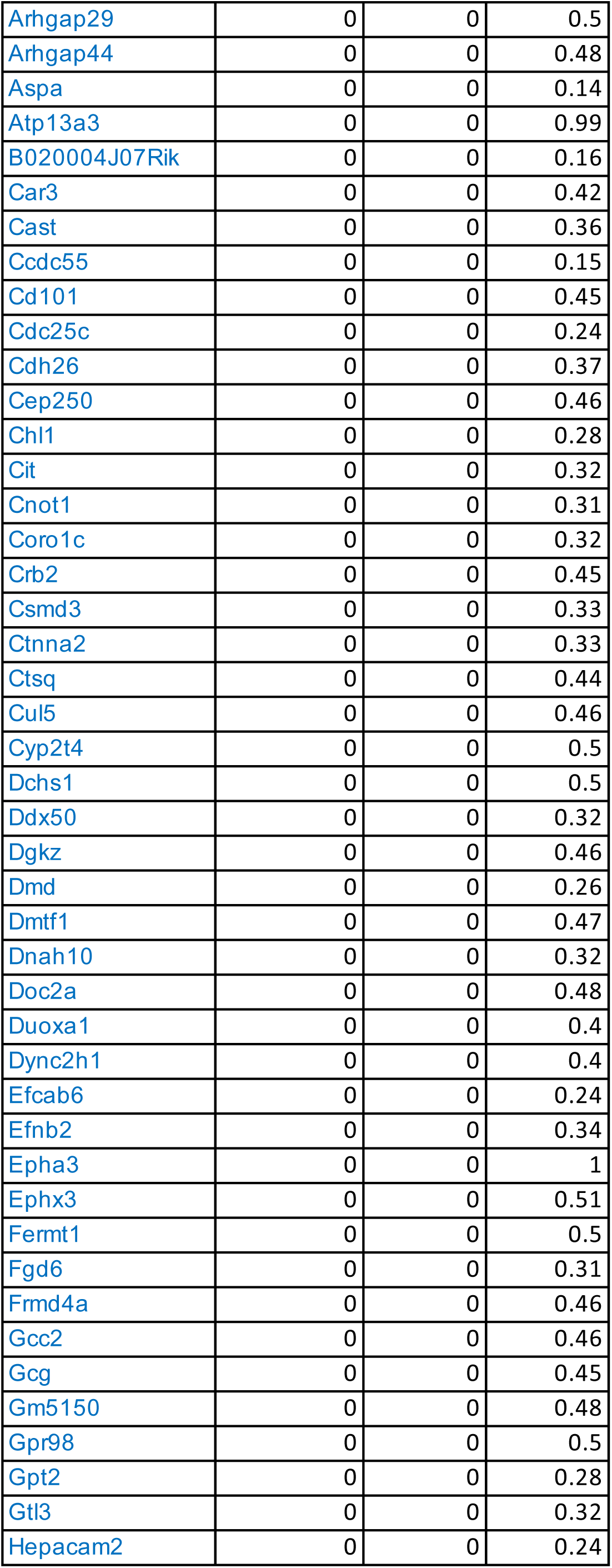
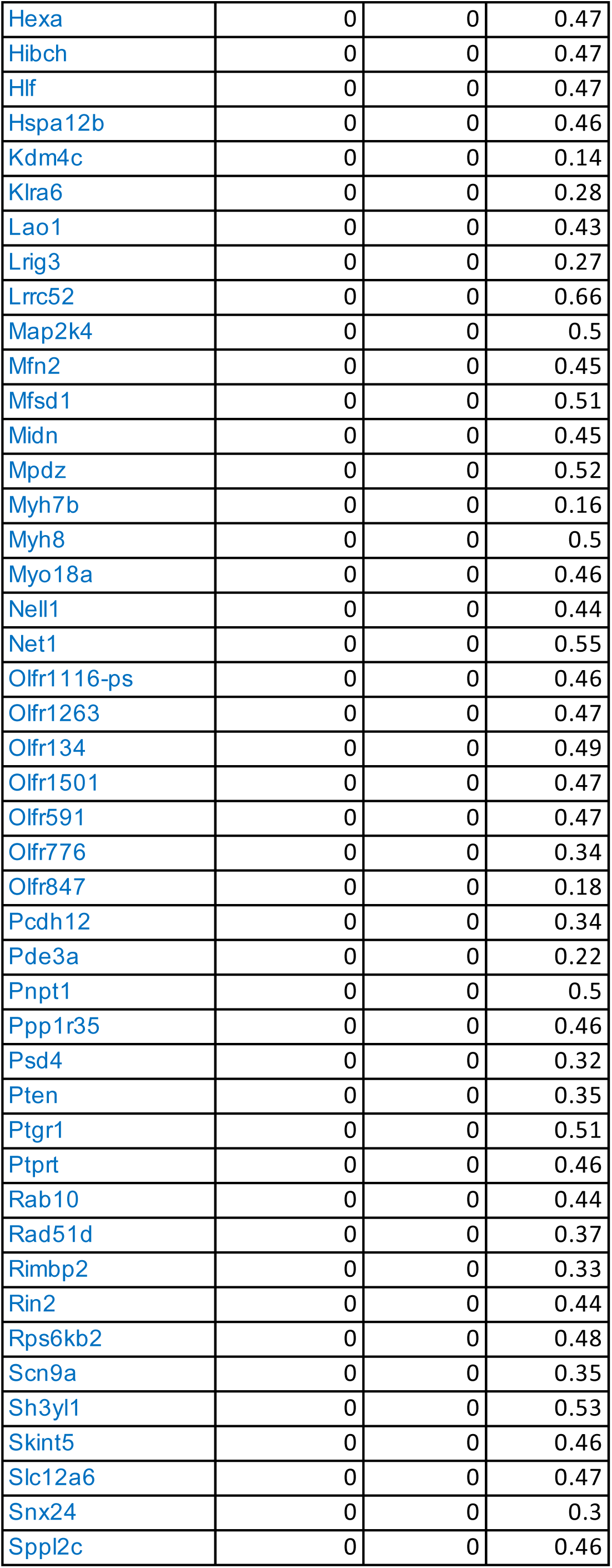
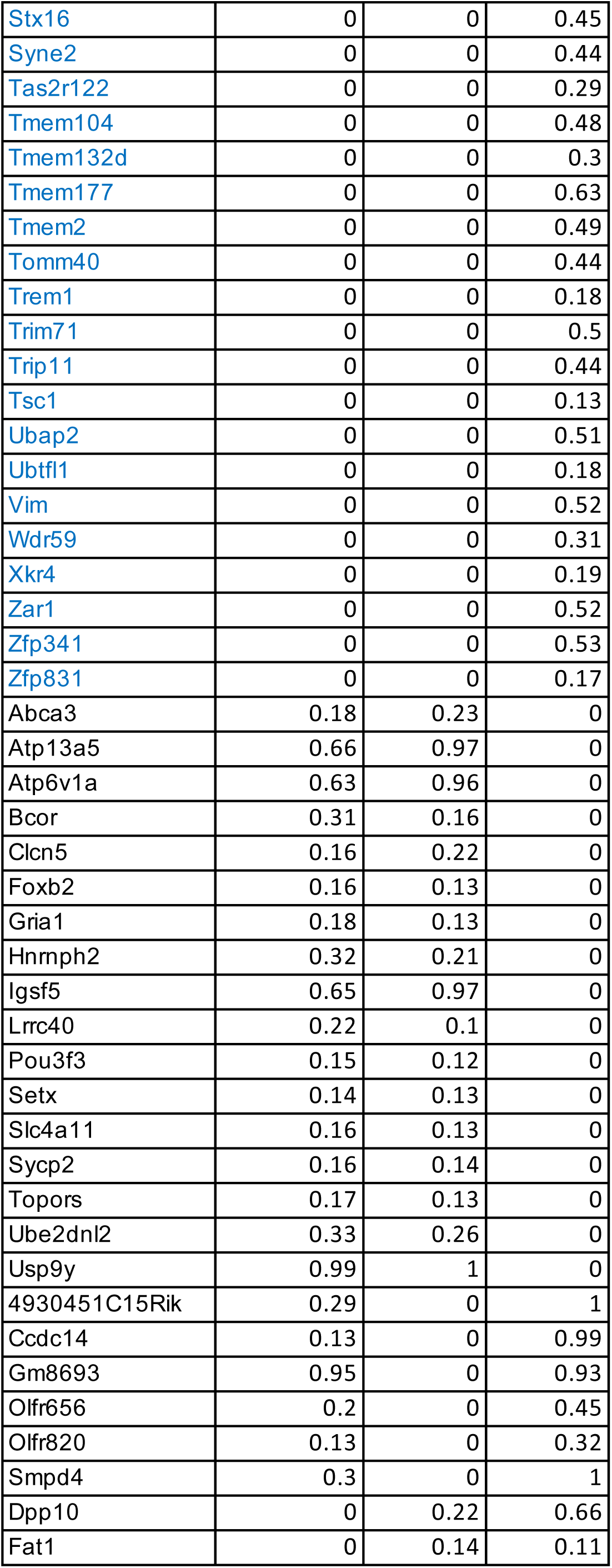
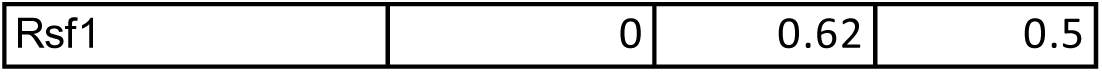
Tumor cell line mutated genes. The number is variant allele frequncy (VAF). Subline-specific mutated genes can be shown by filtering genes with VAF = 0 in other two lines. In red: unique mutations found in BR1 In blue: unique mutations found in BR3

**Table S2.**
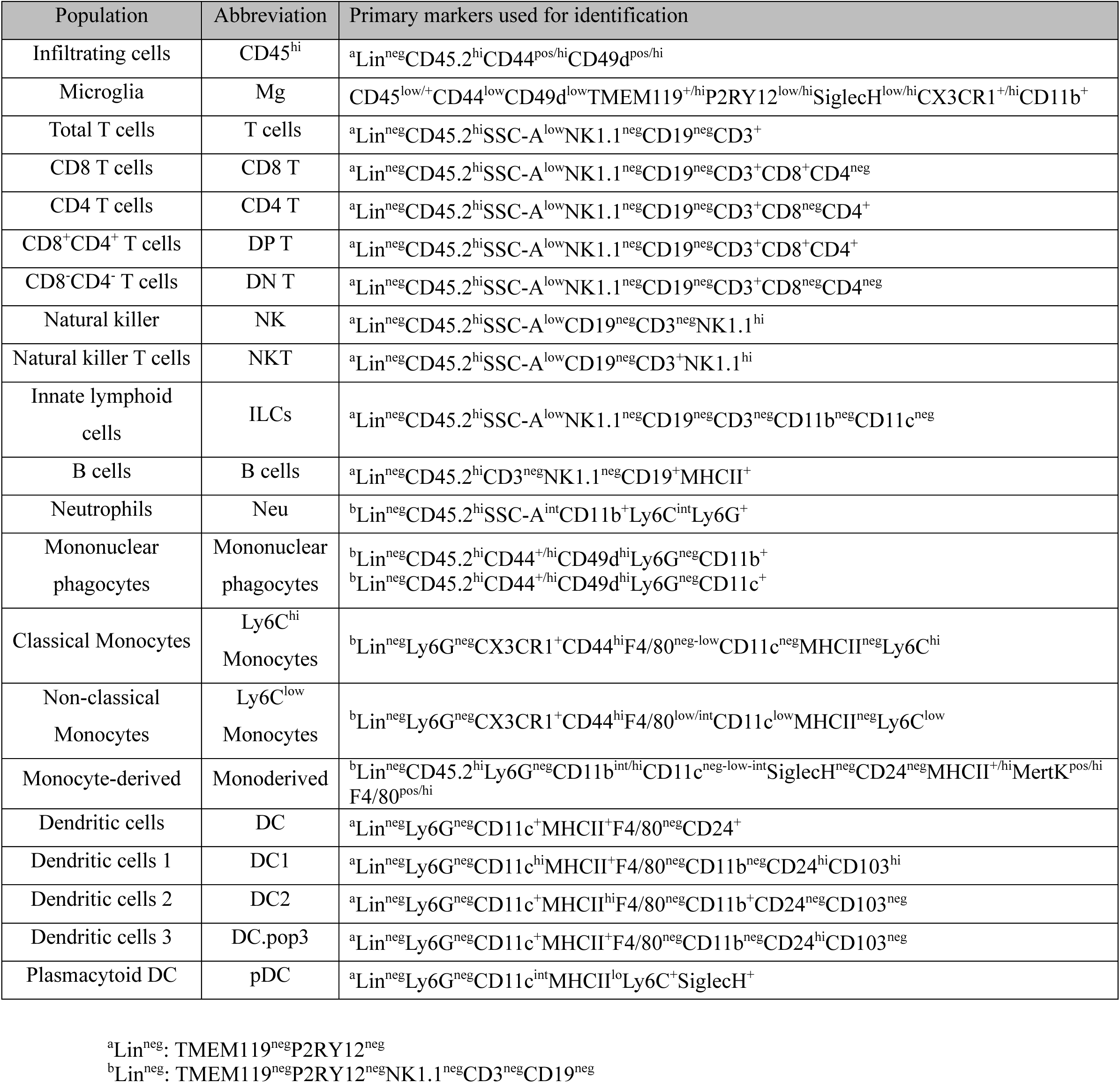
Immune cell annotations.

**Table S3:** CD8+ T cell cluster-defining genes.

| p_val | avg_log<br>2FC | pct.1 | pct.2 | p_val_adj | cluster | cluster<br>name | gene |
| --- | --- | --- | --- | --- | --- | --- | --- |
| 1.95E-68 | 1.931 | 0.983 | 0.550 | 5.11E-64 | 0 | sc.5 | S100a6 |
| 2.54E-66 | 1.235 | 1.000 | 0.817 | 6.67E-62 | 0 | sc.5 | Cd3g |
| 2.61E-65 | 1.362 | 1.000 | 0.788 | 6.85E-61 | 0 | sc.5 | Nkg7 |
| 9.43E-62 | 1.314 | 1.000 | 0.790 | 2.47E-57 | 0 | sc.5 | AW112010 |
| 9.58E-62 | 1.411 | 1.000 | 0.581 | 2.51E-57 | 0 | sc.5 | Cxcr6 |
| 1.55E-60 | 2.079 | 0.991 | 0.658 | 4.05E-56 | 0 | sc.5 | Ccl5 |
| 2.26E-59 | 1.733 | 0.945 | 0.396 | 5.92E-55 | 0 | sc.5 | S100a4 |
| 3.96E-59 | 1.364 | 0.766 | 0.162 | 1.04E-54 | 0 | sc.5 | Gm36723 |
| 6.75E-52 | 1.044 | 0.830 | 0.254 | 1.77E-47 | 0 | sc.5 | Fgl2 |
| 2.33E-49 | 1.852 | 0.770 | 0.223 | 6.10E-45 | 0 | sc.5 | Gzmb |
| 2.16E-48 | 0.979 | 0.996 | 0.750 | 5.67E-44 | 0 | sc.5 | Cd8a |
| 1.43E-47 | 1.160 | 0.689 | 0.175 | 3.75E-43 | 0 | sc.5 | Tcrg-C2 |
| 7.43E-47 | 1.383 | 0.923 | 0.406 | 1.95E-42 | 0 | sc.5 | Ccl4 |
| 6.27E-46 | 1.055 | 0.953 | 0.642 | 1.64E-41 | 0 | sc.5 | S100a11 |
| 2.05E-43 | 1.104 | 0.983 | 0.565 | 5.36E-39 | 0 | sc.5 | Id2 |
| 4.23E-43 | 1.288 | 0.932 | 0.633 | 1.11E-38 | 0 | sc.5 | Klrd1 |
| 7.42E-42 | 0.948 | 0.987 | 0.571 | 1.95E-37 | 0 | sc.5 | Rbpj |
| 1.46E-41 | 0.941 | 0.932 | 0.429 | 3.84E-37 | 0 | sc.5 | Pdcd1 |
| 1.95E-41 | 1.196 | 0.983 | 0.681 | 5.12E-37 | 0 | sc.5 | Lgals1 |
| 1.06E-39 | 1.087 | 0.677 | 0.202 | 2.78E-35 | 0 | sc.5 | Trgv2 |
| 1.23E-37 | 0.933 | 0.783 | 0.292 | 3.24E-33 | 0 | sc.5 | Lag3 |
| 2.82E-37 | 0.949 | 0.685 | 0.192 | 7.39E-33 | 0 | sc.5 | Havcr2 |
| 2.16E-35 | 0.935 | 0.647 | 0.200 | 5.66E-31 | 0 | sc.5 | Cd200r1 |
| 3.43E-35 | 1.140 | 0.651 | 0.192 | 9.00E-31 | 0 | sc.5 | Ccl3 |
| 1.97E-33 | 0.941 | 0.877 | 0.467 | 5.16E-29 | 0 | sc.5 | Cdk6 |
| 1.02E-30 | 1.011 | 0.728 | 0.294 | 2.68E-26 | 0 | sc.5 | Prf1 |
| 7.87E-28 | 1.052 | 0.553 | 0.181 | 2.06E-23 | 0 | sc.5 | Klre1 |
| 1.80E-27 | 1.007 | 0.685 | 0.277 | 4.72E-23 | 0 | sc.5 | Ifng |
| 1.69E-22 | 0.955 | 0.749 | 0.375 | 4.44E-18 | 0 | sc.5 | Ctla4 |
| 1.20E-16 | 0.956 | 0.677 | 0.419 | 3.14E-12 | 0 | sc.5 | Plac8 |
| 7.42E-22 | 1.600 | 0.639 | 0.290 | 1.95E-17 | 1 | sc.3 | Ly6c2 |
| 1.22E-20 | 0.629 | 0.614 | 0.241 | 3.20E-16 | 1 | sc.3 | Cxcr3 |
| 1.91E-16 | 0.638 | 0.975 | 0.791 | 5.02E-12 | 1 | sc.3 | Hcst |
| 1.07E-15 | 0.617 | 0.772 | 0.425 | 2.82E-11 | 1 | sc.3 | Gramd3 |
| 1.96E-15 | 0.561 | 0.810 | 0.564 | 5.15E-11 | 1 | sc.3 | Smap1 |
| 1.33E-14 | 0.537 | 0.709 | 0.392 | 3.48E-10 | 1 | sc.3 | Hopx |
| 2.46E-14 | 0.454 | 1.000 | 0.993 | 6.46E-10 | 1 | sc.3 | Rpl13a |
| 3.79E-14 | 0.457 | 0.994 | 0.700 | 9.95E-10 | 1 | sc.3 | Ccl5 |
| 6.85E-14 | 0.732 | 0.854 | 0.621 | 1.80E-09 | 1 | sc.3 | Zfp36l2 |
| 1.04E-13 | 0.795 | 0.918 | 0.794 | 2.72E-09 | 1 | sc.3 | Itgb1 |
| 2.79E-13 | 0.490 | 1.000 | 0.992 | 7.31E-09 | 1 | sc.3 | Rps27 |
| 3.63E-13 | 0.593 | 0.962 | 0.844 | 9.52E-09 | 1 | sc.3 | Gimap4 |
| 1.11E-11 | 0.497 | 0.956 | 0.901 | 2.92E-07 | 1 | sc.3 | Ly6e |
| 1.56E-11 | 0.526 | 1.000 | 0.987 | 4.09E-07 | 1 | sc.3 | Tmsb10 |
| 6.09E-11 | 0.449 | 0.981 | 0.814 | 1.60E-06 | 1 | sc.3 | Ptpn18 |
| 6.78E-11 | 0.581 | 0.892 | 0.786 | 1.78E-06 | 1 | sc.3 | Ifngr1 |
| 1.30E-10 | 0.561 | 0.949 | 0.841 | 3.41E-06 | 1 | sc.3 | Epsti1 |
| 2.28E-10 | 0.531 | 0.842 | 0.595 | 5.98E-06 | 1 | sc.3 | Gm2682 |
| 4.01E-10 | 0.490 | 0.880 | 0.626 | 1.05E-05 | 1 | sc.3 | Gimap7 |
| 6.25E-10 | 0.528 | 0.570 | 0.298 | 1.64E-05 | 1 | sc.3 | Gzmk |
| 8.08E-10 | 0.440 | 0.506 | 0.260 | 2.12E-05 | 1 | sc.3 | Rtp4 |
| 1.18E-09 | 0.563 | 0.627 | 0.410 | 3.09E-05 | 1 | sc.3 | Slfn1 |
| 1.70E-09 | 0.451 | 0.709 | 0.513 | 4.46E-05 | 1 | sc.3 | Sorl1 |
| 3.38E-09 | 0.437 | 0.728 | 0.516 | 8.85E-05 | 1 | sc.3 | Sp110 |
| 4.12E-09 | 0.436 | 0.905 | 0.797 | 0.000 | 1 | sc.3 | Gimap3 |
| 1.01E-08 | 0.453 | 0.994 | 0.945 | 0.000 | 1 | sc.3 | Ms4a4b |
| 3.58E-08 | 0.709 | 0.772 | 0.591 | 0.001 | 1 | sc.3 | Ly6a |
| 3.98E-08 | 0.461 | 0.753 | 0.538 | 0.001 | 1 | sc.3 | Txnip |
| 6.03E-08 | 0.495 | 0.684 | 0.471 | 0.002 | 1 | sc.3 | Slco3a1 |
| 1.31E-55 | 1.145 | 0.866 | 0.186 | 3.45E-51 | 2 | sc.2 | Cd81 |
| 1.62E-54 | 1.324 | 0.843 | 0.170 | 4.25E-50 | 2 | sc.2 | Sparc |
| 2.40E-49 | 1.119 | 0.827 | 0.180 | 6.29E-45 | 2 | sc.2 | Sparcl1 |
| 1.28E-45 | 1.068 | 0.764 | 0.161 | 3.35E-41 | 2 | sc.2 | Ctsl |
| 1.06E-41 | 1.374 | 0.858 | 0.304 | 2.78E-37 | 2 | sc.2 | Mt1 |
| 6.87E-40 | 1.173 | 0.835 | 0.272 | 1.80E-35 | 2 | sc.2 | Ckb |
| 1.37E-38 | 1.117 | 0.906 | 0.346 | 3.60E-34 | 2 | sc.2 | Qk |
| 2.72E-38 | 1.808 | 0.984 | 0.973 | 7.13E-34 | 2 | sc.2 | mt-Nd1 |
| 5.42E-37 | 1.498 | 1.000 | 1.000 | 1.42E-32 | 2 | sc.2 | mt-Co1 |
| 7.78E-37 | 1.540 | 1.000 | 0.984 | 2.04E-32 | 2 | sc.2 | mt-Nd2 |
| 1.05E-36 | 1.736 | 1.000 | 0.994 | 2.76E-32 | 2 | sc.2 | mt-Nd4 |
| 1.10E-36 | 1.084 | 0.819 | 0.245 | 2.89E-32 | 2 | sc.2 | Cd63 |
| 1.61E-36 | 1.578 | 1.000 | 1.000 | 4.21E-32 | 2 | sc.2 | mt-Atp6 |
| 2.87E-35 | 1.566 | 1.000 | 1.000 | 7.52E-31 | 2 | sc.2 | mt-Cytb |
| 5.45E-35 | 1.099 | 0.874 | 0.336 | 1.43E-30 | 2 | sc.2 | C1qb |
| 6.74E-35 | 1.676 | 1.000 | 1.000 | 1.77E-30 | 2 | sc.2 | mt-Co2 |
| 4.97E-33 | 1.568 | 1.000 | 1.000 | 1.30E-28 | 2 | sc.2 | mt-Co3 |
| 9.81E-33 | 1.282 | 0.984 | 0.682 | 2.57E-28 | 2 | sc.2 | Apoe |
| 1.76E-32 | 1.027 | 0.866 | 0.374 | 4.61E-28 | 2 | sc.2 | C1qa |
| 7.42E-32 | 1.477 | 0.992 | 0.846 | 1.95E-27 | 2 | sc.2 | Ttr |
| 6.28E-31 | 1.314 | 0.992 | 0.811 | 1.65E-26 | 2 | sc.2 | Macf1 |
| 2.72E-26 | 1.213 | 0.992 | 0.954 | 7.14E-22 | 2 | sc.2 | AY036118 |
| 4.96E-21 | 1.076 | 0.984 | 0.893 | 1.30E-16 | 2 | sc.2 | Cst3 |
| 6.15E-21 | 1.237 | 0.969 | 0.667 | 1.61E-16 | 2 | sc.2 | Zbtb20 |
| 1.67E-20 | 1.218 | 0.992 | 0.978 | 4.38E-16 | 2 | sc.2 | Camk1d |
| 4.60E-17 | 1.064 | 0.992 | 0.900 | 1.21E-12 | 2 | sc.2 | Elmo1 |
| 6.89E-17 | 1.055 | 0.969 | 0.775 | 1.81E-12 | 2 | sc.2 | Gphn |
| 4.70E-15 | 1.165 | 0.843 | 0.621 | 1.23E-10 | 2 | sc.2 | Nr3c1 |
| 9.94E-15 | 1.090 | 0.945 | 0.868 | 2.61E-10 | 2 | sc.2 | mt-Nd5 |
| 1.21E-10 | 1.043 | 0.976 | 0.882 | 3.17E-06 | 2 | sc.2 | Xist |
| ##### | 2.066 | 0.886 | 0.043 | 1.15E-109 | 3 | sc.1 | Sell |
| 1.23E-90 | 1.827 | 0.927 | 0.111 | 3.23E-86 | 3 | sc.1 | Cmah |
| 2.59E-89 | 1.253 | 0.740 | 0.035 | 6.78E-85 | 3 | sc.1 | Ccr7 |
| 2.08E-84 | 2.628 | 0.984 | 0.209 | 5.45E-80 | 3 | sc.1 | Lef1 |
| 1.12E-83 | 2.250 | 0.959 | 0.139 | 2.93E-79 | 3 | sc.1 | Klf2 |
| 1.74E-77 | 1.778 | 0.967 | 0.188 | 4.55E-73 | 3 | sc.1 | Tcf7 |
| 1.28E-73 | 2.291 | 0.943 | 0.220 | 3.36E-69 | 3 | sc.1 | Satb1 |
| 1.62E-65 | 1.349 | 0.862 | 0.150 | 4.24E-61 | 3 | sc.1 | S1pr1 |
| 1.66E-65 | 1.829 | 0.935 | 0.241 | 4.35E-61 | 3 | sc.1 | Txk |
| 1.12E-64 | 1.471 | 0.837 | 0.139 | 2.95E-60 | 3 | sc.1 | Klf3 |
| 3.84E-60 | 1.448 | 0.959 | 0.220 | 1.01E-55 | 3 | sc.1 | Aff3 |
| 2.27E-58 | 1.832 | 0.821 | 0.185 | 5.95E-54 | 3 | sc.1 | Fam241a |
| 1.24E-55 | 1.273 | 0.854 | 0.166 | 3.26E-51 | 3 | sc.1 | Slamf6 |
| 1.78E-53 | 1.676 | 0.951 | 0.301 | 4.68E-49 | 3 | sc.1 | Bach2 |
| 1.67E-49 | 1.126 | 1.000 | 1.000 | 4.37E-45 | 3 | sc.1 | Rps29 |
| 5.83E-49 | 1.556 | 0.984 | 0.710 | 1.53E-44 | 3 | sc.1 | Foxp1 |
| 3.10E-47 | 1.844 | 0.927 | 0.391 | 8.12E-43 | 3 | sc.1 | Il7r |
| 2.97E-46 | 1.667 | 0.967 | 0.483 | 7.78E-42 | 3 | sc.1 | Ripor2 |
| 9.54E-46 | 1.450 | 0.870 | 0.231 | 2.50E-41 | 3 | sc.1 | Fgf13 |
| 3.22E-45 | 1.426 | 0.943 | 0.337 | 8.45E-41 | 3 | sc.1 | Emb |
| 7.15E-44 | 1.712 | 0.862 | 0.323 | 1.88E-39 | 3 | sc.1 | Sidt1 |
| 8.78E-44 | 1.197 | 0.919 | 0.421 | 2.30E-39 | 3 | sc.1 | Dgka |
| 5.35E-43 | 1.623 | 0.992 | 0.579 | 1.40E-38 | 3 | sc.1 | Gm2682 |
| 1.00E-41 | 1.081 | 0.992 | 0.994 | 2.62E-37 | 3 | sc.1 | Rps19 |
| 3.64E-40 | 1.329 | 0.764 | 0.236 | 9.55E-36 | 3 | sc.1 | Rras2 |
| 1.96E-38 | 1.346 | 0.902 | 0.380 | 5.14E-34 | 3 | sc.1 | Scml4 |
| 1.56E-34 | 1.231 | 0.959 | 0.642 | 4.09E-30 | 3 | sc.1 | Crlf3 |
| 1.80E-33 | 1.154 | 1.000 | 0.967 | 4.72E-29 | 3 | sc.1 | Arhgap15 |
| 2.79E-31 | 1.080 | 0.959 | 0.616 | 7.32E-27 | 3 | sc.1 | Ssh2 |
| 1.23E-26 | 1.110 | 0.902 | 0.419 | 3.23E-22 | 3 | sc.1 | Gramd3 |
| 4.72E-47 | 3.287 | 0.838 | 0.166 | 1.24E-42 | 4 | sc.4 | Xcl1 |
| 1.73E-34 | 1.333 | 0.865 | 0.236 | 4.53E-30 | 4 | sc.4 | Cd160 |
| 2.83E-17 | 1.037 | 0.878 | 0.476 | 7.41E-13 | 4 | sc.4 | Ybx3 |
| 9.62E-14 | 0.921 | 0.716 | 0.338 | 2.52E-09 | 4 | sc.4 | Bcl2a1d |
| 3.24E-12 | 1.158 | 0.716 | 0.407 | 8.50E-08 | 4 | sc.4 | Il18r1 |
| 1.53E-11 | 0.918 | 0.649 | 0.292 | 4.02E-07 | 4 | sc.4 | Tnfrsf9 |
| 3.20E-11 | 0.729 | 0.635 | 0.276 | 8.40E-07 | 4 | sc.4 | Il18rap |
| 3.24E-11 | 0.584 | 0.689 | 0.338 | 8.49E-07 | 4 | sc.4 | Fam162a |
| 5.44E-11 | 0.800 | 0.649 | 0.294 | 1.43E-06 | 4 | sc.4 | Alcam |
| 6.70E-11 | 0.587 | 0.878 | 0.540 | 1.76E-06 | 4 | sc.4 | Elk3 |
| 1.05E-10 | 0.846 | 0.622 | 0.302 | 2.74E-06 | 4 | sc.4 | Phactr2 |
| 2.15E-10 | 0.645 | 0.986 | 0.809 | 5.63E-06 | 4 | sc.4 | Cd8a |
| 2.55E-10 | 0.964 | 0.959 | 0.758 | 6.69E-06 | 4 | sc.4 | Ifi27l2a |
| 2.85E-10 | 0.746 | 0.824 | 0.496 | 7.48E-06 | 4 | sc.4 | Bcl2a1b |
| 1.11E-09 | 1.041 | 0.797 | 0.499 | 2.90E-05 | 4 | sc.4 | St6galnac3 |
| 1.19E-09 | 0.759 | 0.838 | 0.562 | 3.12E-05 | 4 | sc.4 | Nfat5 |
| 1.22E-09 | 0.627 | 0.757 | 0.417 | 3.21E-05 | 4 | sc.4 | Cd44 |
| 1.53E-09 | 0.667 | 0.838 | 0.512 | 4.01E-05 | 4 | sc.4 | Adamts6 |
| 2.57E-09 | 0.611 | 0.757 | 0.411 | 6.74E-05 | 4 | sc.4 | Lag3 |
| 4.27E-09 | 0.679 | 0.986 | 0.863 | 1.119E-04 | 4 | sc.4 | Rabgap1l |
| 4.79E-09 | 0.704 | 0.932 | 0.675 | 1.255E-04 | 4 | sc.4 | Rbpj |
| 7.63E-09 | 0.589 | 1.000 | 0.993 | 2.001E-04 | 4 | sc.4 | Hsp90ab1 |
| 8.38E-09 | 0.795 | 0.608 | 0.285 | 2.198E-04 | 4 | sc.4 | Rgs16 |
| 9.43E-09 | 0.789 | 0.649 | 0.385 | 2.474E-04 | 4 | sc.4 | Nrp1 |
| 2.13E-08 | 0.595 | 0.959 | 0.812 | 5.592E-04 | 4 | sc.4 | Eif5a |
| 3.04E-08 | 0.581 | 0.824 | 0.592 | 7.972E-04 | 4 | sc.4 | Trps1 |
| 3.80E-08 | 0.684 | 0.986 | 0.802 | 9.962E-04 | 4 | sc.4 | Itgb1 |
| 1.80E-07 | 0.603 | 0.568 | 0.297 | 4.729E-03 | 4 | sc.4 | Gpm6b |
| 2.91E-07 | 0.600 | 0.824 | 0.537 | 7.634E-03 | 4 | sc.4 | Mdfic |
| 7.11E-96 | 3.221 | 1.000 | 0.043 | 1.87E-91 | 5 | sc.6 | Hist1h3c |
| 1.57E-94 | 3.143 | 0.974 | 0.040 | 4.11E-90 | 5 | sc.6 | Hist1h2ab |
| 5.21E-78 | 2.336 | 1.000 | 0.063 | 1.37E-73 | 5 | sc.6 | Birc5 |
| 7.01E-76 | 2.096 | 1.000 | 0.066 | 1.84E-71 | 5 | sc.6 | Cdca8 |
| 1.60E-72 | 2.753 | 1.000 | 0.074 | 4.20E-68 | 5 | sc.6 | Pclaf |
| 3.11E-72 | 2.367 | 0.974 | 0.067 | 8.16E-68 | 5 | sc.6 | Nusap1 |
| 7.96E-69 | 2.104 | 0.921 | 0.064 | 2.09E-64 | 5 | sc.6 | Hist1h3f |
| 7.78E-65 | 2.006 | 0.974 | 0.081 | 2.04E-60 | 5 | sc.6 | Kif11 |
| 4.16E-56 | 2.658 | 0.974 | 0.117 | 1.09E-51 | 5 | sc.6 | Hist1h3e |
| 2.28E-55 | 2.500 | 0.789 | 0.054 | 5.99E-51 | 5 | sc.6 | Cenpf |
| 3.95E-52 | 2.195 | 0.974 | 0.117 | 1.04E-47 | 5 | sc.6 | Prc1 |
| 1.04E-49 | 2.671 | 0.816 | 0.074 | 2.73E-45 | 5 | sc.6 | Ube2c |
| 5.00E-49 | 2.215 | 0.974 | 0.142 | 1.31E-44 | 5 | sc.6 | Hist1h4h |
| 5.70E-49 | 2.465 | 0.974 | 0.148 | 1.50E-44 | 5 | sc.6 | Hist2h2ac |
| 7.13E-48 | 4.304 | 1.000 | 0.169 | 1.87E-43 | 5 | sc.6 | Hist1h1b |
| 5.15E-46 | 2.696 | 1.000 | 0.165 | 1.35E-41 | 5 | sc.6 | Mki67 |
| 1.73E-44 | 4.493 | 1.000 | 0.191 | 4.55E-40 | 5 | sc.6 | Hist1h2ae |
| 2.73E-40 | 1.990 | 1.000 | 0.209 | 7.16E-36 | 5 | sc.6 | Smc2 |
| 1.39E-38 | 2.888 | 1.000 | 0.229 | 3.64E-34 | 5 | sc.6 | Stmn1 |
| 7.12E-37 | 2.137 | 0.947 | 0.185 | 1.87E-32 | 5 | sc.6 | Tubb4b |
| 7.40E-37 | 3.210 | 1.000 | 0.264 | 1.94E-32 | 5 | sc.6 | Top2a |
| 8.30E-37 | 2.115 | 0.921 | 0.183 | 2.18E-32 | 5 | sc.6 | H2afx |
| 6.29E-36 | 4.842 | 1.000 | 0.290 | 1.65E-31 | 5 | sc.6 | Hist1h2ap |
| 1.43E-31 | 2.358 | 1.000 | 0.365 | 3.76E-27 | 5 | sc.6 | Tuba1b |
| 7.01E-29 | 3.416 | 1.000 | 0.497 | 1.84E-24 | 5 | sc.6 | Hist1h4d |
| 6.92E-27 | 2.571 | 1.000 | 0.540 | 1.82E-22 | 5 | sc.6 | Hist1h1e |
| 6.78E-26 | 2.500 | 1.000 | 0.656 | 1.78E-21 | 5 | sc.6 | Tubb5 |
| 3.00E-25 | 3.335 | 1.000 | 0.706 | 7.87E-21 | 5 | sc.6 | Hmgb2 |
| 5.01E-25 | 1.976 | 1.000 | 0.494 | 1.32E-20 | 5 | sc.6 | Ube2s |
| 2.55E-24 | 2.124 | 1.000 | 0.859 | 6.69E-20 | 5 | sc.6 | H2afz |

**Table S4:** CD4+ T cell cluster-defining genes.

| p_val | avg_lo<br>g2FC | pct.1 | pct.2 | p_val_adj | cluster | cluster<br>name | gene |
| --- | --- | --- | --- | --- | --- | --- | --- |
| 2.70E-74 | 2.4658 | 0.972 | 0.382 | 7.09E-70 | 0 | CD4.sc1 | Lef1 |
| 4.65E-70 | 2.2771 | 0.983 | 0.299 | 1.22E-65 | 0 | CD4.sc1 | Klf2 |
| 1.81E-66 | 2.4356 | 0.932 | 0.312 | 4.75E-62 | 0 | CD4.sc1 | Satb1 |
| 2.37E-62 | 1.4292 | 0.864 | 0.151 | 6.21E-58 | 0 | CD4.sc1 | Sell |
| 2.69E-62 | 2.1106 | 1 | 0.497 | 7.04E-58 | 0 | CD4.sc1 | Gm2682 |
| 4.68E-62 | 2.0159 | 0.994 | 0.59 | 1.23E-57 | 0 | CD4.sc1 | Foxp1 |
| 9.41E-62 | 1.8122 | 0.909 | 0.246 | 2.47E-57 | 0 | CD4.sc1 | Txk |
| 1.29E-58 | 1.5158 | 0.938 | 0.281 | 3.39E-54 | 0 | CD4.sc1 | S1pr1 |
| 2.65E-58 | 1.92 | 0.926 | 0.354 | 6.95E-54 | 0 | CD4.sc1 | Bach2 |
| 6.09E-55 | 1.0629 | 1 | 1 | 1.60E-50 | 0 | CD4.sc1 | Rps29 |
| 2.41E-52 | 0.9964 | 0.67 | 0.078 | 6.33E-48 | 0 | CD4.sc1 | Pik3ip1 |
| 2.75E-51 | 1.5112 | 0.926 | 0.374 | 7.20E-47 | 0 | CD4.sc1 | Tcf7 |
| 5.81E-48 | 1.1393 | 0.71 | 0.133 | 1.52E-43 | 0 | CD4.sc1 | Actn1 |
| 9.80E-48 | 1.4496 | 0.972 | 0.698 | 2.57E-43 | 0 | CD4.sc1 | Crlf3 |
| 2.28E-47 | 1.2499 | 0.812 | 0.214 | 5.99E-43 | 0 | CD4.sc1 | Klf3 |
| 5.35E-46 | 1.1001 | 0.591 | 0.058 | 1.40E-41 | 0 | CD4.sc1 | Rflnb |
| 3.98E-45 | 1.4041 | 0.972 | 0.636 | 1.04E-40 | 0 | CD4.sc1 | Ripor2 |
| 1.43E-44 | 1.3212 | 1 | 0.965 | 3.74E-40 | 0 | CD4.sc1 | Arhgap15 |
| 2.40E-44 | 1.6137 | 0.506 | 0.03 | 6.29E-40 | 0 | CD4.sc1 | Igfbp4 |
| 8.62E-44 | 1.143 | 0.545 | 0.048 | 2.26E-39 | 0 | CD4.sc1 | St8sia6 |
| 6.04E-43 | 1.0328 | 0.71 | 0.136 | 1.58E-38 | 0 | CD4.sc1 | Ccr7 |
| 1.23E-42 | 1.1636 | 0.932 | 0.497 | 3.23E-38 | 0 | CD4.sc1 | Dgka |
| 1.35E-40 | 1.2443 | 0.858 | 0.364 | 3.55E-36 | 0 | CD4.sc1 | Scml4 |
| 9.21E-40 | 1.3161 | 0.778 | 0.249 | 2.42E-35 | 0 | CD4.sc1 | Cmah |
| 6.09E-35 | 1.2343 | 0.835 | 0.397 | 1.60E-30 | 0 | CD4.sc1 | Tmem71 |
| 1.56E-34 | 1.1863 | 0.955 | 0.678 | 4.09E-30 | 0 | CD4.sc1 | Prkcq |
| 8.82E-34 | 1.0426 | 0.977 | 0.907 | 2.31E-29 | 0 | CD4.sc1 | Rapgef6 |
| 1.68E-33 | 1.1313 | 0.761 | 0.266 | 4.41E-29 | 0 | CD4.sc1 | St6gal1 |
| 2.80E-27 | 1.0327 | 0.943 | 0.751 | 7.34E-23 | 0 | CD4.sc1 | Ikzf1 |
| 1.48E-26 | 1.0186 | 0.688 | 0.271 | 3.88E-22 | 0 | CD4.sc1 | Ugcg |
| 1.45E-69 | 2.6831 | 0.981 | 0.296 | 3.79E-65 | 1 | CD4.sc5 | Ikzf2 |
| 6.35E-65 | 1.5118 | 0.786 | 0.077 | 1.66E-60 | 1 | CD4.sc5 | Foxp3 |
| 6.43E-53 | 1.8706 | 0.78 | 0.14 | 1.69E-48 | 1 | CD4.sc5 | Tnfrsf9 |
| 6.99E-47 | 1.6338 | 0.761 | 0.154 | 1.83E-42 | 1 | CD4.sc5 | Il2ra |
| 2.77E-46 | 1.833 | 0.95 | 0.361 | 7.26E-42 | 1 | CD4.sc5 | Tnfrsf4 |
| 1.39E-45 | 1.5504 | 0.969 | 0.513 | 3.65E-41 | 1 | CD4.sc5 | Tnfrsf18 |
| 3.04E-44 | 1.208 | 0.629 | 0.087 | 7.97E-40 | 1 | CD4.sc5 | Neb |
| 1.90E-38 | 1.6722 | 0.962 | 0.448 | 5.00E-34 | 1 | CD4.sc5 | Ctla4 |
| 1.32E-35 | 1.4683 | 0.78 | 0.318 | 3.46E-31 | 1 | CD4.sc5 | Arl5a |
| 4.44E-35 | 1.3279 | 0.824 | 0.34 | 1.16E-30 | 1 | CD4.sc5 | Izumo1r |
| 2.70E-32 | 1.0323 | 0.547 | 0.089 | 7.08E-28 | 1 | CD4.sc5 | Itgb8 |
| 3.08E-32 | 1.1847 | 0.874 | 0.414 | 8.09E-28 | 1 | CD4.sc5 | Capg |
| 7.62E-31 | 1.0991 | 0.792 | 0.318 | 2.00E-26 | 1 | CD4.sc5 | Tnfrsf1b |
| 4.00E-30 | 1.0045 | 0.906 | 0.407 | 1.05E-25 | 1 | CD4.sc5 | Il2rb |
| 7.21E-29 | 1.3823 | 0.824 | 0.402 | 1.89E-24 | 1 | CD4.sc5 | Itgav |
| 1.83E-28 | 1.1439 | 0.855 | 0.508 | 4.79E-24 | 1 | CD4.sc5 | Gimap7 |
| 1.02E-27 | 1.2783 | 0.736 | 0.323 | 2.67E-23 | 1 | CD4.sc5 | Ighm |
| 4.78E-27 | 1.3767 | 0.975 | 0.848 | 1.25E-22 | 1 | CD4.sc5 | Ifi27l2a |
| 1.40E-26 | 0.9938 | 0.906 | 0.571 | 3.66E-22 | 1 | CD4.sc5 | Pim1 |
| 2.49E-23 | 1.0318 | 0.881 | 0.525 | 6.53E-19 | 1 | CD4.sc5 | Samsn1 |
| 3.30E-21 | 1.1678 | 0.717 | 0.337 | 8.66E-17 | 1 | CD4.sc5 | Hopx |
| 2.02E-20 | 0.9867 | 0.811 | 0.431 | 5.29E-16 | 1 | CD4.sc5 | Maf |
| 6.45E-20 | 1.0151 | 0.969 | 0.798 | 1.69E-15 | 1 | CD4.sc5 | Rabgap1l |
| 8.10E-19 | 1.0454 | 0.912 | 0.713 | 2.12E-14 | 1 | CD4.sc5 | Sdf4 |
| 1.59E-18 | 1.1913 | 0.818 | 0.453 | 4.17E-14 | 1 | CD4.sc5 | S100a6 |
| 1.14E-17 | 0.9536 | 0.566 | 0.214 | 3.00E-13 | 1 | CD4.sc5 | ligp1 |
| 1.60E-17 | 0.9873 | 0.748 | 0.373 | 4.21E-13 | 1 | CD4.sc5 | Odc1 |
| 1.91E-15 | 0.9527 | 0.893 | 0.672 | 5.01E-11 | 1 | CD4.sc5 | Ly6a |
| 2.28E-15 | 0.9349 | 0.881 | 0.646 | 5.98E-11 | 1 | CD4.sc5 | Icos |
| 2.97E-12 | 0.9491 | 0.836 | 0.677 | 7.80E-08 | 1 | CD4.sc5 | Vim |
| 1.04E-13 | 0.69 | 0.672 | 0.345 | 2.73E-09 | 2 | CD4.sc2 | Arhgap26 |
| 6.88E-09 | 0.5019 | 0.513 | 0.262 | 0.00018 | 2 | CD4.sc2 | Cd63 |
| 1.06E-07 | 0.6665 | 0.807 | 0.582 | 0.00277 | 2 | CD4.sc2 | Itgb1 |
| 2.40E-07 | 0.7834 | 0.756 | 0.582 | 0.006293 | 2 | CD4.sc2 | Emb |
| 4.60E-07 | 0.635 | 0.697 | 0.519 | 0.012059 | 2 | CD4.sc2 | Pde7a |
| 6.15E-07 | 0.4858 | 0.958 | 0.879 | 0.016137 | 2 | CD4.sc2 | Celf2 |
| 2.18E-06 | 0.6497 | 0.723 | 0.569 | 0.05709 | 2 | CD4.sc2 | Btg2 |
| 3.53E-06 | 0.7988 | 0.605 | 0.378 | 0.092528 | 2 | CD4.sc2 | C1qb |
| 4.44E-42 | 2.1955 | 0.965 | 0.203 | 1.16E-37 | 3 | CD4.sc3 | Cxcr6 |
| 6.45E-30 | 2.1697 | 0.965 | 0.311 | 1.69E-25 | 3 | CD4.sc3 | Nkg7 |
| 3.56E-25 | 3.0164 | 0.895 | 0.331 | 9.35E-21 | 3 | CD4.sc3 | Ccl5 |
| 8.47E-24 | 1.6638 | 0.544 | 0.085 | 2.22E-19 | 3 | CD4.sc3 | Ifng |
| 1.25E-21 | 1.1158 | 0.579 | 0.104 | 3.29E-17 | 3 | CD4.sc3 | Tmem163 |
| 1.65E-21 | 1.5962 | 0.912 | 0.427 | 4.33E-17 | 3 | CD4.sc3 | Id2 |
| 9.99E-21 | 1.3405 | 0.86 | 0.294 | 2.62E-16 | 3 | CD4.sc3 | Pdcd1 |
| 2.02E-18 | 1.081 | 1 | 0.954 | 5.31E-14 | 3 | CD4.sc3 | Cd3g |
| 8.21E-18 | 1.2564 | 0.93 | 0.538 | 2.15E-13 | 3 | CD4.sc3 | Rbpj |
| 5.91E-17 | 1.6558 | 0.877 | 0.414 | 1.55E-12 | 3 | CD4.sc3 | S100a4 |
| 6.78E-17 | 1.3834 | 1 | 0.814 | 1.78E-12 | 3 | CD4.sc3 | AW112010 |
| 2.36E-15 | 1.0615 | 0.825 | 0.346 | 6.20E-11 | 3 | CD4.sc3 | Hip1 |
| 2.96E-15 | 1.3437 | 0.772 | 0.304 | 7.78E-11 | 3 | CD4.sc3 | Dusp2 |
| 9.25E-15 | 0.8408 | 0.684 | 0.244 | 2.43E-10 | 3 | CD4.sc3 | Tnfsf10 |
| 1.25E-14 | 1.9371 | 0.509 | 0.13 | 3.27E-10 | 3 | CD4.sc3 | Ccl4 |
| 4.34E-14 | 0.9849 | 0.561 | 0.159 | 1.14E-09 | 3 | CD4.sc3 | Rgs16 |
| 4.95E-14 | 1.3471 | 0.86 | 0.453 | 1.30E-09 | 3 | CD4.sc3 | Rgs1 |
| 7.77E-14 | 1.032 | 0.947 | 0.578 | 2.04E-09 | 3 | CD4.sc3 | AU020206 |
| 6.53E-13 | 1.2485 | 0.947 | 0.511 | 1.71E-08 | 3 | CD4.sc3 | S100a6 |
| 1.62E-12 | 0.9067 | 1 | 0.662 | 4.26E-08 | 3 | CD4.sc3 | S100a11 |
| 5.45E-12 | 0.8826 | 1 | 0.973 | 1.43E-07 | 3 | CD4.sc3 | Cd52 |
| 6.76E-12 | 0.9882 | 0.684 | 0.273 | 1.77E-07 | 3 | CD4.sc3 | Bhlhe40 |
| 1.43E-11 | 0.972 | 0.982 | 0.677 | 3.75E-07 | 3 | CD4.sc3 | Lgals1 |
| 7.31E-11 | 0.7638 | 0.877 | 0.507 | 1.92E-06 | 3 | CD4.sc3 | Gng2 |
| 6.32E-10 | 0.7972 | 0.544 | 0.193 | 1.66E-05 | 3 | CD4.sc3 | Lgals3 |
| 7.11E-10 | 0.9605 | 0.807 | 0.472 | 1.86E-05 | 3 | CD4.sc3 | Cd226 |
| 7.43E-10 | 0.7606 | 1 | 0.901 | 1.95E-05 | 3 | CD4.sc3 | Arl6ip1 |
| 7.47E-08 | 1.1071 | 0.649 | 0.338 | 0.001958 | 3 | CD4.sc3 | Ctla2a |
| 1.15E-07 | 0.7851 | 0.965 | 0.865 | 0.003019 | 3 | CD4.sc3 | Sub1 |
| 1.62E-07 | 0.7853 | 0.649 | 0.354 | 0.004261 | 3 | CD4.sc3 | Ctsw |
| 6.60E-39 | 1.395 | 0.775 | 0.079 | 1.73E-34 | 4 | CD4.sc4 | Gm49180 |
| 1.69E-29 | 2.5873 | 0.95 | 0.257 | 4.44E-25 | 4 | CD4.sc4 | Tnfsf8 |
| 1.13E-25 | 1.6995 | 0.8 | 0.157 | 2.96E-21 | 4 | CD4.sc4 | Tnfsf11 |
| 7.07E-20 | 1.4357 | 0.875 | 0.272 | 1.85E-15 | 4 | CD4.sc4 | Bhlhe40 |
| 7.14E-20 | 1.9977 | 0.925 | 0.378 | 1.87E-15 | 4 | CD4.sc4 | Eea1 |
| 1.70E-17 | 1.7789 | 0.975 | 0.515 | 4.47E-13 | 4 | CD4.sc4 | Trps1 |
| 2.27E-17 | 1.5468 | 0.825 | 0.303 | 5.96E-13 | 4 | CD4.sc4 | Tbc1d4 |
| 5.60E-17 | 1.6436 | 0.95 | 0.446 | 1.47E-12 | 4 | CD4.sc4 | Gm15283 |
| 6.28E-17 | 1.3698 | 0.75 | 0.2 | 1.65E-12 | 4 | CD4.sc4 | Mir155hg |
| 1.03E-16 | 1.4134 | 0.825 | 0.281 | 2.71E-12 | 4 | CD4.sc4 | Phactr2 |
| 9.60E-16 | 1.5897 | 0.95 | 0.566 | 2.52E-11 | 4 | CD4.sc4 | Arap2 |
| 4.14E-15 | 1.5608 | 0.95 | 0.691 | 1.09E-10 | 4 | CD4.sc4 | Hif1a |
| 1.06E-13 | 1.1862 | 0.8 | 0.264 | 2.77E-09 | 4 | CD4.sc4 | Kcnq5 |
| 1.57E-13 | 1.1136 | 0.675 | 0.202 | 4.12E-09 | 4 | CD4.sc4 | Cd200 |
| 4.69E-13 | 1.1912 | 0.925 | 0.537 | 1.23E-08 | 4 | CD4.sc4 | Itpr1 |
| 5.07E-13 | 1.3477 | 0.9 | 0.446 | 1.33E-08 | 4 | CD4.sc4 | Odc1 |
| 7.10E-13 | 1.1081 | 0.925 | 0.545 | 1.86E-08 | 4 | CD4.sc4 | Jak2 |
| 7.71E-13 | 1.2137 | 0.95 | 0.624 | 2.02E-08 | 4 | CD4.sc4 | Ostf1 |
| 2.00E-12 | 1.0762 | 0.9 | 0.478 | 5.24E-08 | 4 | CD4.sc4 | Etv6 |
| 2.17E-12 | 1.2973 | 0.975 | 0.547 | 5.69E-08 | 4 | CD4.sc4 | Rbpj |
| 2.60E-12 | 1.1922 | 0.875 | 0.459 | 6.82E-08 | 4 | CD4.sc4 | Cd82 |
| 5.42E-12 | 1.2641 | 0.95 | 0.755 | 1.42E-07 | 4 | CD4.sc4 | Sdf4 |
| 1.69E-11 | 1.0617 | 0.95 | 0.605 | 4.42E-07 | 4 | CD4.sc4 | Itgb1 |
| 1.73E-11 | 1.288 | 0.625 | 0.221 | 4.54E-07 | 4 | CD4.sc4 | Gpm6b |
| 2.00E-11 | 1.0825 | 0.925 | 0.479 | 5.24E-07 | 4 | CD4.sc4 | Nfat5 |
| 1.60E-10 | 1.195 | 0.9 | 0.513 | 4.19E-06 | 4 | CD4.sc4 | Slamf6 |
| 1.80E-10 | 1.3157 | 0.975 | 0.717 | 4.71E-06 | 4 | CD4.sc4 | Stat4 |
| 2.92E-10 | 1.1008 | 0.775 | 0.375 | 7.67E-06 | 4 | CD4.sc4 | Kdm2b |
| 7.90E-09 | 1.1523 | 0.95 | 0.764 | 0.000207 | 4 | CD4.sc4 | Prkca |
| 3.18E-07 | 1.1287 | 0.675 | 0.363 | 0.008349 | 4 | CD4.sc4 | Nrp1 |
| 1.43E-74 | 2.8862 | 1 | 0.031 | 3.76E-70 | 5 | CD4.sc6 | Hist1h2ab |
| 6.56E-68 | 2.311 | 1 | 0.038 | 1.72E-63 | 5 | CD4.sc6 | Nusap1 |
| 5.05E-63 | 1.9574 | 0.957 | 0.038 | 1.33E-58 | 5 | CD4.sc6 | Tpx2 |
| 5.27E-61 | 2.8896 | 1 | 0.049 | 1.38E-56 | 5 | CD4.sc6 | Hist1h3c |
| 3.61E-60 | 2.0618 | 1 | 0.049 | 9.48E-56 | 5 | CD4.sc6 | Kif11 |
| 1.30E-53 | 2.0623 | 0.87 | 0.038 | 3.41E-49 | 5 | CD4.sc6 | Cenpf |
| 4.48E-49 | 2.1381 | 0.957 | 0.062 | 1.17E-44 | 5 | CD4.sc6 | Cdca8 |
| 2.30E-47 | 2.8141 | 1 | 0.08 | 6.02E-43 | 5 | CD4.sc6 | Pclaf |
| 2.46E-45 | 2.1941 | 0.913 | 0.064 | 6.45E-41 | 5 | CD4.sc6 | Birc5 |
| 3.76E-43 | 2.3881 | 0.913 | 0.067 | 9.87E-39 | 5 | CD4.sc6 | Ube2c |
| 1.27E-40 | 2.1334 | 1 | 0.098 | 3.34E-36 | 5 | CD4.sc6 | Hist1h4h |
| 4.34E-40 | 2.2946 | 0.957 | 0.089 | 1.14E-35 | 5 | CD4.sc6 | Prc1 |
| 8.40E-38 | 2.1106 | 1 | 0.111 | 2.20E-33 | 5 | CD4.sc6 | Diaph3 |
| 1.16E-37 | 4.2177 | 1 | 0.122 | 3.05E-33 | 5 | CD4.sc6 | Hist1h1b |
| 2.68E-35 | 2.3752 | 0.957 | 0.116 | 7.03E-31 | 5 | CD4.sc6 | Hist1h3e |
| 4.67E-34 | 3.9954 | 1 | 0.142 | 1.22E-29 | 5 | CD4.sc6 | Hist1h2ae |
| 5.19E-34 | 3.1412 | 1 | 0.138 | 1.36E-29 | 5 | CD4.sc6 | Mki67 |
| 6.56E-28 | 2.124 | 0.957 | 0.16 | 1.72E-23 | 5 | CD4.sc6 | Hist2h2ac |
| 2.16E-27 | 3.3427 | 1 | 0.205 | 5.66E-23 | 5 | CD4.sc6 | Top2a |
| 2.22E-26 | 2.0812 | 1 | 0.212 | 5.81E-22 | 5 | CD4.sc6 | Smc2 |
| 2.63E-25 | 1.9485 | 0.957 | 0.16 | 6.91E-21 | 5 | CD4.sc6 | Rad51b |
| 1.17E-24 | 4.4356 | 1 | 0.247 | 3.06E-20 | 5 | CD4.sc6 | Hist1h2ap |
| 1.38E-23 | 2.9391 | 1 | 0.26 | 3.62E-19 | 5 | CD4.sc6 | Stmn1 |
| 1.53E-19 | 2.034 | 1 | 0.365 | 4.01E-15 | 5 | CD4.sc6 | Lmnbl |
| 8.97E-19 | 2.045 | 1 | 0.408 | 2.35E-14 | 5 | CD4.sc6 | Tuba1b |
| 2.74E-18 | 3.2597 | 1 | 0.497 | 7.18E-14 | 5 | CD4.sc6 | Hist1h4d |
| 8.10E-17 | 2.4069 | 1 | 0.675 | 2.13E-12 | 5 | CD4.sc6 | Tubb5 |
| 8.39E-16 | 3.1063 | 1 | 0.793 | 2.20E-11 | 5 | CD4.sc6 | Hmgb2 |
| 1.23E-15 | 2.6079 | 1 | 0.833 | 3.23E-11 | 5 | CD4.sc6 | H2afz |
| 3.83E-15 | 2.2363 | 1 | 0.519 | 1.01E-10 | 5 | CD4.sc6 | Hist1h1e |

**Table S5:**
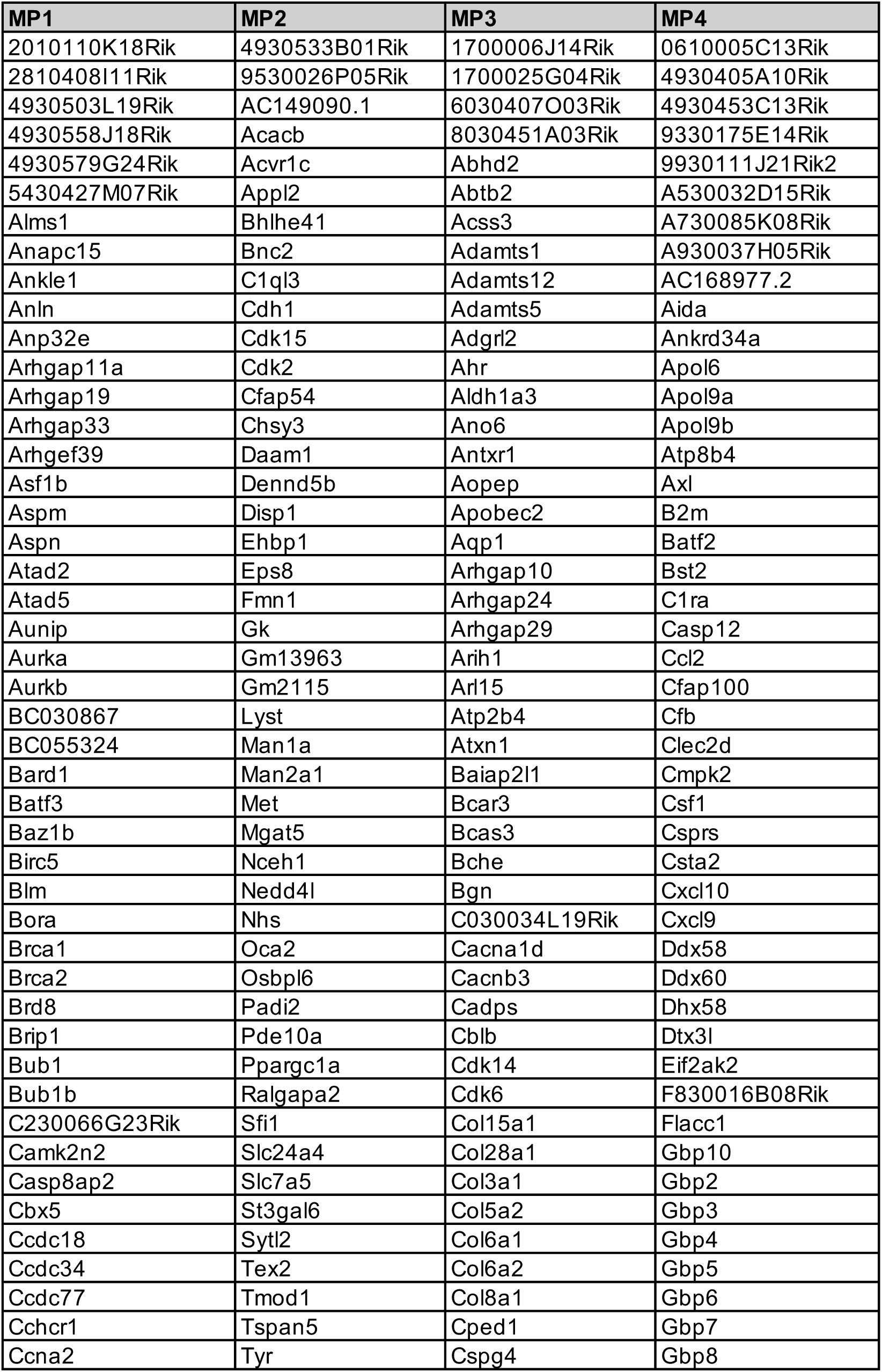
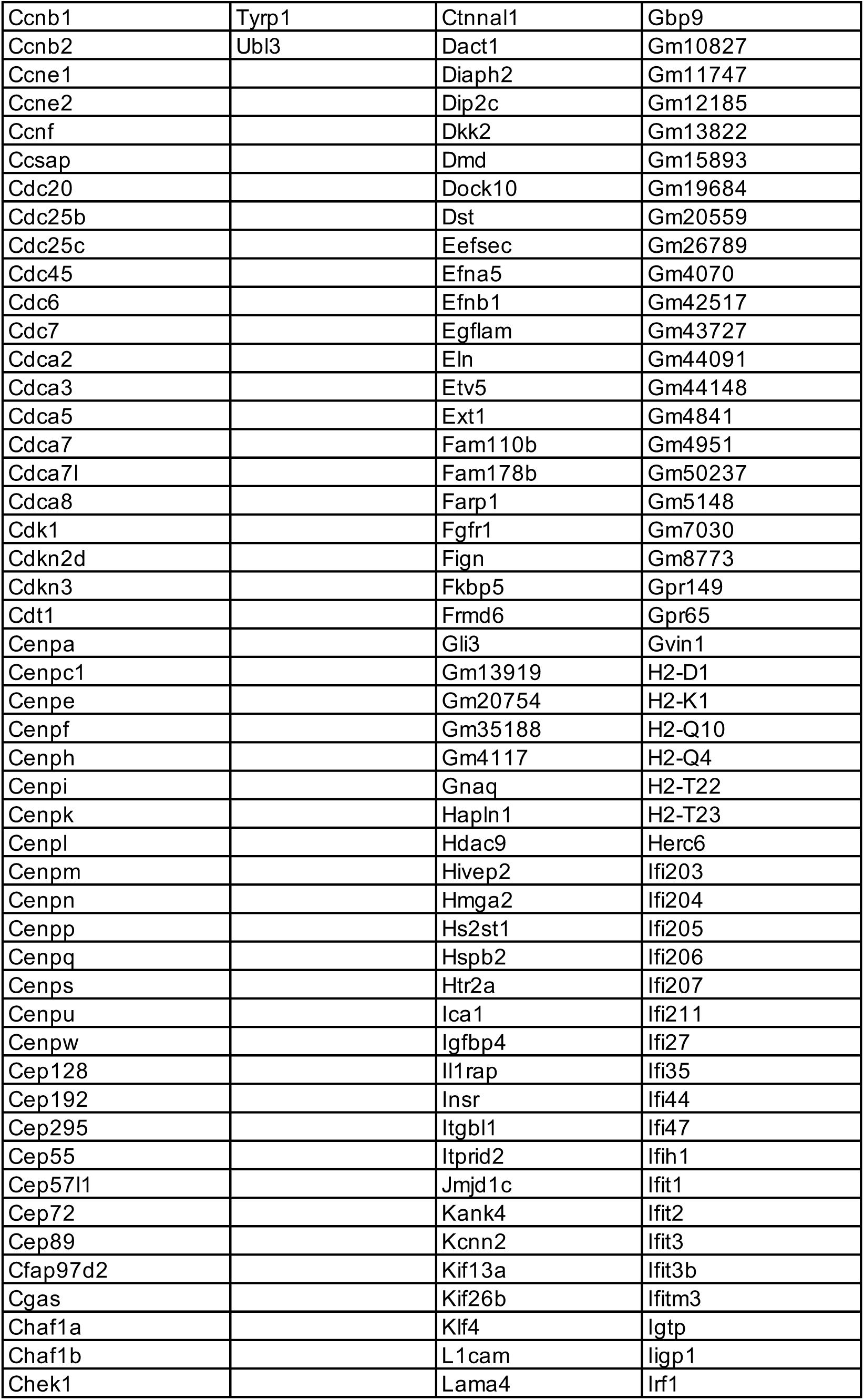
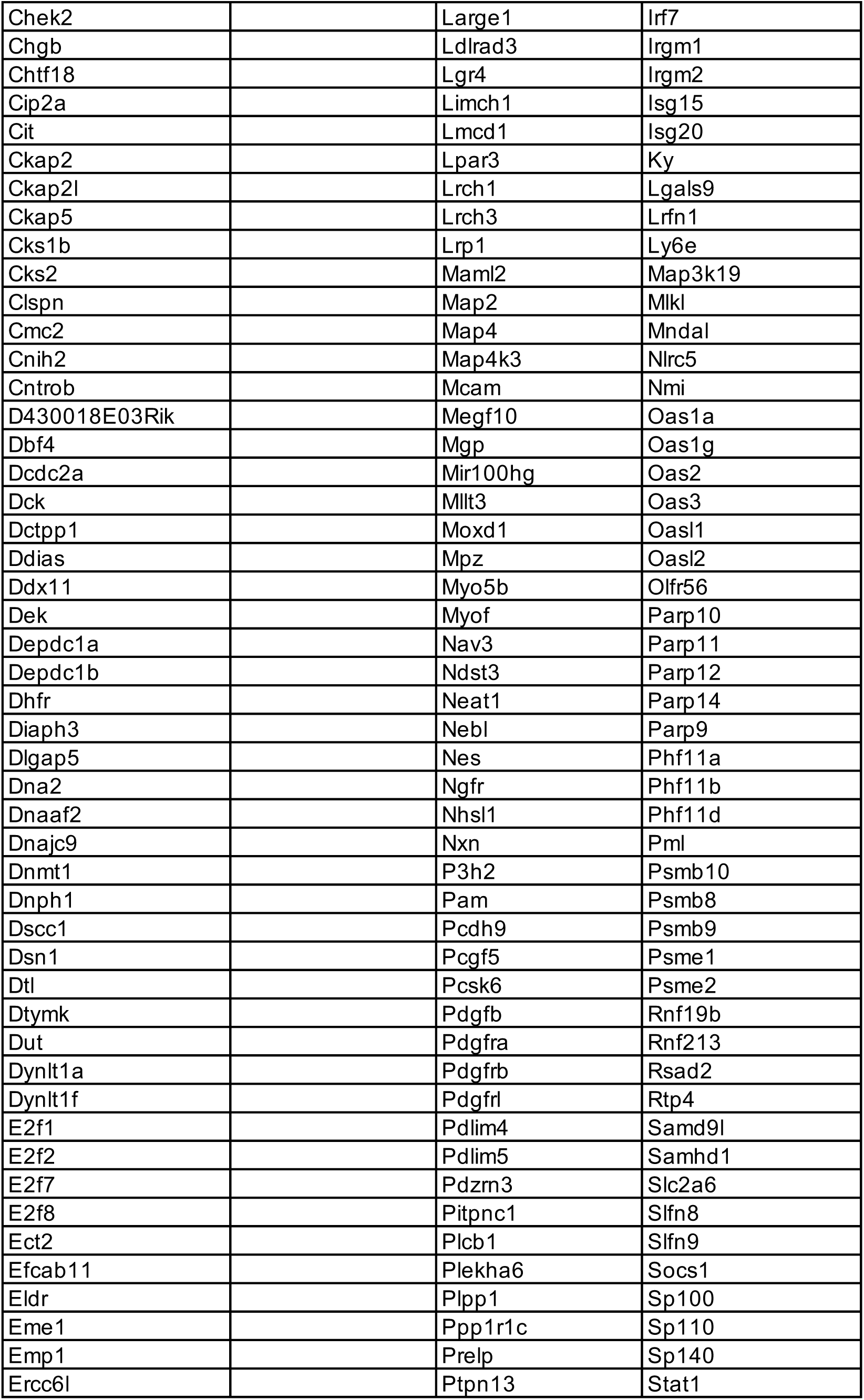
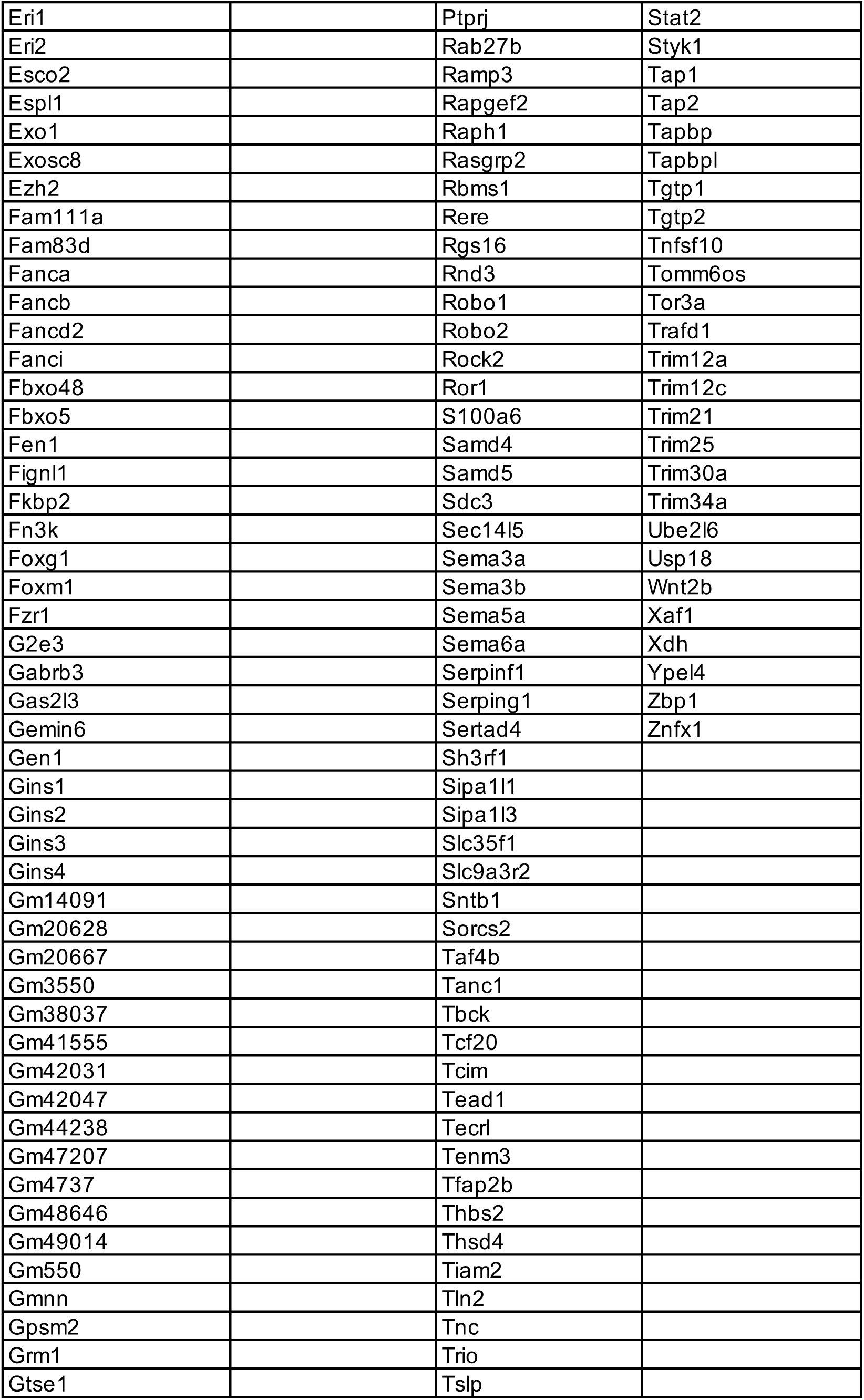
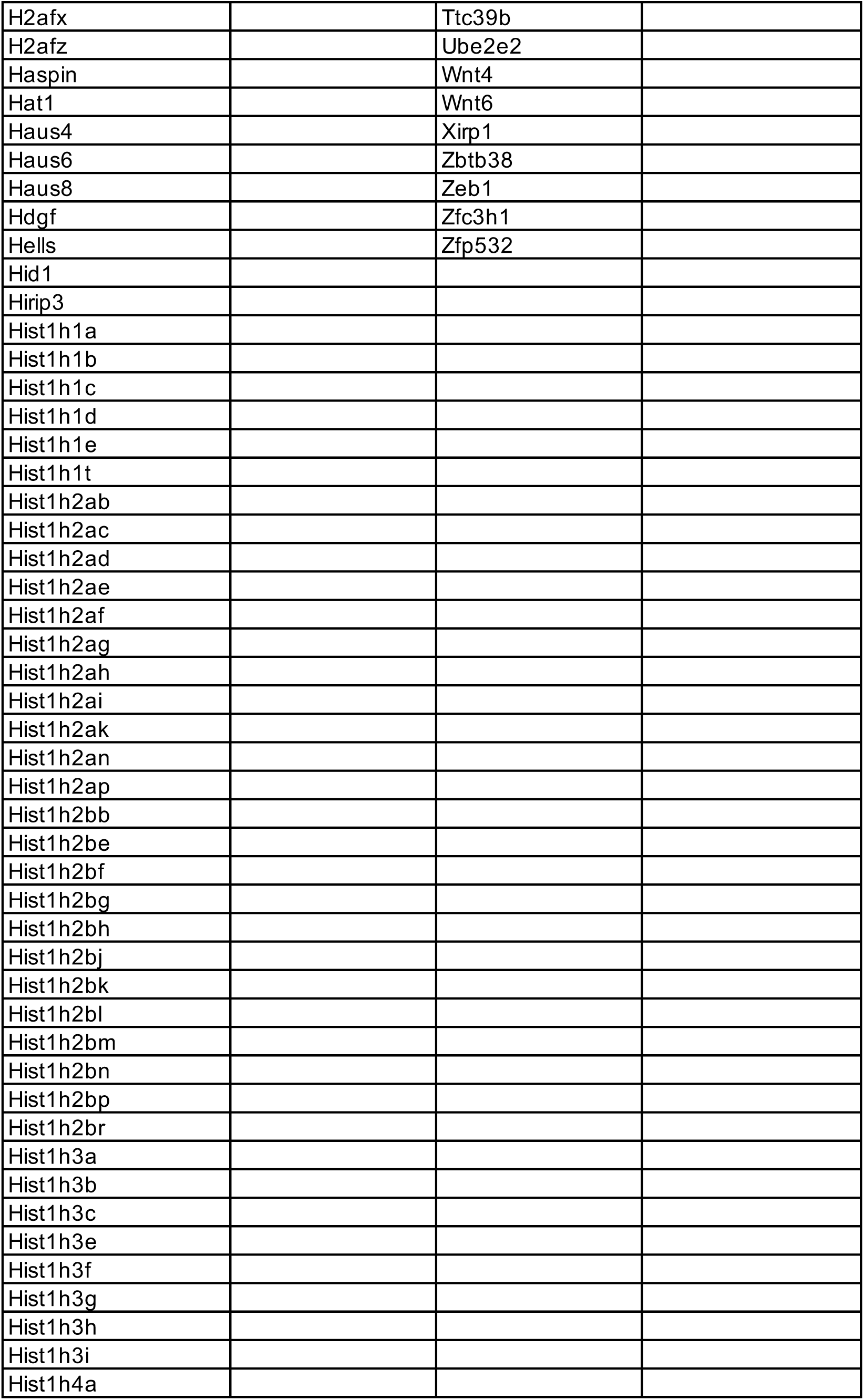
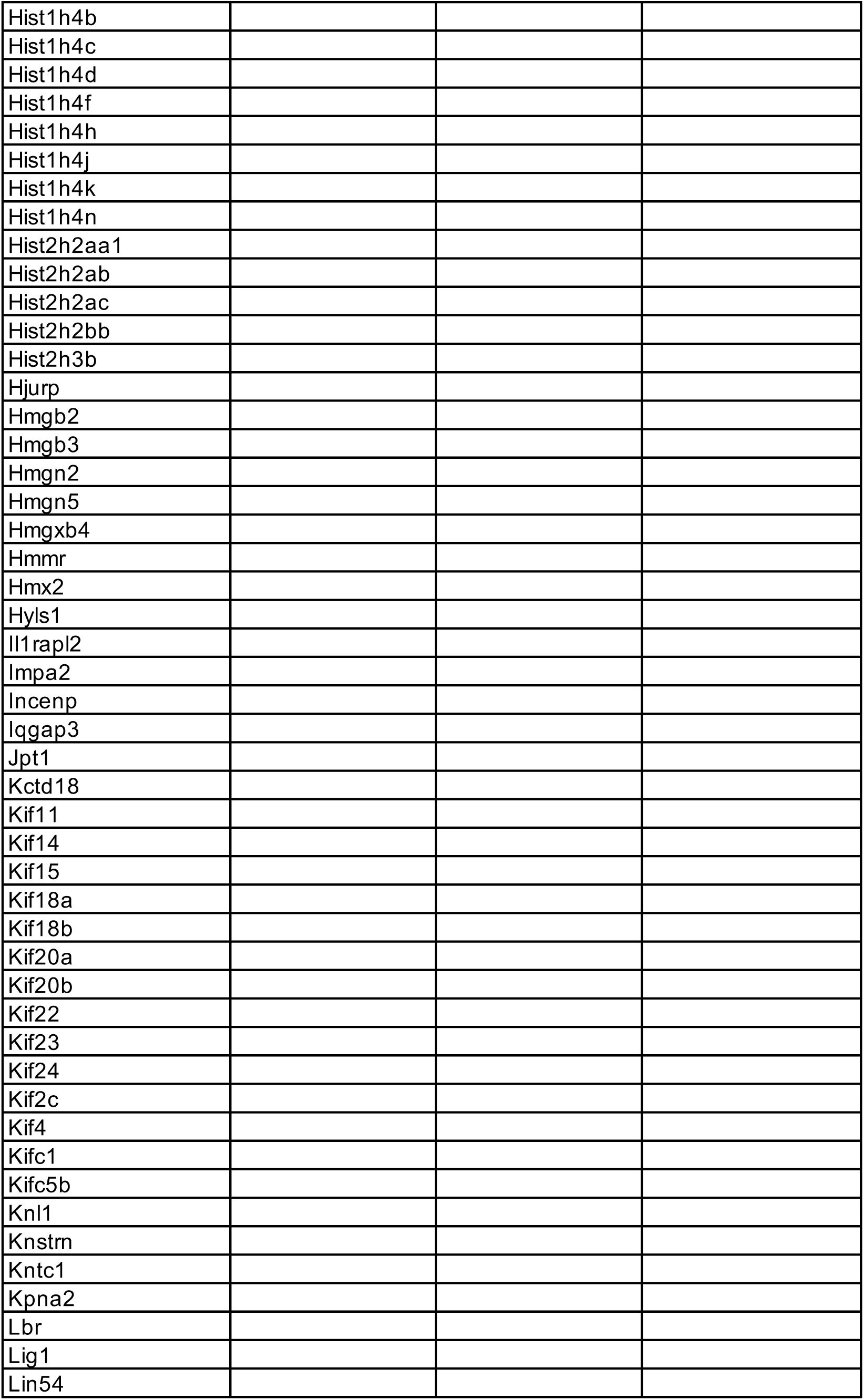
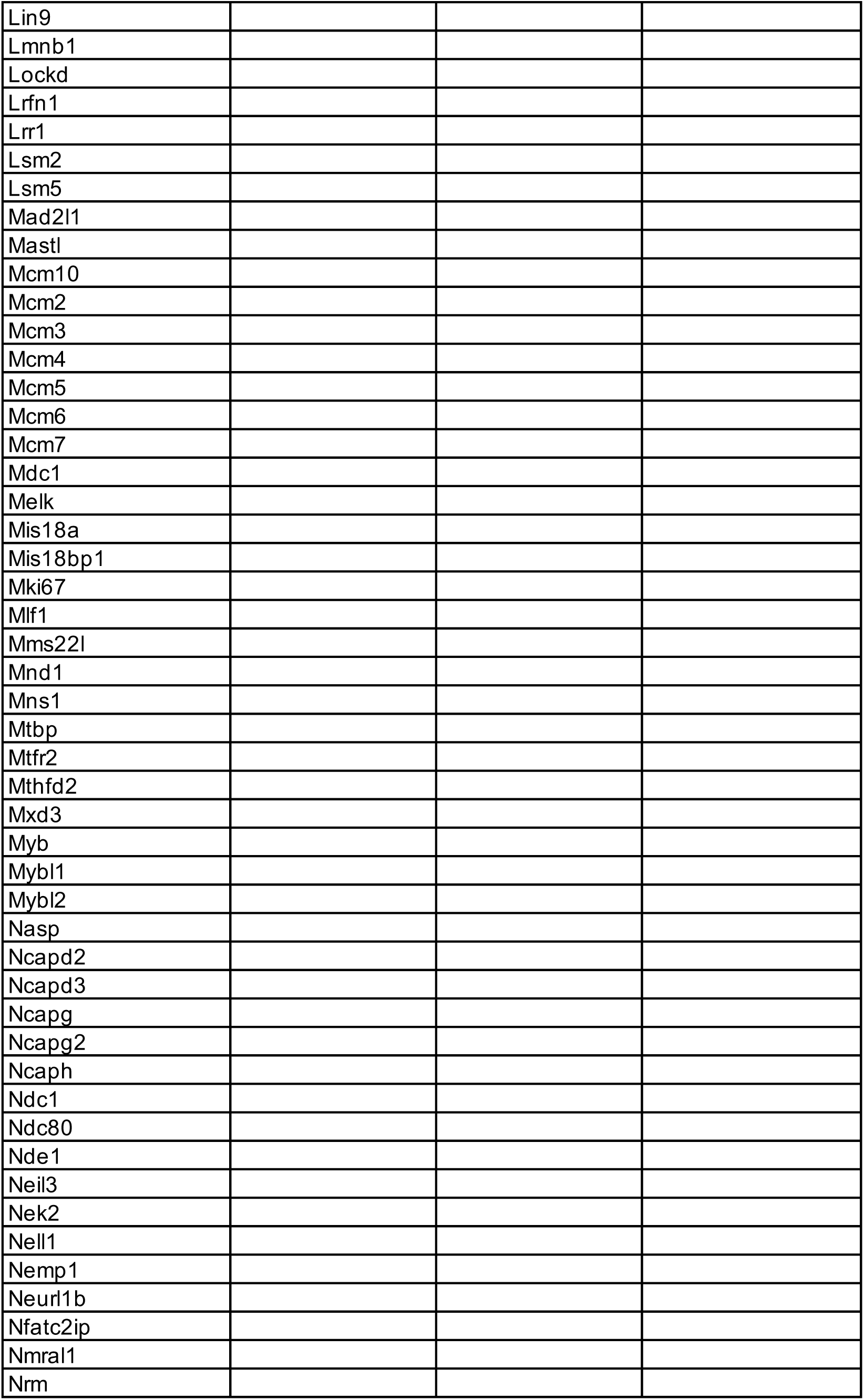
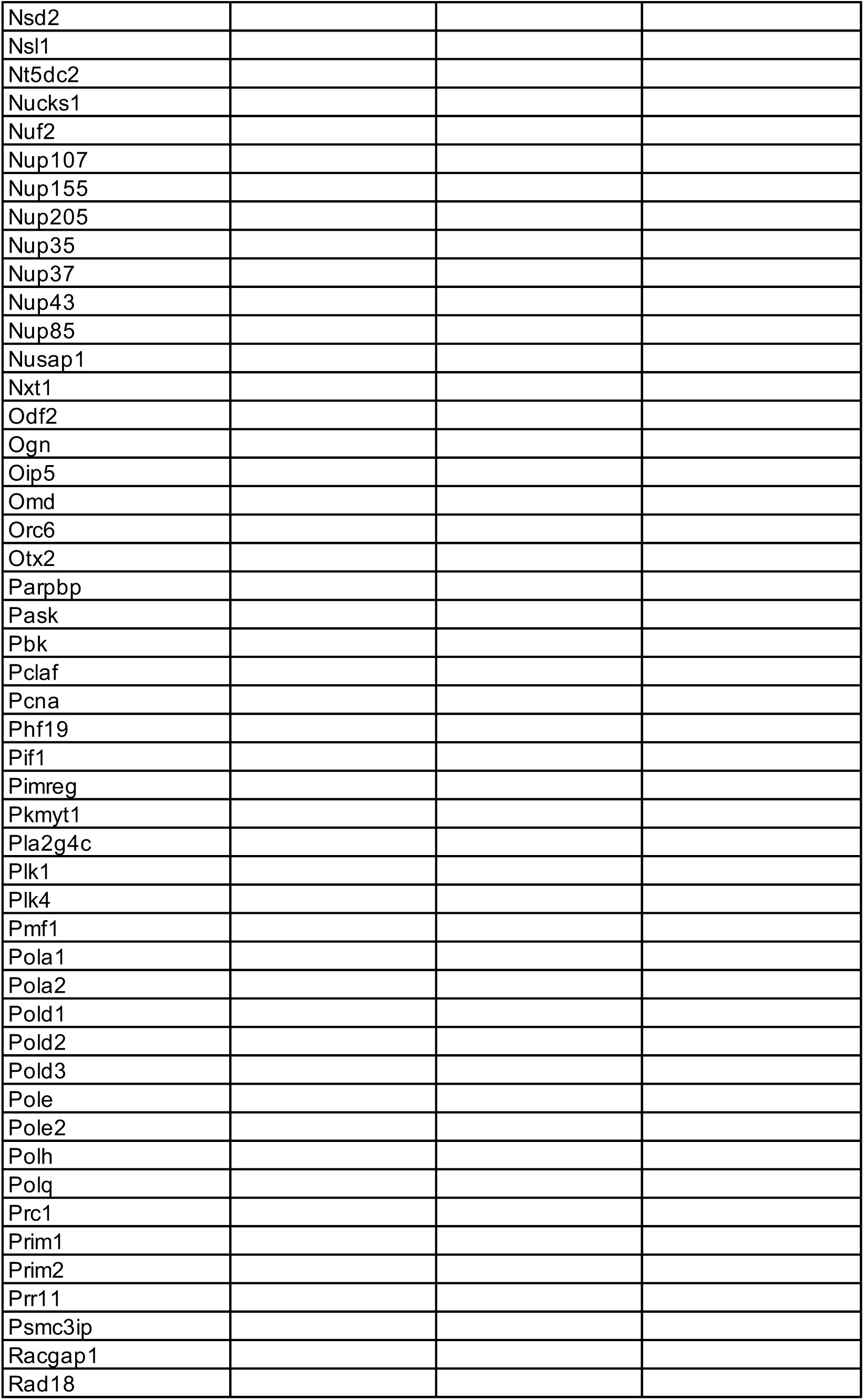
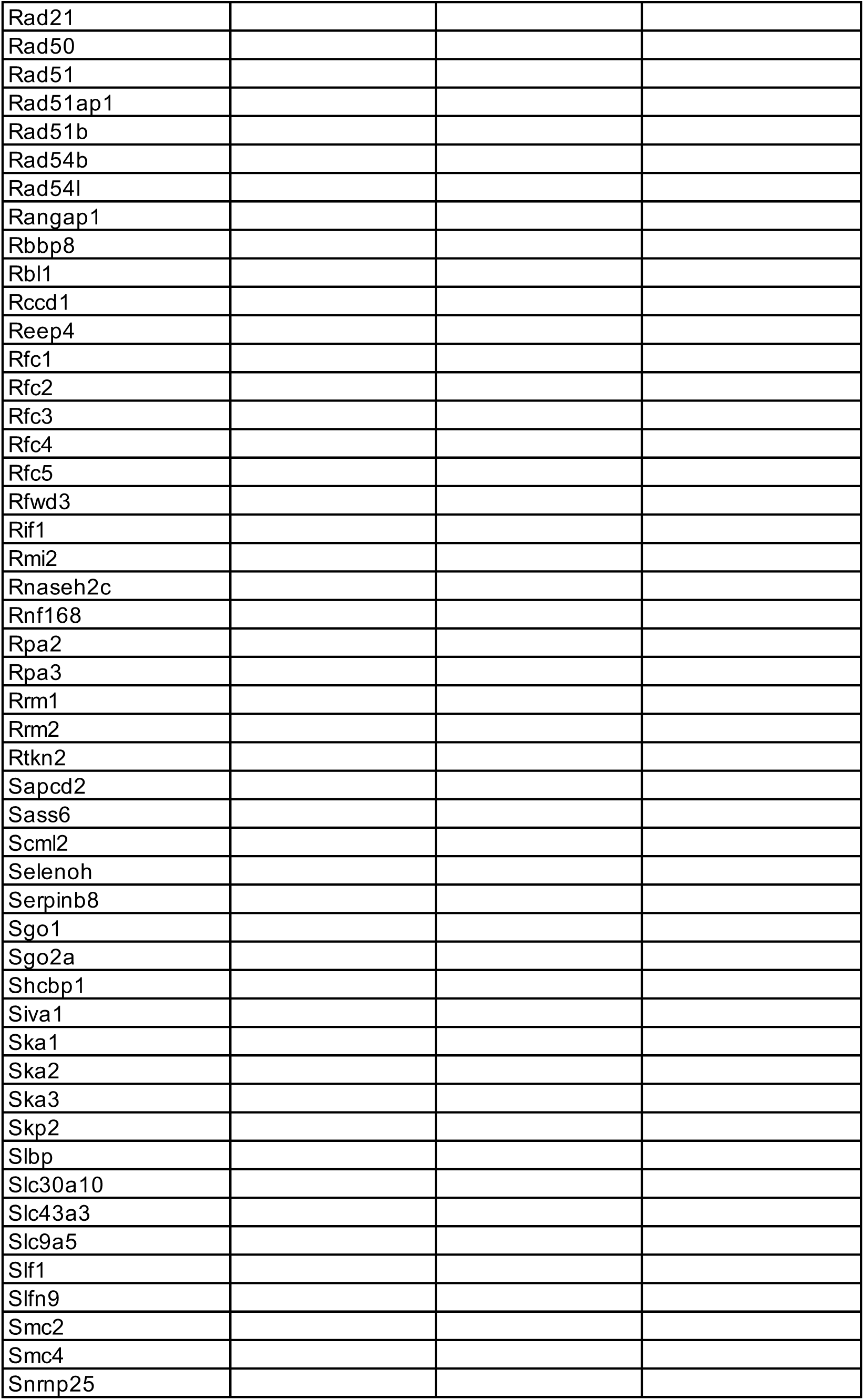
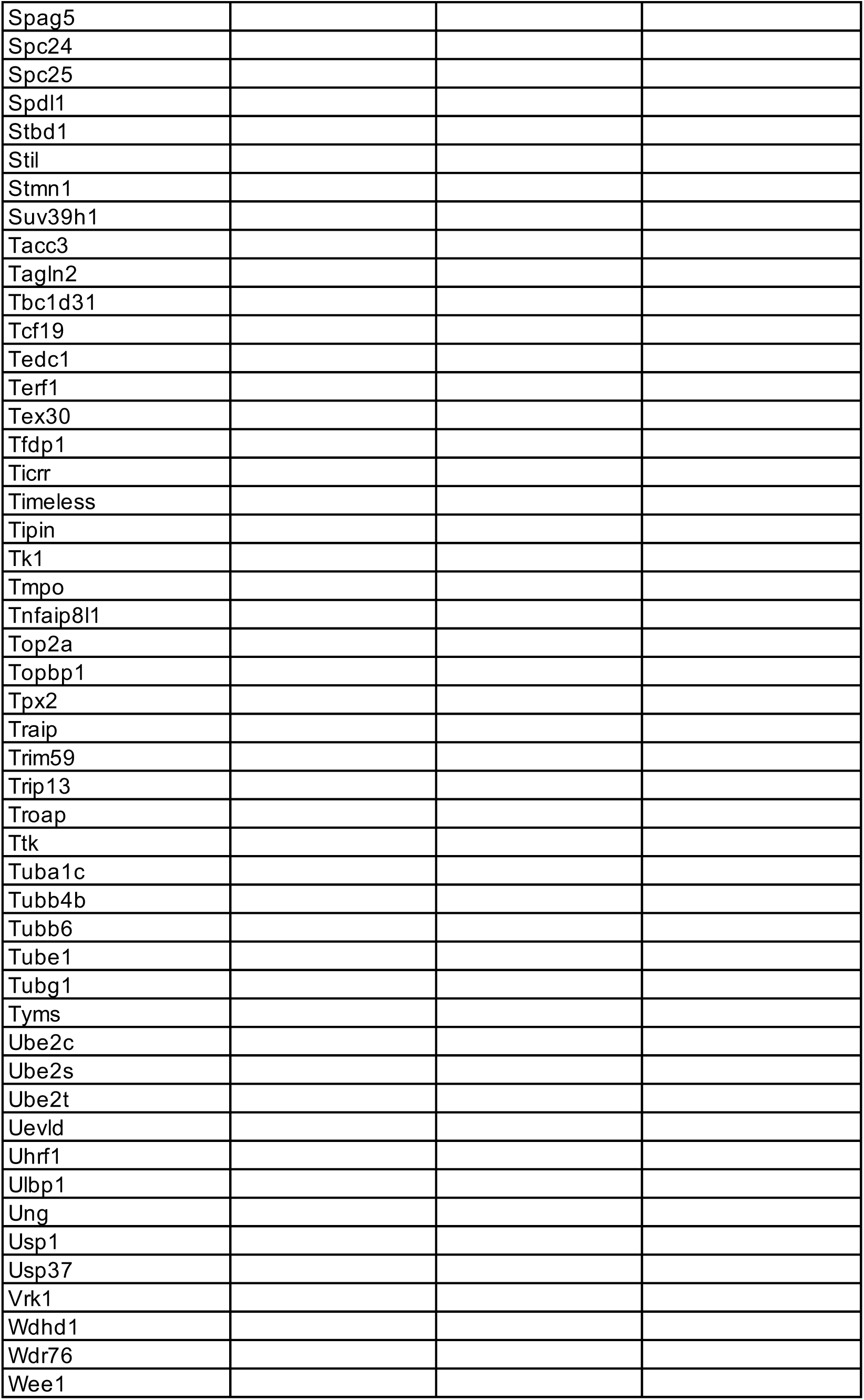
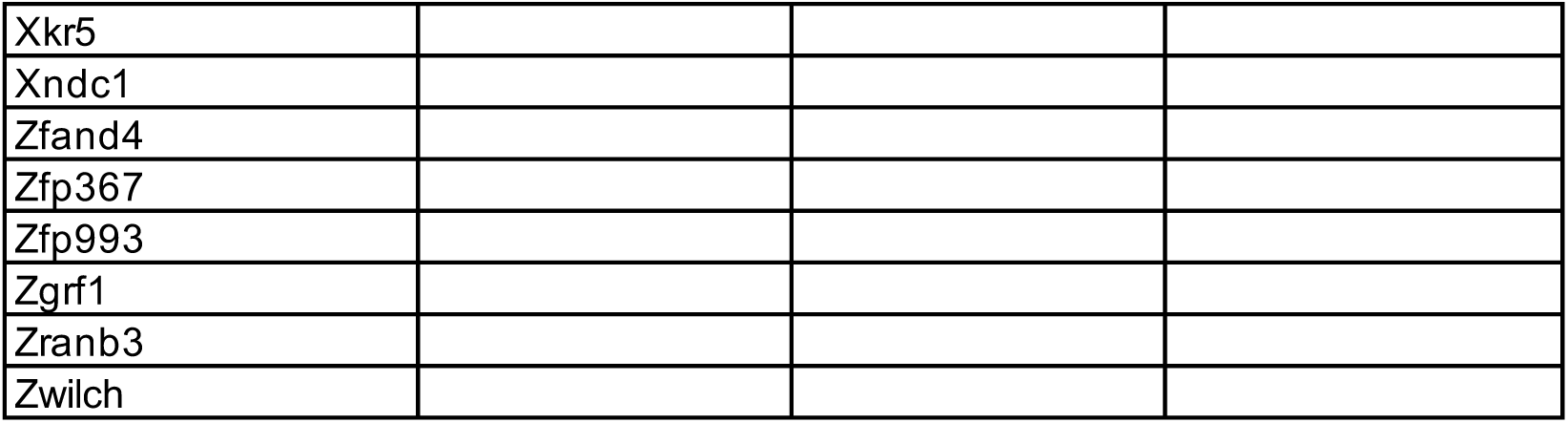
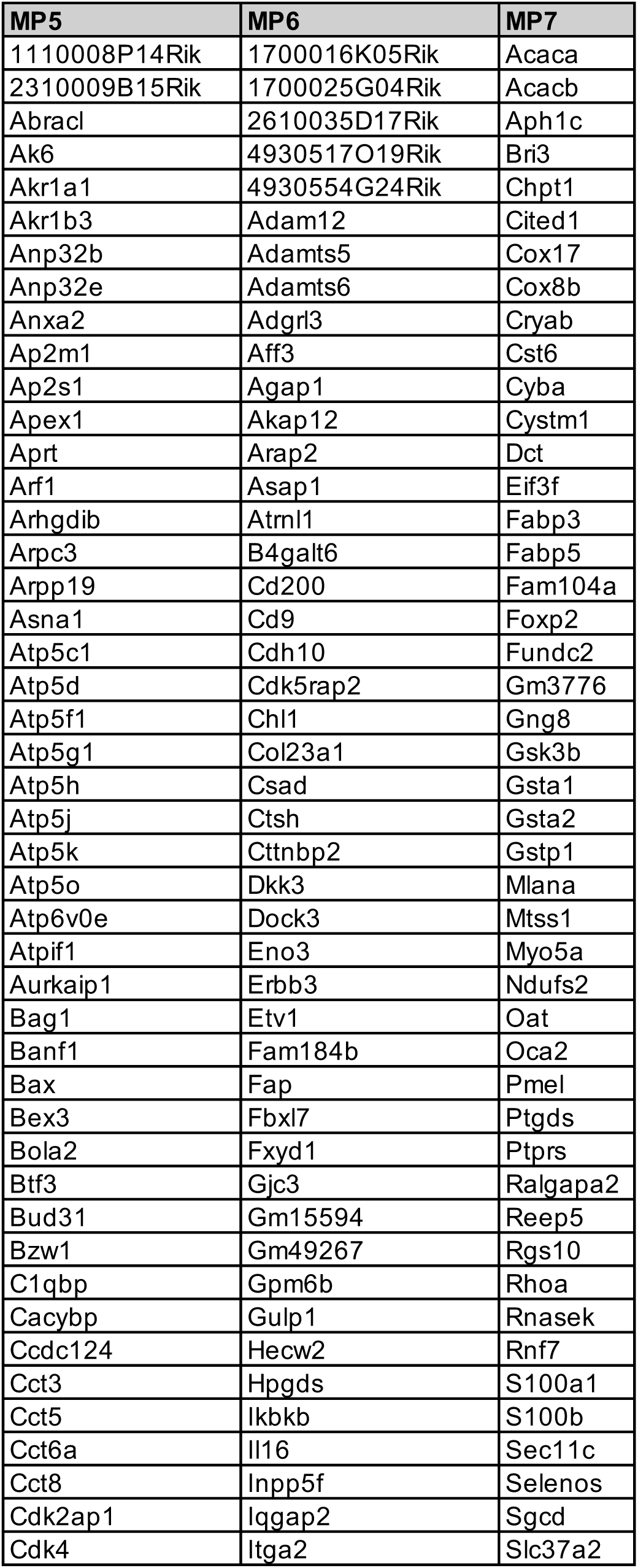
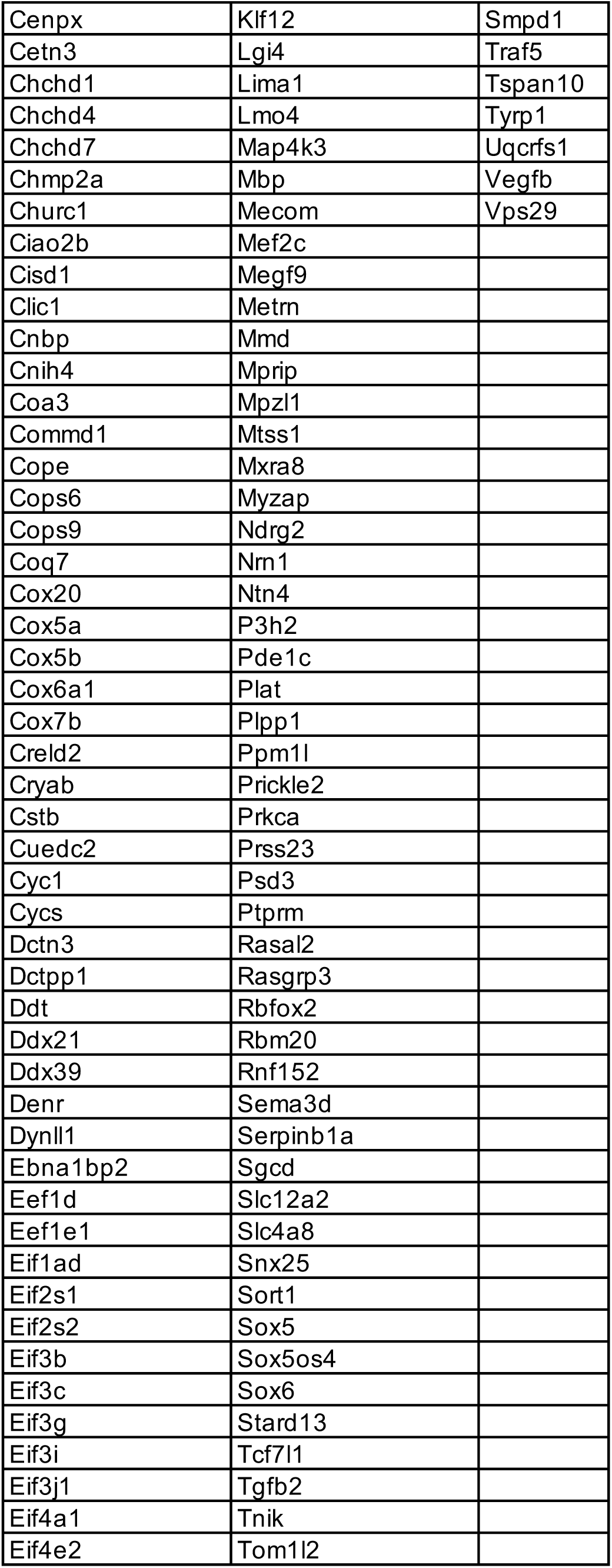
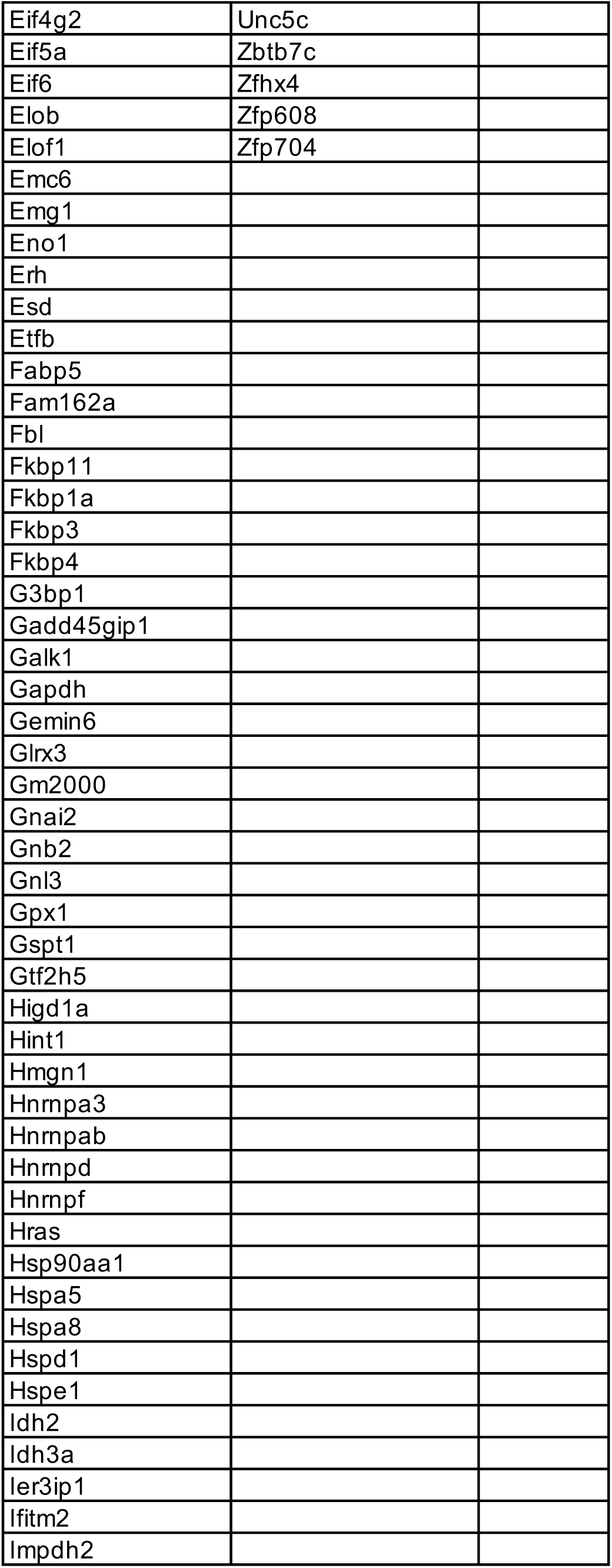
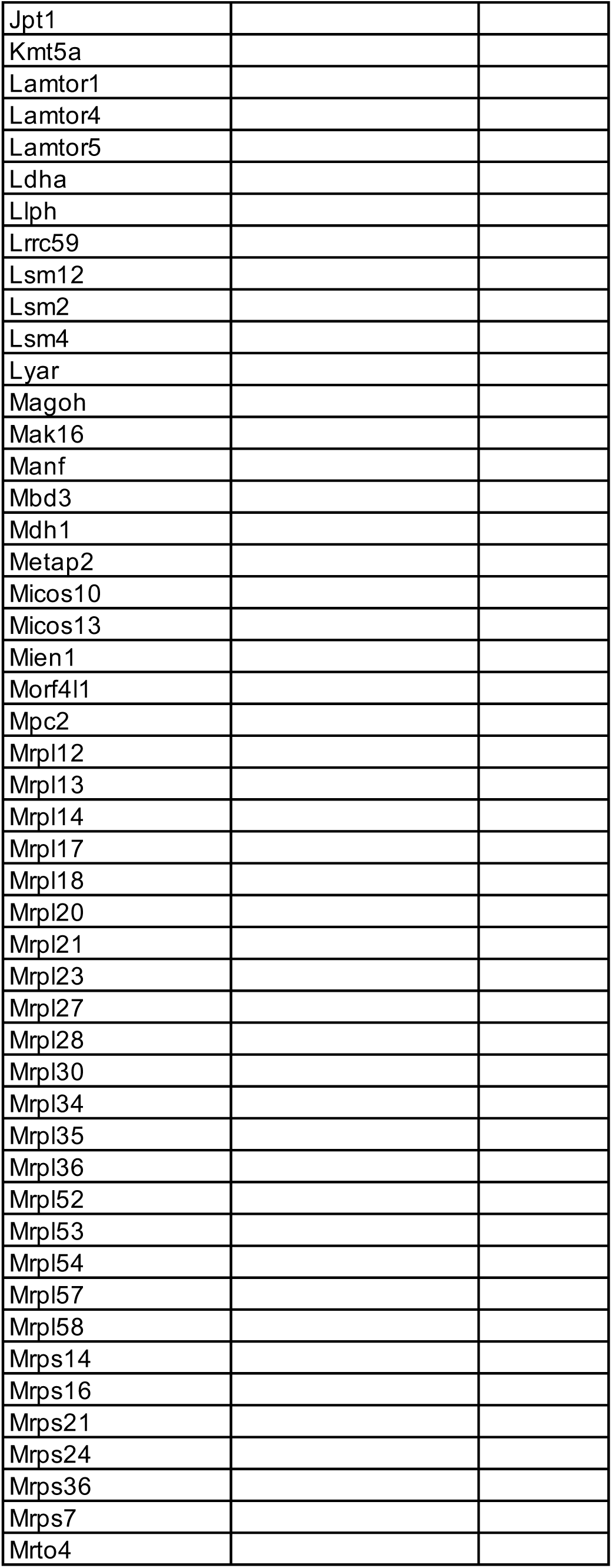
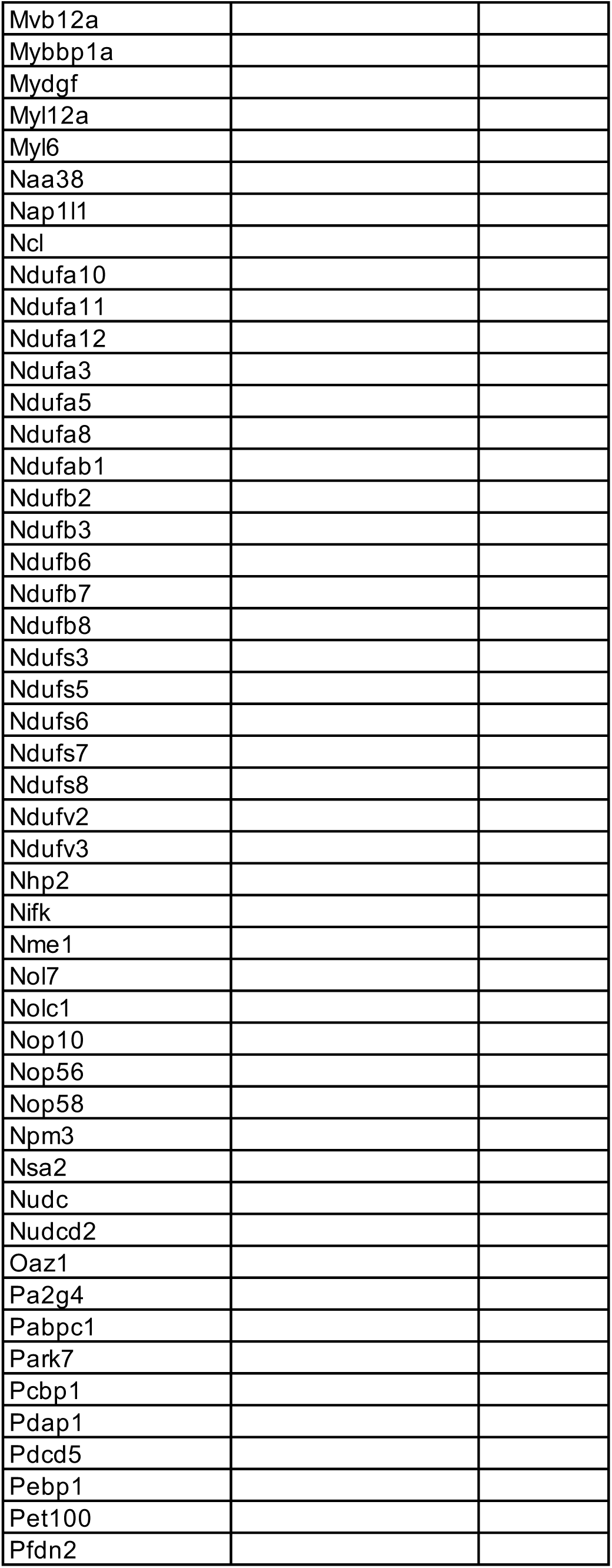
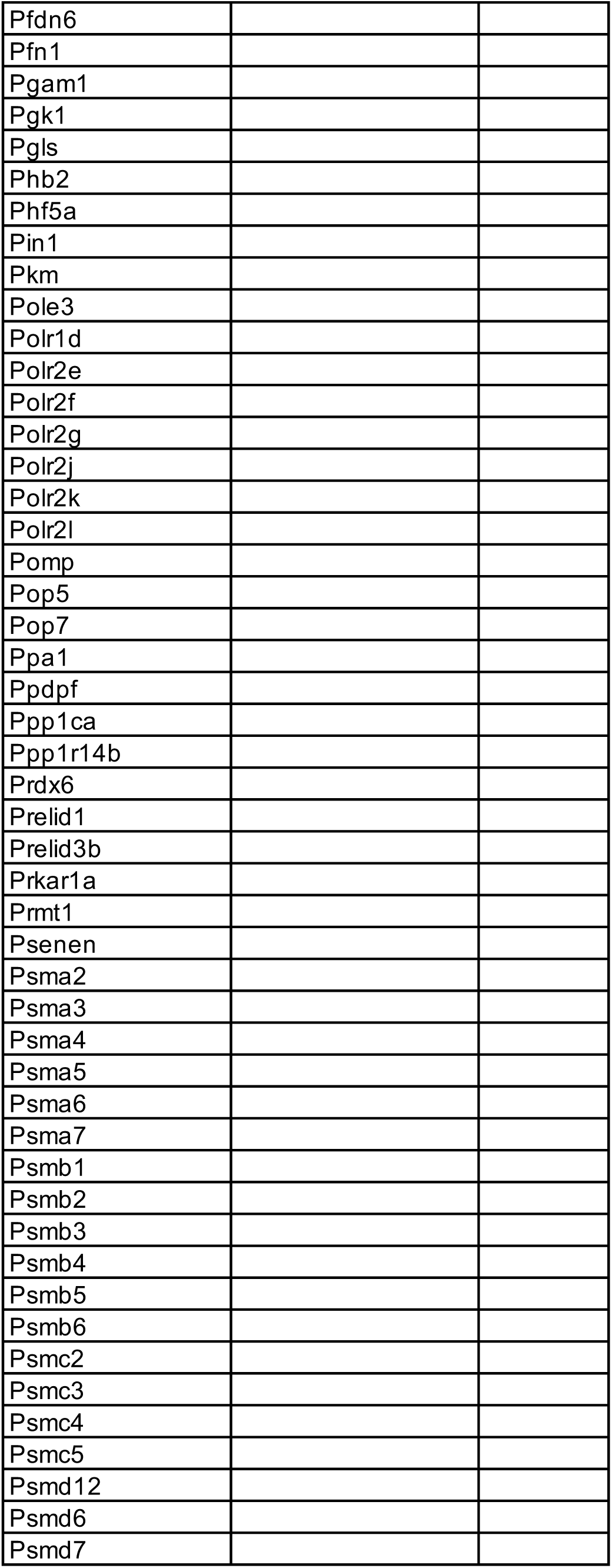
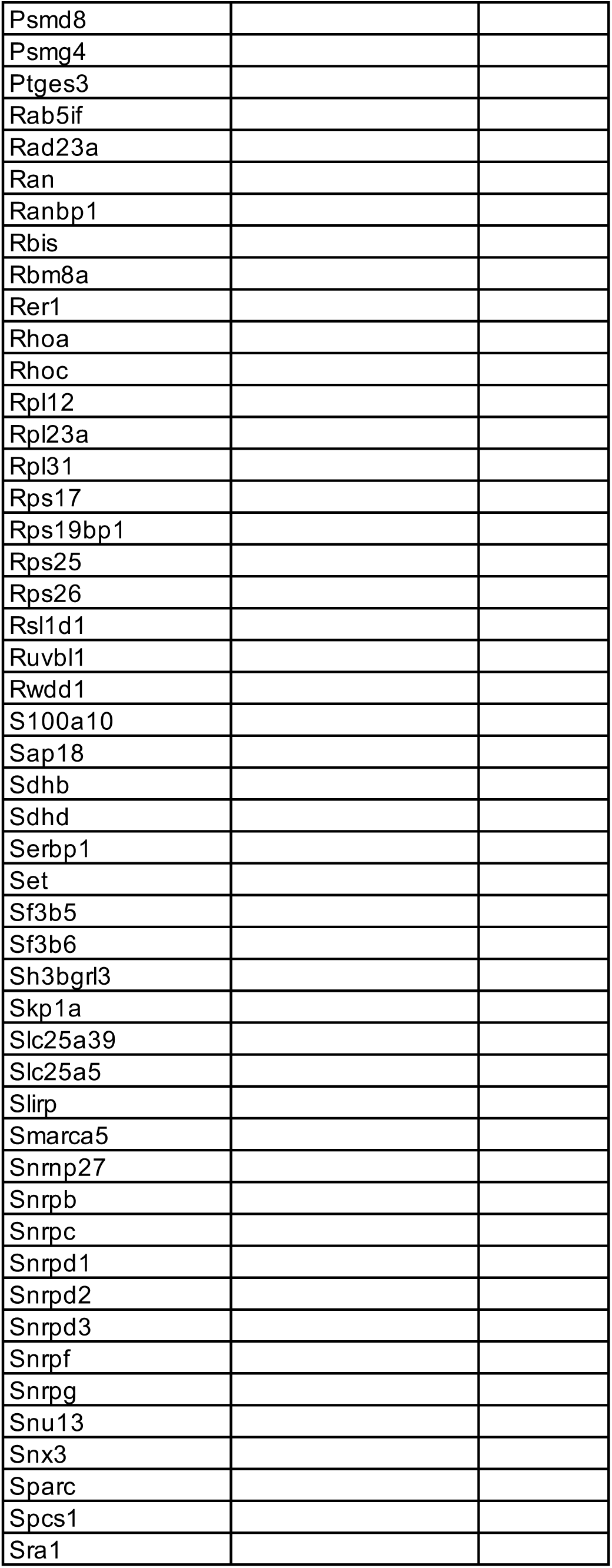
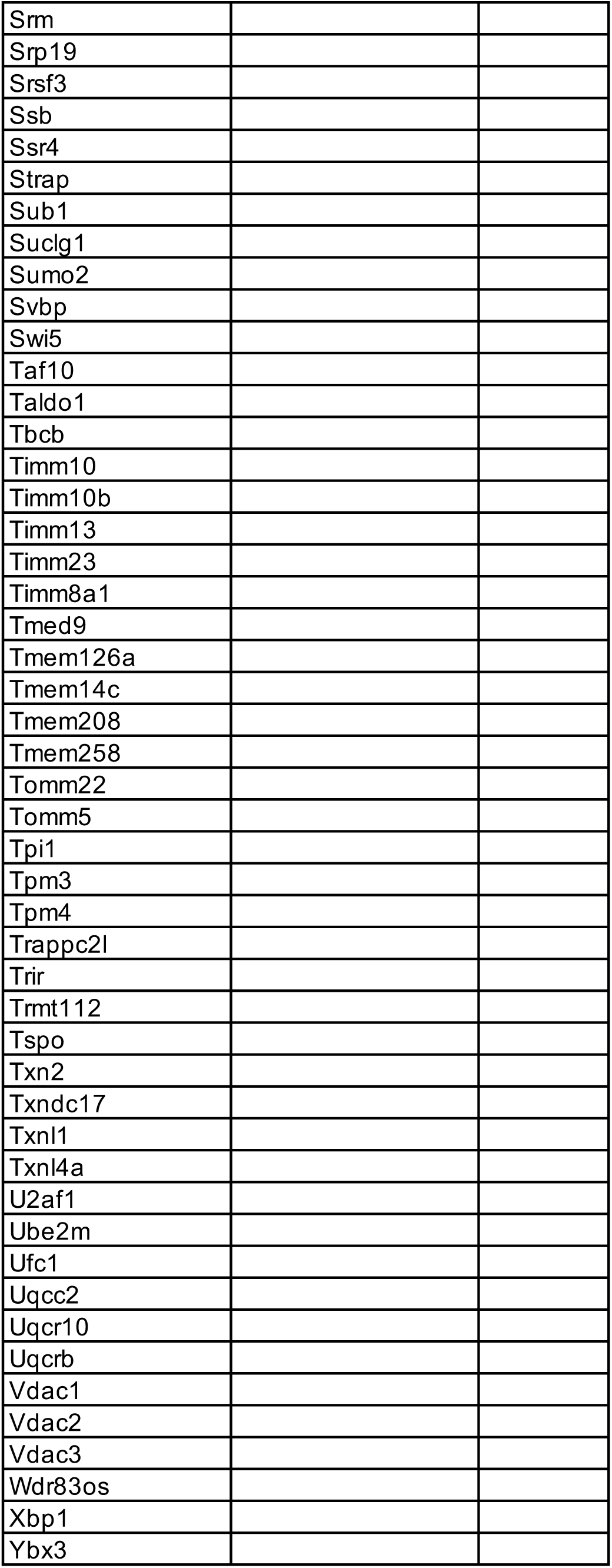
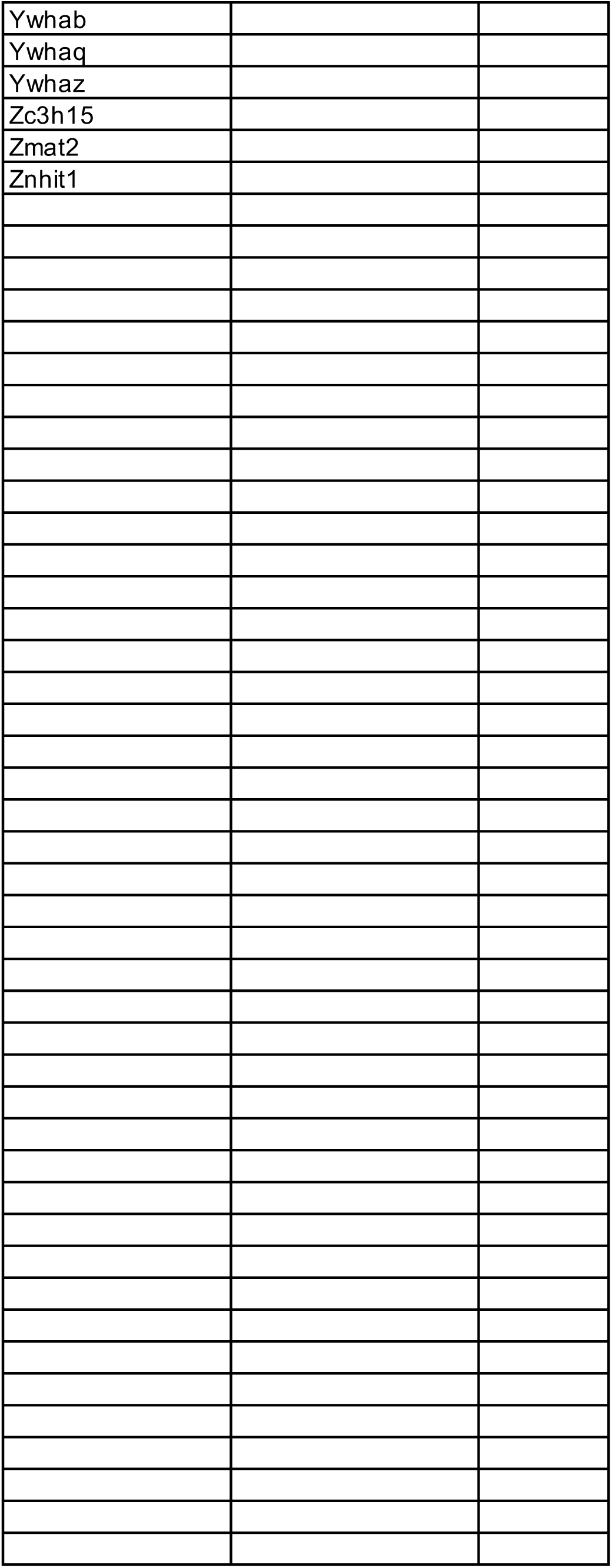
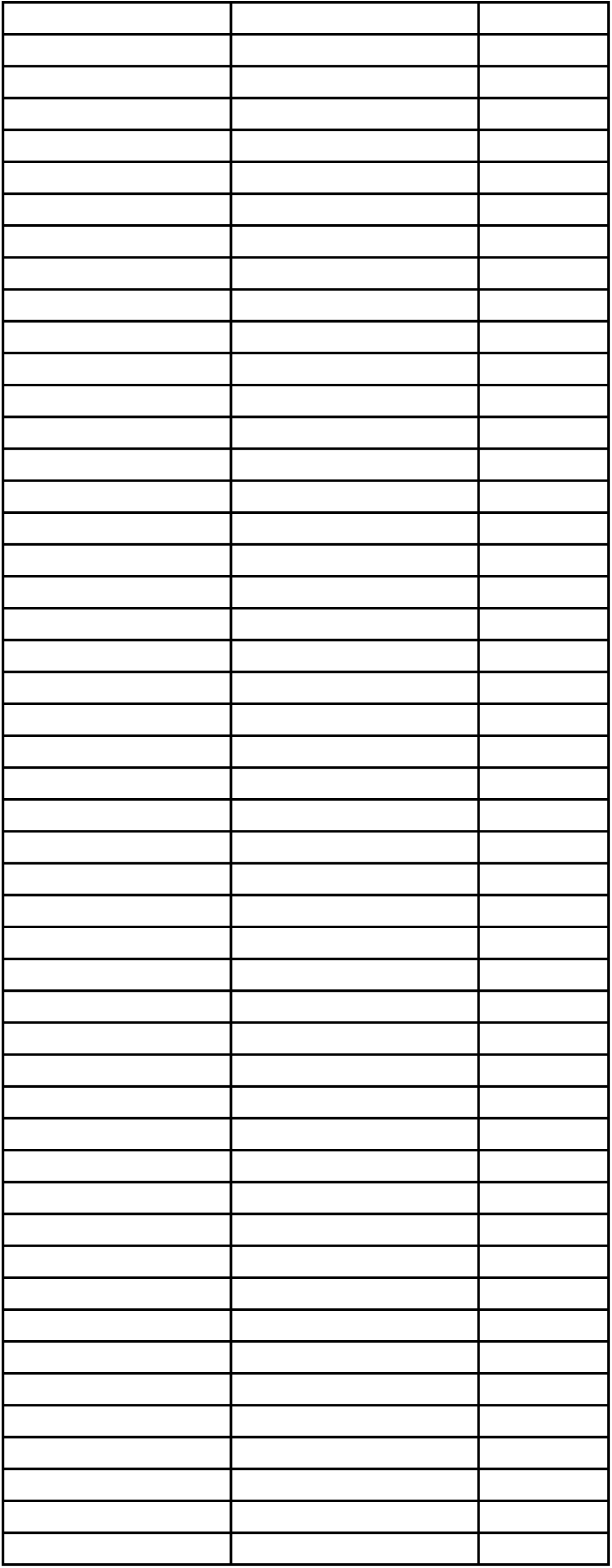
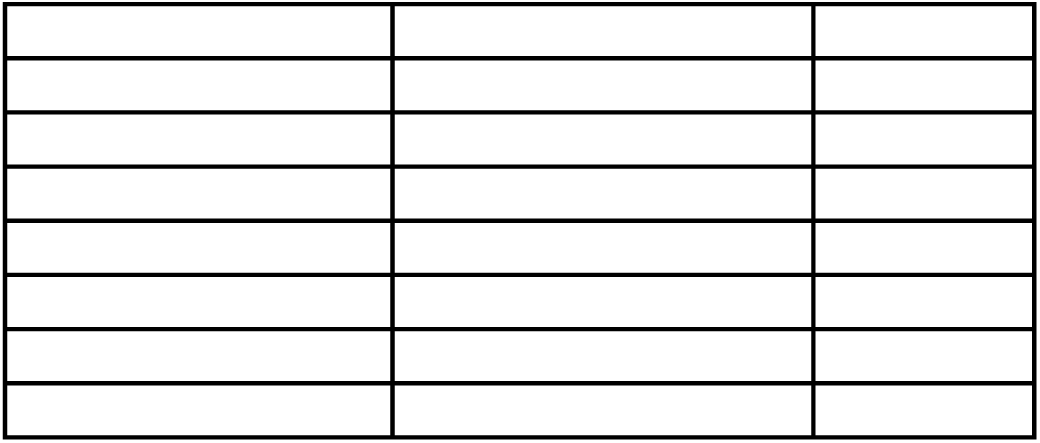
List of genes associated with each metaprogram.

**Table S6:**
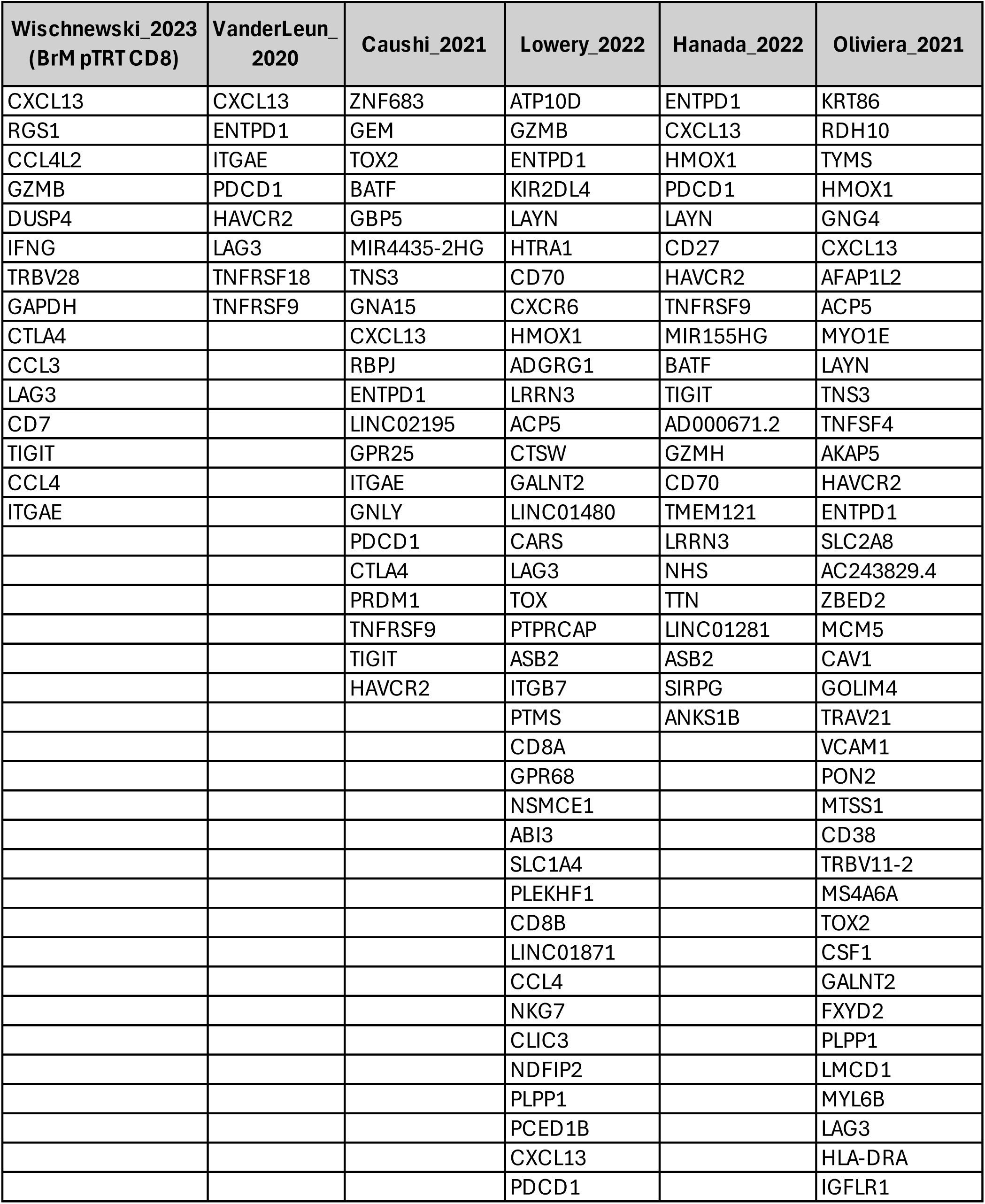
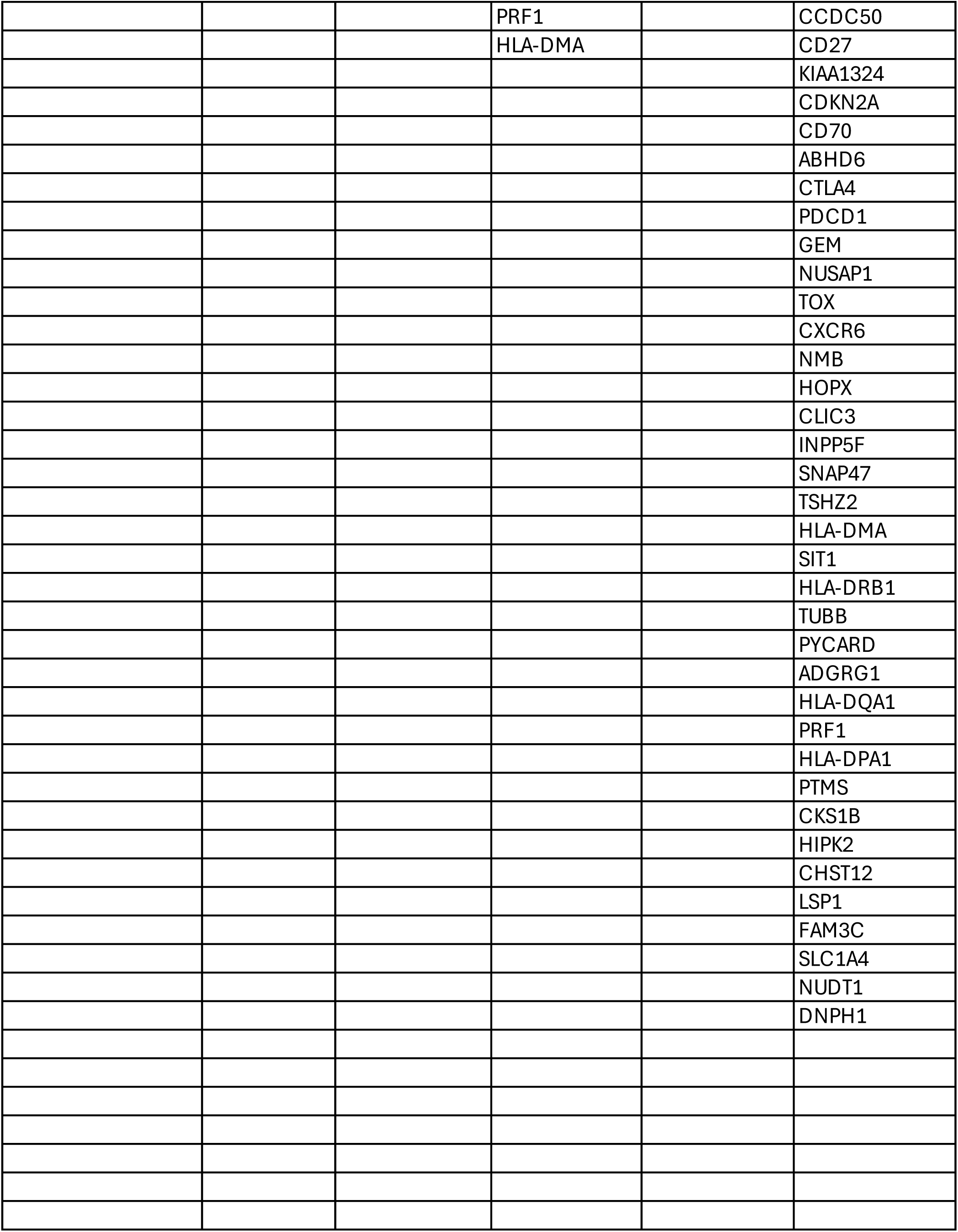
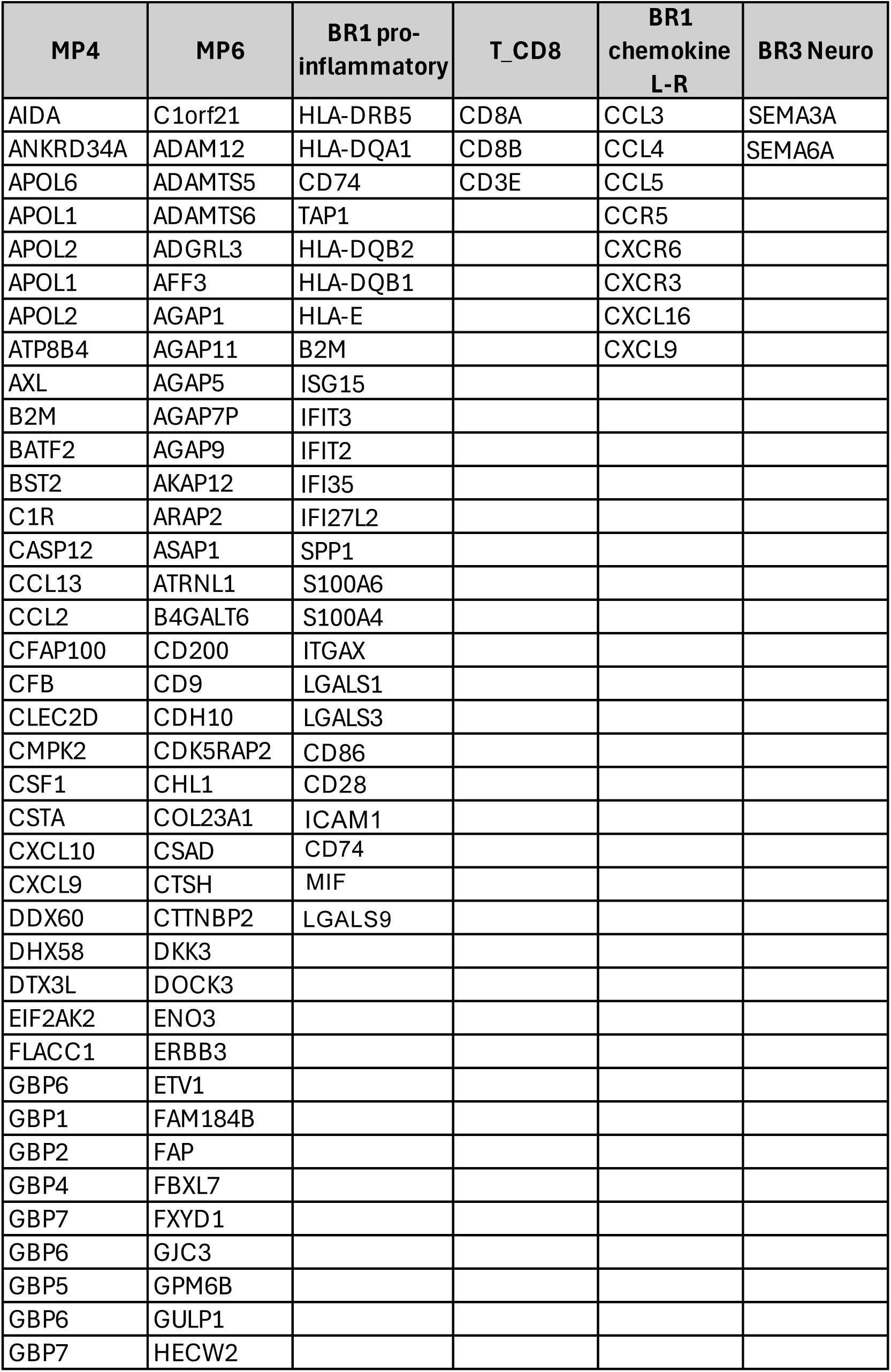
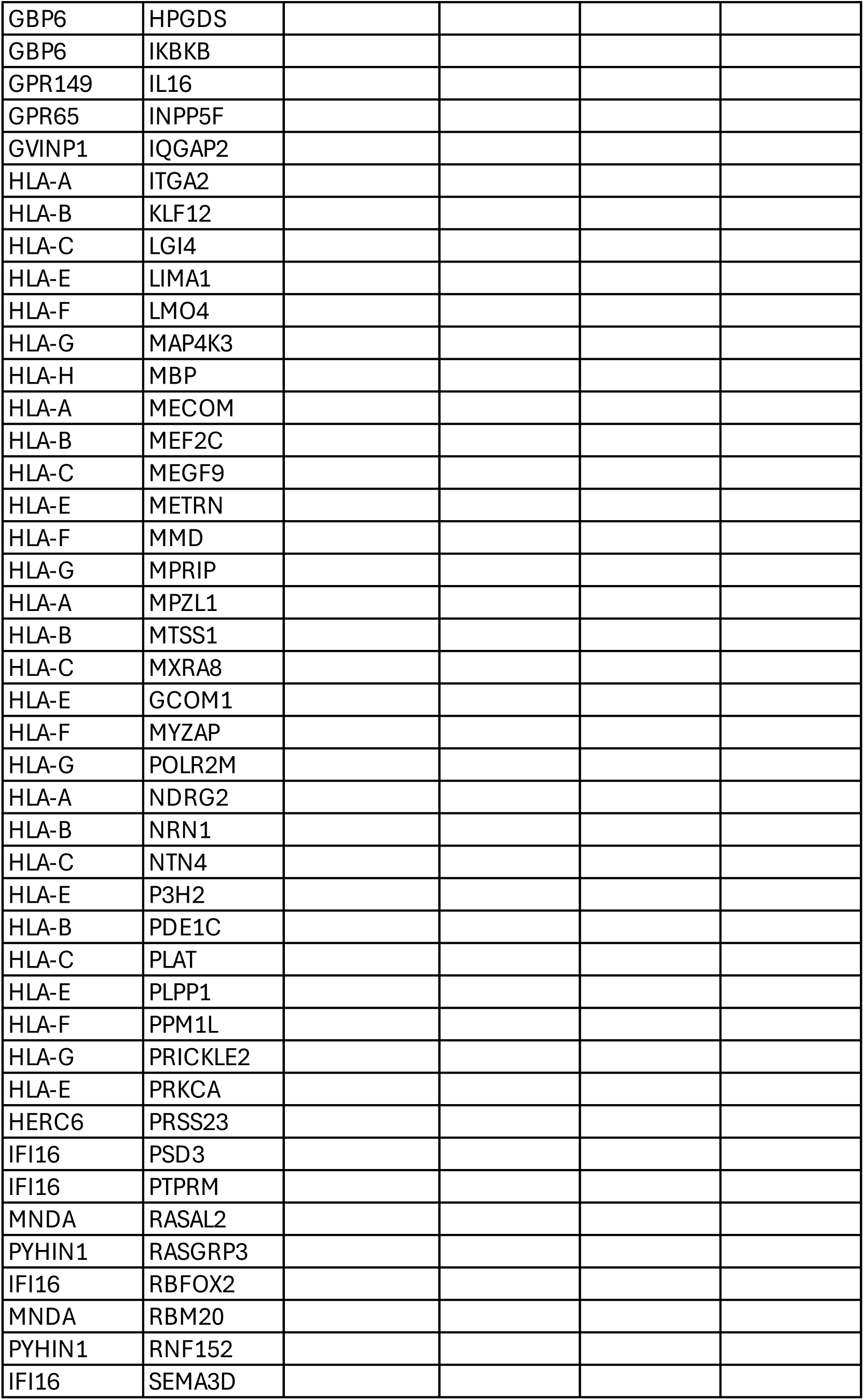
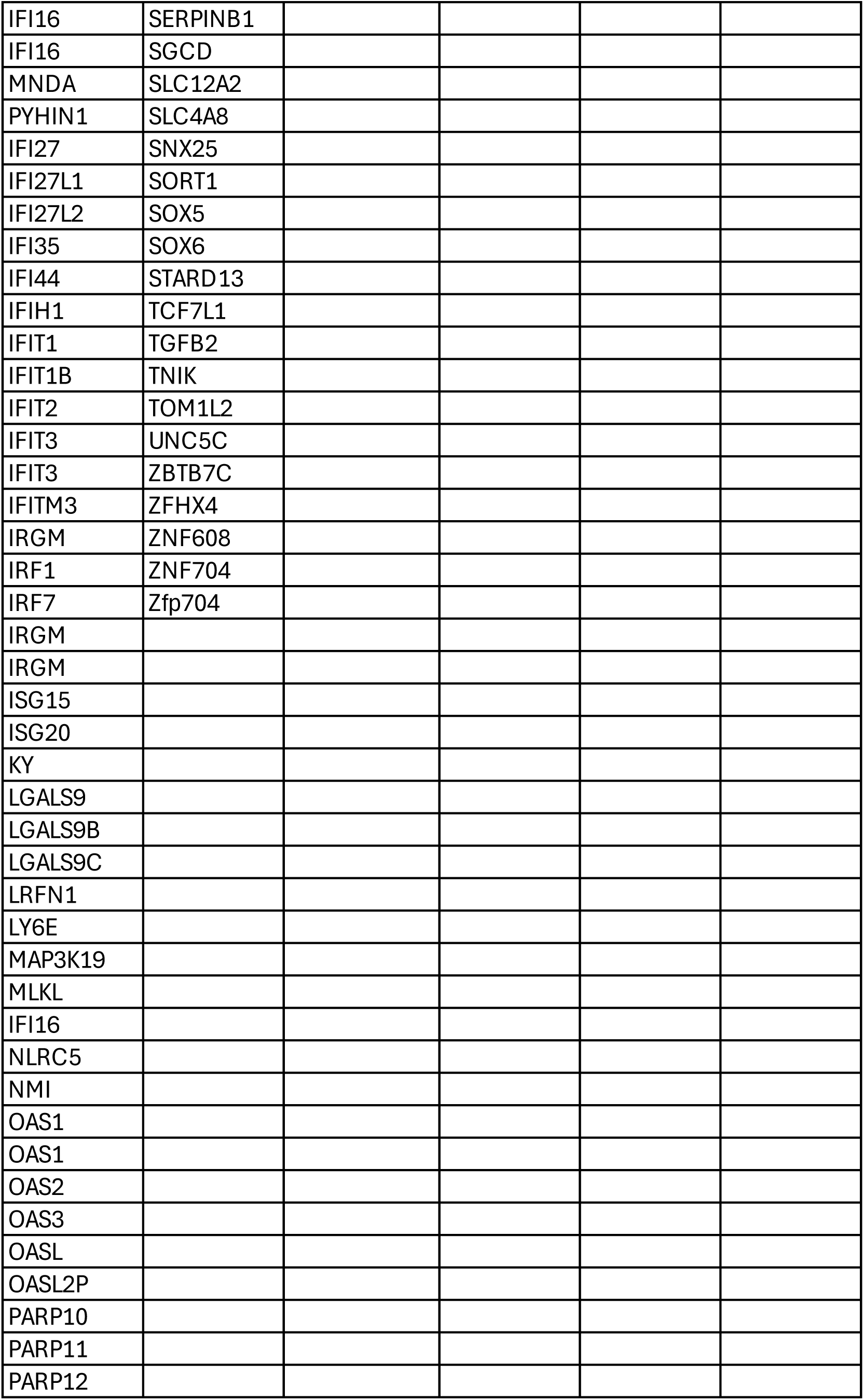
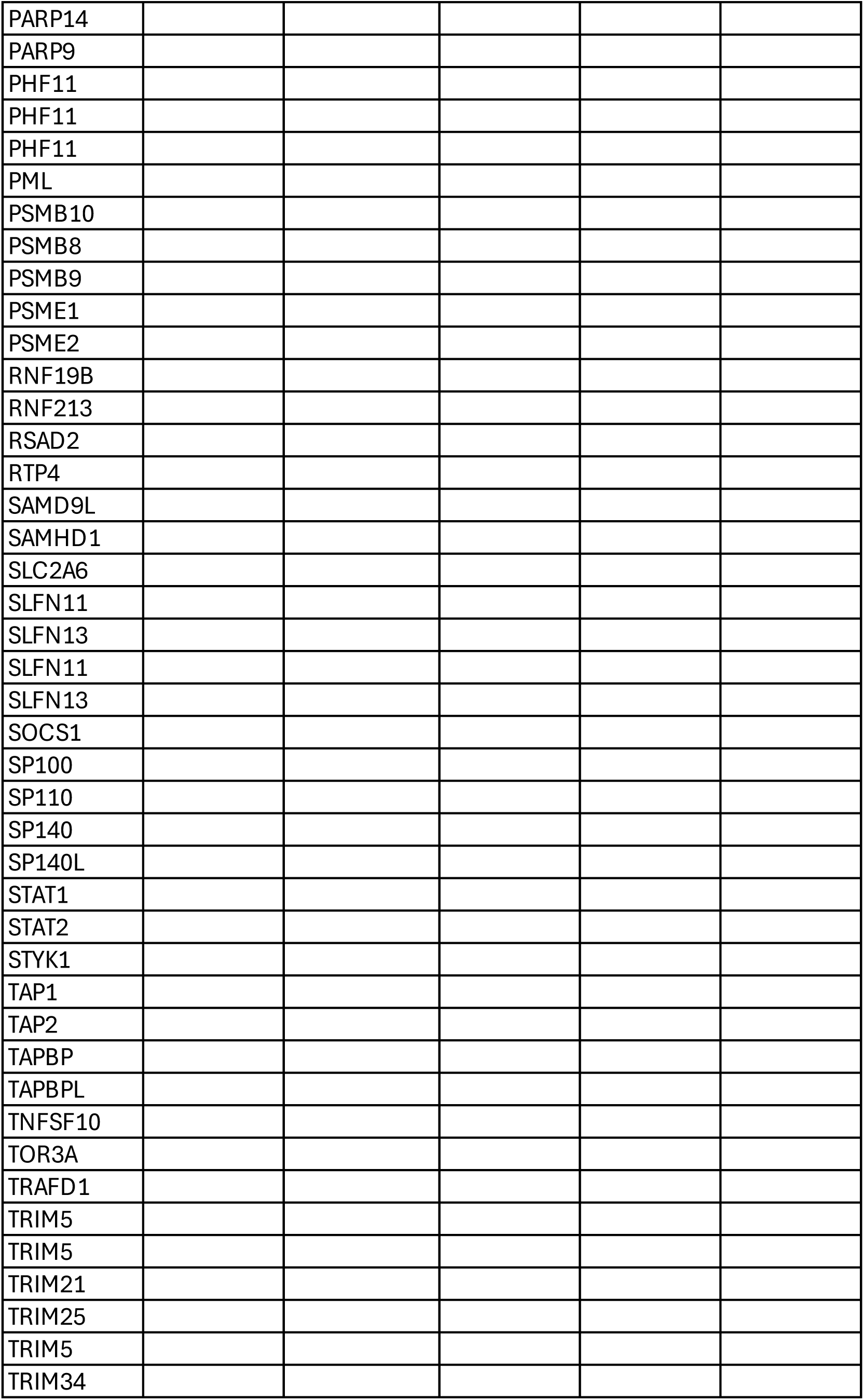
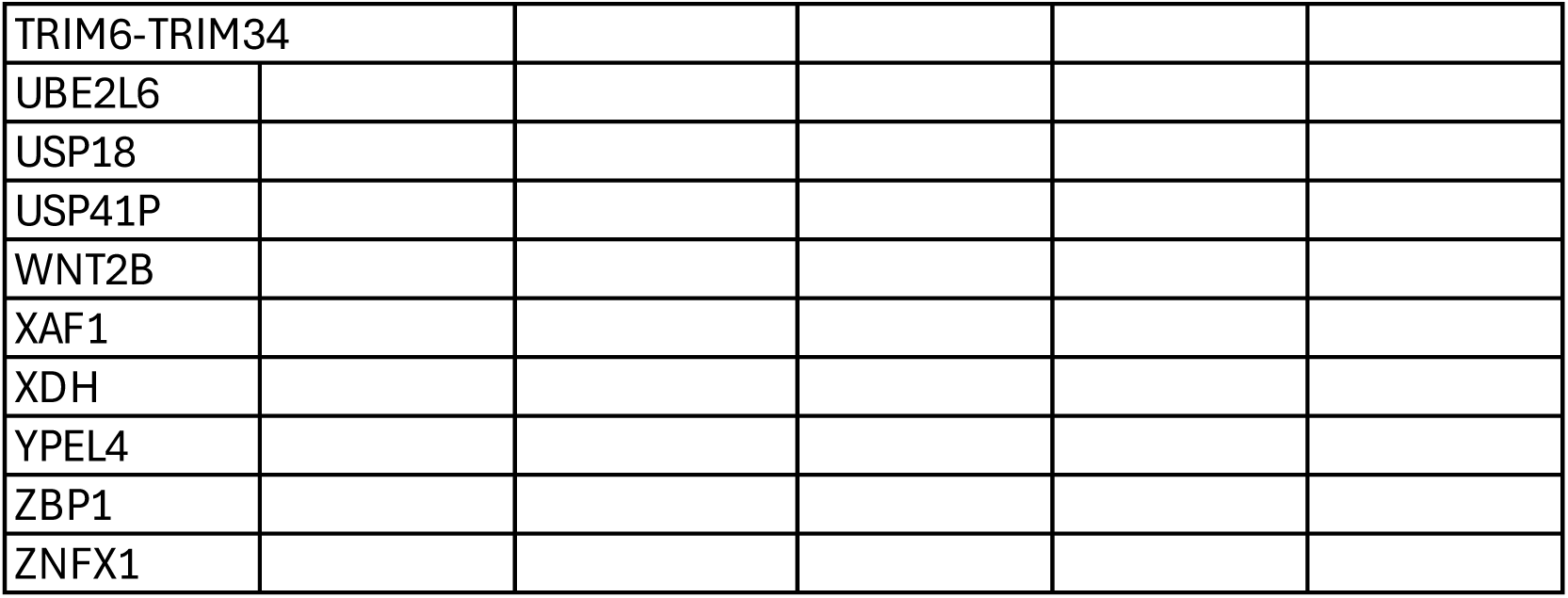
Patient T cell, BR1 and BR3 genes signatures.

## REFERENCES

1. Sperduto, P.W., Mesko, S., Li, J., Cagney, D., Aizer, A., Lin, N.U., Nesbit, E., Kruser, T.J., Chan, J., Braunstein, S., et al. (2020). Survival in Patients With Brain Metastases: Summary Report on the Updated Diagnosis-Specific Graded Prognostic Assessment and Definition of the Eligibility Quotient. J Clin Oncol 38, 3773–3784. 10.1200/JCO.20.01255.

2. Bander, E.D., Yuan, M., Carnevale, J.A., Reiner, A.S., Panageas, K.S., Postow, M.A., Tabar, V., and Moss, N.S. (2021). Melanoma brain metastasis presentation, treatment, and outcomes in the age of targeted and immunotherapies. Cancer 127, 2062–2073. 10.1002/cncr.33459.

3. Long, G.V., Atkinson, V., Lo, S., Sandhu, S., Guminski, A.D., Brown, M.P., Wilmott, J.S., Edwards, J., Gonzalez, M., Scolyer, R.A., et al. (2018). Combination nivolumab and ipilimumab or nivolumab alone in melanoma brain metastases: a multicentre randomised phase 2 study. The Lancet Oncology 19, 672– 681. 10.1016/S1470-2045(18)30139-6.

4. Tawbi, H.A., Forsyth, P.A., Hodi, F.S., Algazi, A.P., Hamid, O., Lao, C.D., Moschos, S.J., Atkins, M.B., Lewis, K., Postow, M.A., et al. (2021). Long-term outcomes of patients with active melanoma brain metastases treated with combination nivolumab plus ipilimumab (CheckMate 204): final results of an open-label, multicentre, phase 2 study. The Lancet Oncology 22, 1692–1704. 10.1016/S1470-2045(21)00545-3.

5. Tawbi, H.A., Forsyth, P.A., Algazi, A., Hamid, O., Hodi, F.S., Moschos, S.J., Khushalani, N.I., Lewis, K., Lao, C.D., Postow, M.A., et al. (2018). Combined Nivolumab and Ipilimumab in Melanoma Metastatic to the Brain. N Engl J Med 379, 722–730. 10.1056/NEJMoa1805453.

6. Tawbi, H.A., Forsyth, P.A., Hodi, F.S., Lao, C.D., Moschos, S.J., Hamid, O., Atkins, M.B., Lewis, K., Thomas, R.P., Glaspy, J.A., et al. (2021). Safety and efficacy of the combination of nivolumab plus ipilimumab in patients with melanoma and asymptomatic or symptomatic brain metastases (CheckMate 204). Neuro Oncol 23, 1961–1973. 10.1093/neuonc/noab094.

7. Brastianos, P.K., Kim, A.E., Giobbie-Hurder, A., Lee, E.Q., Lin, N.U., Overmoyer, B., Wen, P.Y., Nayak, L., Cohen, J.V., Dietrich, J., et al. (2023). Pembrolizumab in brain metastases of diverse histologies: phase 2 trial results. Nat Med 29, 1728–1737. 10.1038/s41591-023-02392-7.

8. Quail, D.F., and Joyce, J.A. (2017). The Microenvironmental Landscape of Brain Tumors. Cancer Cell 31, 326–341. 10.1016/j.ccell.2017.02.009.

9. Aili, Y., Maimaitiming, N., Qin, H., Ji, W., Fan, G., Wang, Z., and Wang, Y. (2022). Tumor microenvironment and exosomes in brain metastasis: Molecular mechanisms and clinical application. Front. Oncol. 12. 10.3389/fonc.2022.983878.

10. Caffarel, M.M., and Braza, M.S. (2022). Microglia and metastases to the central nervous system: victim, ravager, or something else? J Exp Clin Cancer Res 41, 1–13. 10.1186/s13046-022-02535-7.

11. Arvanitis, C.D., Ferraro, G.B., and Jain, R.K. (2020). The blood–brain barrier and blood–tumour barrier in brain tumours and metastases. Nat Rev Cancer 20, 26–41. 10.1038/s41568-019-0205-x.

12. Wu, D., Chen, Q., Chen, X., Han, F., Chen, Z., and Wang, Y. (2023). The blood-brain barrier: structure, regulation, and drug delivery. Signal Transduct Target Ther 8, 217. 10.1038/s41392-023-01481-w.

13. Biermann, J., Melms, J.C., Amin, A.D., Wang, Y., Caprio, L.A., Karz, A., Tagore, S., Barrera, I., Ibarra-Arellano, M.A., Andreatta, M., et al. (2022). Dissecting the treatment-naive ecosystem of human melanoma brain metastasis. Cell 185, 2591–2608.e30. 10.1016/j.cell.2022.06.007.

14. Fischer, G.M., Jalali, A., Kircher, D.A., Lee, W.-C., McQuade, J.L., Haydu, L.E., Joon, A.Y., Reuben, A., de Macedo, M.P., Carapeto, F.C.L., et al. (2019). Molecular Profiling Reveals Unique Immune and Metabolic Features of Melanoma Brain Metastases. Cancer Discov 9, 628–645. 10.1158/2159-8290.CD-18-1489.

15. Lusby, R., Carl, S., and Tiwari, V.K. (2024). Integrating single-cell transcriptomics with Artificial Intelligence reveals pan-cancer biomarkers of brain metastasis. bioRxiv, 2024.03.08.584083. 10.1101/2024.03.08.584083.

16. Ikarashi, D., Okimoto, T., Shukuya, T., Onagi, H., Hayashi, T., Sinicropi-Yao, S.L., Amann, J.M., Nakatsura, T., Kitano, S., and Carbone, D.P. (2021). Comparison of Tumor Microenvironments Between Primary Tumors and Brain Metastases in Patients With NSCLC. JTO Clin Res Rep 2, 100230. 10.1016/j.jtocrr.2021.100230.

17. Friebel, E., Kapolou, K., Unger, S., Núñez, N.G., Utz, S., Rushing, E.J., Regli, L., Weller, M., Greter, M., Tugues, S., et al. (2020). Single-Cell Mapping of Human Brain Cancer Reveals Tumor-Specific Instruction of Tissue-Invading Leukocytes. Cell 181, 1626–1642.e20. 10.1016/j.cell.2020.04.055.

18. Klemm, F., Maas, R.R., Bowman, R.L., Kornete, M., Soukup, K., Nassiri, S., Brouland, J.-P., Iacobuzio-Donahue, C.A., Brennan, C., Tabar, V., et al. (2020). Interrogation of the Microenvironmental Landscape in Brain Tumors Reveals Disease-Specific Alterations of Immune Cells. Cell 181, 1643–1660.e17. 10.1016/j.cell.2020.05.007.

19. Wischnewski, V., Maas, R.R., Aruffo, P.G., Soukup, K., Galletti, G., Kornete, M., Galland, S., Fournier, N., Lilja, J., Wirapati, P., et al. (2023). Phenotypic diversity of T cells in human primary and metastatic brain tumors revealed by multiomic interrogation. Nat Cancer 4, 908–924. 10.1038/s43018-023-00566-3.

20. Gonzalez, H., Mei, W., Robles, I., Hagerling, C., Allen, B.M., Hauge Okholm, T.L., Nanjaraj, A., Verbeek, T., Kalavacherla, S., van Gogh, M., et al. (2022). Cellular architecture of human brain metastases. Cell 185, 729–745.e20. 10.1016/j.cell.2021.12.043.

21. Sun, L., Kienzler, J.C., Reynoso, J.G., Lee, A., Shiuan, E., Li, S., Kim, J., Ding, L., Monteleone, A.J., Owens, G.C., et al. (2023). Immune checkpoint blockade induces distinct alterations in the microenvironments of primary and metastatic brain tumors. J Clin Invest 133, e169314. 10.1172/JCI169314.

22. Álvarez-Prado, Á.F., Maas, R.R., Soukup, K., Klemm, F., Kornete, M., Krebs, F.S., Zoete, V., Berezowska, S., Brouland, J.-P., Hottinger, A.F., et al. (2023). Immunogenomic analysis of human brain metastases reveals diverse immune landscapes across genetically distinct tumors. Cell Reports Medicine 4, 100900. 10.1016/j.xcrm.2022.100900.

23. Miarka, L., and Valiente, M. (2021). Animal models of brain metastasis. Neurooncol Adv 3, v144–v156. 10.1093/noajnl/vdab115.

24. Valiente, M., Van Swearingen, A.E.D., Anders, C.K., Bairoch, A., Boire, A., Bos, P.D., Cittelly, D.M., Erez, N., Ferraro, G.B., Fukumura, D., et al. (2020). Brain Metastasis Cell Lines Panel: A Public Resource of Organotropic Cell Lines. Cancer Research 80, 4314–4323. 10.1158/0008-5472.CAN-20-0291.

25. Cho, J.H., Robinson, J.P., Arave, R.A., Burnett, W.J., Kircher, D.A., Chen, G., Davies, M.A., Grossmann, A.H., VanBrocklin, M.W., McMahon, M., et al. (2015). AKT1 Activation Promotes Development of Melanoma Metastases. Cell Rep 13, 898–905. 10.1016/j.celrep.2015.09.057.

26. Kircher, D.A., Trombetti, K.A., Silvis, M.R., Parkman, G.L., Fischer, G.M., Angel, S.N., Stehn, C.M., Strain, S.C., Grossmann, A.H., Duffy, K.L., et al. (2019). AKT1E17K Activates Focal Adhesion Kinase and Promotes Melanoma Brain Metastasis. Mol Cancer Res 17, 1787–1800. 10.1158/1541-7786.MCR-18-1372.

27. Pérez-Guijarro, E., Yang, H.H., Araya, R.E., El Meskini, R., Michael, H.T., Vodnala, S.K., Marie, K.L., Smith, C., Chin, S., Lam, K.C., et al. (2020). Multimodel preclinical platform predicts clinical response of melanoma to immunotherapy. Nat Med 26, 781–791. 10.1038/s41591-020-0818-3.

28. Bloom, M.B., Perry-Lalley, D., Robbins, P.F., Li, Y., El-Gamil, M., Rosenberg, S.A., and Yang, J.C. (1997). Identification of Tyrosinase-related Protein 2 as a Tumor Rejection Antigen for the B16 Melanoma. J Exp Med 185, 453–460.

29. Tímár, J., Mészáros, L., Ladányi, A., Puskás, L.G., and Rásó, E. (2006). Melanoma genomics reveals signatures of sensitivity to bio- and targeted therapies. Cell Immunol 244, 154–157. 10.1016/j.cellimm.2006.12.009.

30. Rappaport, J., Chen, Q., McGuire, T., Daugherty-Lopes, A., and Goldszmid, R.S. (2024). Development of an optimized machine learning approach to enhance brain metastatic burden assessment in preclinical models. bioRxiv, 2024.08.21.608131. 10.1101/2024.08.21.608131.

31. Nguyen, B., Fong, C., Luthra, A., Smith, S.A., DiNatale, R.G., Nandakumar, S., Walch, H., Chatila, W.K., Madupuri, R., Kundra, R., et al. (2022). Genomic characterization of metastatic patterns from prospective clinical sequencing of 25,000 patients. Cell 185, 563–575.e11. 10.1016/j.cell.2022.01.003.

32. Cerami1, E., Gao, J., Dogrusoz, U., Gross, B.E., Sumer, S.O., Aksoy, B.A., Jacobsen, A., Byrne, C.J., Heuer, M.L., Larsson, E., et al. (2012). The cBio Cancer Genomics Portal: An Open Platform for Exploring Multidimensional Cancer Genomics Data. Cancer Discov 2, 401–404. 10.1158/2159-8290.CD-12-0095.

33. Gao, J., Aksoy, B.A., Dogrusoz, U., Dresdner, G., Gross, B., Sumer, S.O., Sun, Y., Jacobsen, A., Sinha, R., Larsson, E., et al. (2013). Integrative Analysis of Complex Cancer Genomics and Clinical Profiles Using the cBioPortal. Science Signaling. 10.1126/scisignal.2004088.

34. de Bruijn, I., Kundra, R., Mastrogiacomo, B., Tran, T.N., Sikina, L., Mazor, T., Li, X., Ochoa, A., Zhao, G., Lai, B., et al. (2023). Analysis and Visualization of Longitudinal Genomic and Clinical Data from the AACR Project GENIE Biopharma Collaborative in cBioPortal. Cancer Res 83, 3861–3867. 10.1158/0008-5472.CAN-23-0816.

35. Chen, G., Chakravarti, N., Aardalen, K., Lazar, A.J., Tetzlaff, M.T., Wubbenhorst, B., Kim, S.-B., Kopetz, S., Ledoux, A.A., Gopal, Y.N.V., et al. (2014). Molecular Profiling of Patient-Matched Brain and Extracranial Melanoma Metastases Implicates the PI3K Pathway as a Therapeutic Target. Clin Cancer Res 20, 5537– 5546. 10.1158/1078-0432.CCR-13-3003.

36. Brastianos, P.K., Carter, S.L., Santagata, S., Cahill, D.P., Taylor-Weiner, A., Jones, R.T., Van Allen, E.M., Lawrence, M.S., Horowitz, P.M., Cibulskis, K., et al. (2015). Genomic Characterization of Brain Metastases Reveals Branched Evolution and Potential Therapeutic Targets. Cancer Discov 5, 1164–1177. 10.1158/2159-8290.CD-15-0369.

37. Morad, G., Helmink, B.A., Sharma, P., and Wargo, J.A. (2021). Hallmarks of response, resistance, and toxicity to immune checkpoint blockade. Cell 184, 5309–5337. 10.1016/j.cell.2021.09.020.

38. Petitprez, F., Meylan, M., de Reyniès, A., Sautès-Fridman, C., and Fridman, W.H. (2020). The Tumor Microenvironment in the Response to Immune Checkpoint Blockade Therapies. Front Immunol 11, 784. 10.3389/fimmu.2020.00784.

39. Lam, K.C., Araya, R.E., Huang, A., Chen, Q., Di Modica, M., Rodrigues, R.R., Lopès, A., Johnson, S.B., Schwarz, B., Bohrnsen, E., et al. (2021). Microbiota triggers STING-type I IFN-dependent monocyte reprogramming of the tumor microenvironment. Cell 184, 5338–5356.e21. 10.1016/j.cell.2021.09.019.

40. Smalley, I., Chen, Z., Phadke, M., Li, J., Yu, X., Wyatt, C., Evernden, B., Messina, J.L., Sarnaik, A., Sondak, V.K., et al. (2021). Single-Cell Characterization of the Immune Microenvironment of Melanoma Brain and Leptomeningeal Metastases. Clin Cancer Res 27, 4109–4125. 10.1158/1078-0432.CCR-21-1694.

41. Zakaria, R., Jenkinson, M.D., Radon, M., Das, K., Poptani, H., Rathi, N., and Rudland, P.S. (2023). Immune checkpoint inhibitor treatment of brain metastasis associated with a less invasive growth pattern, higher T-cell infiltration and raised tumor ADC on diffusion weighted MRI. Cancer Immunol Immunother 72, 3387–3393. 10.1007/s00262-023-03499-z.

42. Andreatta, M., Corria-Osorio, J., Müller, S., Cubas, R., Coukos, G., and Carmona, S.J. (2021). Interpretation of T cell states from single-cell transcriptomics data using reference atlases. Nat Commun 12, 2965. 10.1038/s41467-021-23324-4.

43. Sade-Feldman, M., Yizhak, K., Bjorgaard, S.L., Ray, J.P., de Boer, C.G., Jenkins, R.W., Lieb, D.J., Chen, J.H., Frederick, D.T., Barzily-Rokni, M., et al. (2018). Defining T cell states associated with response to checkpoint immunotherapy in melanoma. Cell 175, 998–1013.e20. 10.1016/j.cell.2018.10.038.

44. Giles, J.R., Globig, A.-M., Kaech, S.M., and Wherry, E.J. (2023). CD8+ T cells in the cancer-immunity cycle. Immunity 56, 2231–2253. 10.1016/j.immuni.2023.09.005.

45. Miller, B.C., Sen, D.R., Al Abosy, R., Bi, K., Virkud, Y.V., LaFleur, M.W., Yates, K.B., Lako, A., Felt, K., Naik, G.S., et al. (2019). Subsets of exhausted CD8+ T cells differentially mediate tumor control and respond to checkpoint blockade. Nat Immunol 20, 326–336. 10.1038/s41590-019-0312-6.

46. Jiang, W., He, Y., He, W., Wu, G., Zhou, X., Sheng, Q., Zhong, W., Lu, Y., Ding, Y., Lu, Q., et al. (2021). Exhausted CD8+T Cells in the Tumor Immune Microenvironment: New Pathways to Therapy. Front. Immunol. 11, 622509. 10.3389/fimmu.2020.622509.

47. Acharya, N., Madi, A., Zhang, H., Klapholz, M., Escobar, G., Dulberg, S., Christian, E., Ferreira, M., Dixon, K.O., Fell, G., et al. (2020). Endogenous Glucocorticoid Signaling Regulates CD8+ T Cell Differentiation and Development of Dysfunction in the Tumor Microenvironment. Immunity 53, 658–671.e6. 10.1016/j.immuni.2020.08.005.

48. Damei, I., Trickovic, T., Mami-Chouaib, F., and Corgnac, S. (2023). Tumor-resident memory T cells as a biomarker of the response to cancer immunotherapy. Front. Immunol. 14, 1205984. 10.3389/fimmu.2023.1205984.

49. Corgnac, S., Malenica, I., Mezquita, L., Auclin, E., Voilin, E., Kacher, J., Halse, H., Grynszpan, L., Signolle, N., Dayris, T., et al. (2020). CD103+CD8+ TRM Cells Accumulate in Tumors of Anti-PD-1-Responder Lung Cancer Patients and Are Tumor-Reactive Lymphocytes Enriched with Tc17. Cell Reports Medicine 1. 10.1016/j.xcrm.2020.100127.

50. Edwards, J., Wilmott, J.S., Madore, J., Gide, T.N., Quek, C., Tasker, A., Ferguson, A., Chen, J., Hewavisenti, R., Hersey, P., et al. (2018). CD103+ Tumor-Resident CD8+ T Cells Are Associated with Improved Survival in Immunotherapy-Naïve Melanoma Patients and Expand Significantly During Anti–PD-1 Treatment. Clinical Cancer Research 24, 3036–3045. 10.1158/1078-0432.CCR-17-2257.

51. Colonna, M., and Butovsky, O. (2017). Microglia Function in the Central Nervous System During Health and Neurodegeneration. Annu Rev Immunol 35, 441–468. 10.1146/annurev-immunol-051116-052358.

52. Borst, K., Dumas, A.A., and Prinz, M. (2021). Microglia: Immune and non-immune functions. Immunity 54, 2194–2208. 10.1016/j.immuni.2021.09.014.

53. Paolicelli, R.C., Sierra, A., Stevens, B., Tremblay, M.-E., Aguzzi, A., Ajami, B., Amit, I., Audinat, E., Bechmann, I., Bennett, M., et al. (2022). Microglia states and nomenclature: A field at its crossroads. Neuron 110, 3458. 10.1016/j.neuron.2022.10.020.

54. Guldner, I.H., Wang, Q., Yang, L., Golomb, S.M., Zhao, Z., Lopez, J.A., Brunory, A., Howe, E.N., Zhang, Y., Palakurthi, B., et al. (2020). CNS-Native Myeloid Cells Drive Immune Suppression in the Brain Metastatic Niche through Cxcl10. Cell 183, 1234–1248.e25. 10.1016/j.cell.2020.09.064.

55. You, H., Baluszek, S., and Kaminska, B. (2019). Immune Microenvironment of Brain Metastases—Are Microglia and Other Brain Macrophages Little Helpers? Front. Immunol. 10, 1941. 10.3389/fimmu.2019.01941.

56. Evans, K.T., Blake, K., Longworth, A., Coburn, M.A., Insua-Rodríguez, J., McMullen, T.P., Nguyen, Q.H., Ma, D., Lev, T., Hernandez, G.A., et al. (2023). Microglia promote anti-tumour immunity and suppress breast cancer brain metastasis. Nat Cell Biol 25, 1848–1859. 10.1038/s41556-023-01273-y.

57. Lin, S.-S., Tang, Y., Illes, P., and Verkhratsky, A. (2021). The Safeguarding Microglia: Central Role for P2Y12 Receptors. Front Pharmacol 11, 627760. 10.3389/fphar.2020.627760.

58. Huang, C., Wang, X., Wang, Y., Feng, Y., Wang, X., Chen, S., Yan, P., Liao, J., Zhang, Q., Mao, C., et al. (2024). Sirpα on tumor-associated myeloid cells restrains antitumor immunity in colorectal cancer independent of its interaction with CD47. Nat Cancer 5, 500–516. 10.1038/s43018-023-00691-z.

59. Hutter, G., Theruvath, J., Graef, C.M., Zhang, M., Schoen, M.K., Manz, E.M., Bennett, M.L., Olson, A., Azad, T.D., Sinha, R., et al. (2019). Microglia are effector cells of CD47-SIRPα antiphagocytic axis disruption against glioblastoma. Proceedings of the National Academy of Sciences 116, 997–1006. 10.1073/pnas.1721434116.

60. Bian, Z., Shi, L., Kidder, K., Zen, K., Garnett-Benson, C., and Liu, Y. (2021). Intratumoral SIRPα-deficient macrophages activate tumor antigen-specific cytotoxic T cells under radiotherapy. Nat Commun 12, 1–16. 10.1038/s41467-021-23442-z.

61. Niu, F., Yu, Y., Li, Z., Ren, Y., Li, Z., Ye, Q., Liu, P., Ji, C., Qian, L., and Xiong, Y. (2022). Arginase: An emerging and promising therapeutic target for cancer treatment. Biomed Pharmacother 149, 112840. 10.1016/j.biopha.2022.112840.

62. Dixon, K.O., Lahore, G.F., and Kuchroo, V.K. (2024). Beyond T cell exhaustion: TIM-3 regulation of myeloid cells. Science Immunology. 10.1126/sciimmunol.adf2223.

63. Ausejo-Mauleon, I., Labiano, S., de la Nava, D., Laspidea, V., Zalacain, M., Marrodán, L., García-Moure, M., González-Huarriz, M., Hervás-Corpión, I., Dhandapani, L., et al. (2023). TIM-3 blockade in diffuse intrinsic pontine glioma models promotes tumor regression and antitumor immune memory. Cancer Cell 41, 1911–1926.e8. 10.1016/j.ccell.2023.09.001.

64. Ocaña-Guzman, R., Torre-Bouscoulet, L., and Sada-Ovalle, I. (2016). TIM-3 Regulates Distinct Functions in Macrophages. Front Immunol 7, 229. 10.3389/fimmu.2016.00229.

65. Finak, G., McDavid, A., Yajima, M., Deng, J., Gersuk, V., Shalek, A.K., Slichter, C.K., Miller, H.W., McElrath, M.J., Prlic, M., et al. (2015). MAST: a flexible statistical framework for assessing transcriptional changes and characterizing heterogeneity in single-cell RNA sequencing data. Genome Biol 16, 278. 10.1186/s13059-015-0844-5.

66. Wang, T., Li, B., Nelson, C.E., and Nabavi, S. (2019). Comparative analysis of differential gene expression analysis tools for single-cell RNA sequencing data. BMC Bioinformatics 20, 40. 10.1186/s12859-019-2599-6.

67. Lin, X., and Boutros, P.C. (2020). Optimization and expansion of non-negative matrix factorization. BMC Bioinformatics 21, 7. 10.1186/s12859-019-3312-5.

68. Tirosh, I., Venteicher, A.S., Hebert, C., Escalante, L.E., Patel, A.P., Yizhak, K., Fisher, J.M., Rodman, C., Mount, C., Filbin, M.G., et al. (2016). Single-cell RNA-seq supports a developmental hierarchy in human oligodendroglioma. Nature 539, 309–313. 10.1038/nature20123.

69. Kinker, G.S., Greenwald, A.C., Tal, R., Orlova, Z., Cuoco, M.S., McFarland, J.M., Warren, A., Rodman, C., Roth, J.A., Bender, S.A., et al. (2020). Pan-cancer single-cell RNA-seq identifies recurring programs of cellular heterogeneity. Nat Genet 52, 1208–1218. 10.1038/s41588-020-00726-6.

70. Gavish, A., Tyler, M., Greenwald, A.C., Hoefflin, R., Simkin, D., Tschernichovsky, R., Galili Darnell, N., Somech, E., Barbolin, C., Antman, T., et al. (2023). Hallmarks of transcriptional intratumour heterogeneity across a thousand tumours. Nature 618, 598–606. 10.1038/s41586-023-06130-4.

71. Zhou, Y., Zhou, B., Pache, L., Chang, M., Khodabakhshi, A.H., Tanaseichuk, O., Benner, C., and Chanda, S.K. (2019). Metascape provides a biologist-oriented resource for the analysis of systems-level datasets. Nat Commun 10, 1523. 10.1038/s41467-019-09234-6.

72. Alvarez-Breckenridge, C., Markson, S.C., Stocking, J.H., Nayyar, N., Lastrapes, M., Strickland, M.R., Kim, A.E., de Sauvage, M., Dahal, A., Larson, J.M., et al. (2022). Microenvironmental Landscape of Human Melanoma Brain Metastases in Response to Immune Checkpoint Inhibition. Cancer Immunol Res 10, 996–1012. 10.1158/2326-6066.CIR-21-0870.

73. Jin, C., Shao, Y., Zhang, X., Xiang, J., Zhang, R., Sun, Z., Mei, S., Zhou, J., Zhang, J., and Shi, L. (2021). A Unique Type of Highly-Activated Microglia Evoking Brain Inflammation via Mif/Cd74 Signaling Axis in Aged Mice. Aging Dis 12, 2125–2139. 10.14336/AD.2021.0520.

74. Potru, P.S., and Spittau, B. (2023). CD74: a prospective marker for reactive microglia? Neural Regen Res 18, 2673–2674. 10.4103/1673-5374.371350.

75. Jahn, J., Bollensdorf, A., Kalischer, C., Piecha, R., Weiß-Müller, J., Potru, P.S., Ruß, T., and Spittau, B. (2022). Microglial CD74 Expression Is Regulated by TGFβ Signaling. Int J Mol Sci 23, 10247. 10.3390/ijms231810247.

76. Wiersma, V.R., de Bruyn, M., Helfrich, W., and Bremer, E. (2013). Therapeutic potential of Galectin-9 in human disease. Medicinal Research Reviews 33, E102–E126. 10.1002/med.20249.

77. Nobumoto, A., Oomizu, S., Arikawa, T., Katoh, S., Nagahara, K., Miyake, M., Nishi, N., Takeshita, K., Niki, T., Yamauchi, A., et al. (2009). Galectin-9 expands unique macrophages exhibiting plasmacytoid dendritic cell-like phenotypes that activate NK cells in tumor-bearing mice. Clinical Immunology 130, 322–330. 10.1016/j.clim.2008.09.014.

78. Ozga, A.J., Chow, M.T., and Luster, A.D. (2021). Chemokines and the immune response to cancer. Immunity 54, 859. 10.1016/j.immuni.2021.01.012.

79. Harlin, H., Meng, Y., Peterson, A.C., Zha, Y., Tretiakova, M., Slingluff, C., McKee, M., and Gajewski, T.F. (2009). Chemokine Expression in Melanoma Metastases Associated with CD8+ T-Cell Recruitment. Cancer Res 69, 3077–3085. 10.1158/0008-5472.CAN-08-2281.

80. Allen, F., Bobanga, I.D., Rauhe, P., Barkauskas, D., Teich, N., Tong, C., Myers, J., and Huang, A.Y. (2017). CCL3 augments tumor rejection and enhances CD8+ T cell infiltration through NK and CD103+ dendritic cell recruitment via IFNγ. Oncoimmunology 7, e1393598. 10.1080/2162402X.2017.1393598.

81. Caushi, J.X., Zhang, J., Ji, Z., Vaghasia, A., Zhang, B., Hsiue, E.H.-C., Mog, B.J., Hou, W., Justesen, S., Blosser, R., et al. (2021). Transcriptional programs of neoantigen-specific TIL in anti-PD-1-treated lung cancers. Nature 596, 126–132. 10.1038/s41586-021-03752-4.

82. Lowery, F.J., Krishna, S., Yossef, R., Parikh, N.B., Chatani, P.D., Zacharakis, N., Parkhurst, M.R., Levin, N., Sindiri, S., Sachs, A., et al. (2022). Molecular signatures of antitumor neoantigen-reactive T cells from metastatic human cancers. Science 375, 877–884. 10.1126/science.abl5447.

83. Hanada, K.-I., Zhao, C., Gil-Hoyos, R., Gartner, J.J., Chow-Parmer, C., Lowery, F.J., Krishna, S., Prickett, T.D., Kivitz, S., Parkhurst, M.R., et al. (2022). A phenotypic signature that identifies neoantigen-reactive T cells in fresh human lung cancers. Cancer Cell 40, 479–493.e6. 10.1016/j.ccell.2022.03.012.

84. Oliveira, G., Stromhaug, K., Klaeger, S., Kula, T., Frederick, D.T., Le, P.M., Forman, J., Huang, T., Li, S., Zhang, W., et al. (2021). Phenotype, specificity and avidity of antitumour CD8+ T cells in melanoma. Nature 596, 119–125. 10.1038/s41586-021-03704-y.

85. van der Leun, A.M., Thommen, D.S., and Schumacher, T.N. (2020). CD8+ T cell states in human cancer: insights from single-cell analysis. Nat Rev Cancer 20, 218–232. 10.1038/s41568-019-0235-4.

86. Tan, X.-L., Le, A., Scherrer, E., Tang, H., Kiehl, N., Han, J., Jiang, R., Diede, S.J., and Shui, I.M. (2022). Systematic literature review and meta-analysis of clinical outcomes and prognostic factors for melanoma brain metastases. Frontiers in Oncology 12.

87. Achrol, A.S., Rennert, R.C., Anders, C., Soffietti, R., Ahluwalia, M.S., Nayak, L., Peters, S., Arvold, N.D., Harsh, G.R., Steeg, P.S., et al. (2019). Brain metastases. Nat Rev Dis Primers 5, 5. 10.1038/s41572-018-0055-y.

88. Lee, J.E., and Yang, S.H. (2023). Advances in Brain Metastasis Models. Brain Tumor Res Treat 11, 16–21. 10.14791/btrt.2022.0037.

89. Jansen, C.S., Prabhu, R.S., Pagadala, M.S., Chappa, P., Goyal, S., Zhou, C., Neill, S.G., Prokhnevska, N., Cardenas, M., Hoang, K.B., et al. (2023). Immune niches in brain metastases contain TCF1+ stem-like T cells, are associated with disease control and are modulated by preoperative SRS. Res Sq, rs.3.rs-2722744. 10.21203/rs.3.rs-2722744/v1.

90. Zakaria, R., Platt-Higgins, A., Rathi, N., Radon, M., Das, S., Das, K., Bhojak, M., Brodbelt, A., Chavredakis, E., Jenkinson, M.D., et al. (2018). T-Cell Densities in Brain Metastases Are Associated with Patient Survival Times and Diffusion Tensor MRI Changes. Cancer Res 78, 610–616. 10.1158/0008-5472.CAN-17-1720.

91. Harter, P.N., Bernatz, S., Scholz, A., Zeiner, P.S., Zinke, J., Kiyose, M., Blasel, S., Beschorner, R., Senft, C., Bender, B., et al. (2015). Distribution and prognostic relevance of tumor-infiltrating lymphocytes (TILs) and PD-1/PD-L1 immune checkpoints in human brain metastases. Oncotarget 6. 10.18632/oncotarget.5696.

92. Berghoff, A.S., Ricken, G., Wilhelm, D., Rajky, O., Widhalm, G., Dieckmann, K., Birner, P., Bartsch, R., and Preusser, M. (2016). Tumor infiltrating lymphocytes and PD-L1 expression in brain metastases of small cell lung cancer (SCLC). J Neurooncol 130, 19–29. 10.1007/s11060-016-2216-8.

93. Schetters, S.T.T., Gomez-Nicola, D., Garcia-Vallejo, J.J., and Van Kooyk, Y. (2018). Neuroinflammation: Microglia and T Cells Get Ready to Tango. Front Immunol 8, 1905. 10.3389/fimmu.2017.01905.

94. Rodríguez, A.M., Rodríguez, J., and Giambartolomei, G.H. (2022). Microglia at the Crossroads of Pathogen-Induced Neuroinflammation. ASN Neuro 14, 17590914221104566. 10.1177/17590914221104566.

95. Chen, X., Firulyova, M., Manis, M., Herz, J., Smirnov, I., Aladyeva, E., Wang, C., Bao, X., Finn, M.B., Hu, H., et al. (2023). Microglia-mediated T cell Infiltration Drives Neurodegeneration in Tauopathy. Nature 615, 668–677. 10.1038/s41586-023-05788-0.

96. Chen, D., Varanasi, S.K., Hara, T., Traina, K., Sun, M., McDonald, B., Farsakoglu, Y., Clanton, J., Xu, S., Garcia-Rivera, L., et al. (2023). CTLA-4 blockade induces a microglia-Th1 cell partnership that stimulates microglia phagocytosis and anti-tumor function in glioblastoma. Immunity 56, 2086–2104.e8. 10.1016/j.immuni.2023.07.015.

97. von Roemeling, C.A., Patel, J.A., Carpenter, S.L., Yegorov, O., Yang, C., Bhatia, A., Doonan, B.P., Russell, R., Trivedi, V.S., Klippel, K., et al. (2024). Adeno-associated virus delivered CXCL9 sensitizes glioblastoma to anti-PD-1 immune checkpoint blockade. Nat Commun 15, 5871. 10.1038/s41467-024-49989-1.

98. Wang, L., Li, C., Zhan, H., Li, S., Zeng, K., Xu, C., Zou, Y., Xie, Y., Zhan, Z., Yin, S., et al. (2024). Targeting the HSP47-collagen axis inhibits brain metastasis by reversing M2 microglial polarization and restoring anti-tumor immunity. CR Med 5. 10.1016/j.xcrm.2024.101533.

99. Feng, Y., Hu, X., Zhang, Y., and Wang, Y. (2024). The Role of Microglia in Brain Metastases: Mechanisms and Strategies. Aging Dis 15, 169–185. 10.14336/AD.2023.0514.

100. Nguyen, L.T., Aprico, A., Nwoke, E., Walsh, A.D., Blades, F., Avneri, R., Martin, E., Zalc, B., Kilpatrick, T.J., and Binder, M.D. (2023). Mertk-expressing microglia influence oligodendrogenesis and myelin modelling in the CNS. Journal of Neuroinflammation 20, 253. 10.1186/s12974-023-02921-8.

101. Gilchrist, S.E., Goudarzi, S., and Hafizi, S. (2020). Gas6 Inhibits Toll-Like Receptor-Mediated Inflammatory Pathways in Mouse Microglia via Axl and Mer. Front Cell Neurosci 14, 576650. 10.3389/fncel.2020.576650.

102. Moertel, C.L., Xia, J., LaRue, R., Waldron, N.N., Andersen, B.M., Prins, R.M., Okada, H., Donson, A.M., Foreman, N.K., Hunt, M.A., et al. (2014). CD200 in CNS tumor-induced immunosuppression: the role for CD200 pathway blockade in targeted immunotherapy. J Immunother Cancer 2, 46. 10.1186/s40425-014-0046-9.

103. Ngwa, C., and Liu, F. (2019). CD200-CD200R signaling and diseases: a potential therapeutic target? Int J Physiol Pathophysiol Pharmacol 11, 297–309.

104. Timmerman, R., Zuiderwijk-Sick, E.A., Baron, W., and Bajramovic, J.J. (2023). In silico-in vitro modeling to uncover cues involved in establishing microglia identity: TGF-β3 and laminin can drive microglia signature gene expression. Front Cell Neurosci 17, 1178504. 10.3389/fncel.2023.1178504.

105. Rigby, M.J., Gomez, T.M., and Puglielli, L. (2020). Glial Cell-Axonal Growth Cone Interactions in Neurodevelopment and Regeneration. Front Neurosci 14, 203. 10.3389/fnins.2020.00203.

106. Lee, W.S., Lee, W.-H., Bae, Y.C., and Suk, K. (2019). Axon Guidance Molecules Guiding Neuroinflammation. Exp Neurobiol 28, 311–319. 10.5607/en.2019.28.3.311.

107. Tang, J., Jin, Y., Jia, F., Lv, T., Manaenko, A., Zhang, L.-F., Zhang, Z., Qi, X., Xue, Y., Zhao, B., et al. (2023). Gas6 Promotes Microglia Efferocytosis and Suppresses Inflammation Through Activating Axl/Rac1 Signaling in Subarachnoid Hemorrhage Mice. Transl Stroke Res 14, 955–969. 10.1007/s12975-022-01099-0.

108. Fazio, F., Ulivieri, M., Volpi, C., Gargaro, M., and Fallarino, F. (2018). Targeting metabotropic glutamate receptors for the treatment of neuroinflammation. Current Opinion in Pharmacology 38, 16–23. 10.1016/j.coph.2018.01.010.

109. Szepesi, Z., Manouchehrian, O., Bachiller, S., and Deierborg, T. (2018). Bidirectional Microglia-Neuron Communication in Health and Disease. Front Cell Neurosci 12, 323. 10.3389/fncel.2018.00323.

110. Colonna, M. (2023). The biology of TREM receptors. Nat Rev Immunol 23, 580–594. 10.1038/s41577-023-00837-1.

111. Yin, R.-H., Yu, J.-T., and Tan, L. (2015). The Role of SORL1 in Alzheimer’s Disease. Mol Neurobiol 51, 909–918. 10.1007/s12035-014-8742-5.

112. Rogaeva, E., Meng, Y., Lee, J.H., Gu, Y., Kawarai, T., Zou, F., Katayama, T., Baldwin, C.T., Cheng, R., Hasegawa, H., et al. (2007). The neuronal sortilin-related receptor SORL1 is genetically associated with Alzheimer disease. Nat Genet 39, 168–177. 10.1038/ng1943.

113. Kleffman, K., Levinson, G., Rose, I.V.L., Blumenberg, L.M., Shadaloey, S.A.A., Dhabaria, A., Wong, E., Galán-Echevarría, F., Karz, A., Argibay, D., et al. (2022). Melanoma-Secreted Amyloid Beta Suppresses Neuroinflammation and Promotes Brain Metastasis. Cancer Discovery 12, 1314–1335. 10.1158/2159-8290.CD-21-1006.

114. So, J.-S., Kim, H., and Han, K.-S. (2021). Mechanisms of Invasion in Glioblastoma: Extracellular Matrix, Ca2+ Signaling, and Glutamate. Frontiers in Cellular Neuroscience 15. 10.3389/fncel.2021.663092.

115. Huang, Q., Chen, L., Liang, J., Huang, Q., and Sun, H. (2022). Neurotransmitters: Potential Targets in Glioblastoma. Cancers (Basel) 14, 3970. 10.3390/cancers14163970.

116. Bankhead, P., Loughrey, M.B., Fernández, J.A., Dombrowski, Y., McArt, D.G., Dunne, P.D., McQuaid, S., Gray, R.T., Murray, L.J., Coleman, H.G., et al. (2017). QuPath: Open source software for digital pathology image analysis. Sci Rep 7, 16878. 10.1038/s41598-017-17204-5.

117. Virador, V., Matsunaga, N., Matsunaga, J., Valencia, J., Oldham, R.J., Kameyama, K., Peck, G.L., Ferrans, V.J., Vieira, W.D., Abdel-Malek, Z.A., et al. (2001). Production of Melanocyte-Specific Antibodies to Human Melanosomal Proteins: Expression Patterns in Normal Human Skin and in Cutaneous Pigmented Lesions. Pigment Cell Research 14, 289–297. 10.1034/j.1600-0749.2001.140410.x.

118. Korin, B., Dubovik, T., and Rolls, A. (2018). Mass cytometry analysis of immune cells in the brain. Nat Protoc 13, 377–391. 10.1038/nprot.2017.155.

119. Araya, R.E., and Goldszmid, R.S. (2020). Characterization of the tumor immune infiltrate by multiparametric flow cytometry and unbiased high-dimensional data analysis. Methods Enzymol 632, 309–337. 10.1016/bs.mie.2019.11.012.

120. Ashhurst, T.M., Marsh-Wakefield, F., Putri, G.H., Spiteri, A.G., Shinko, D., Read, M.N., Smith, A.L., and King, N.J.C. (2022). Integration, exploration, and analysis of high-dimensional single-cell cytometry data using Spectre. Cytometry Part A 101, 237–253. 10.1002/cyto.a.24350.

121. Hao, Y., Hao, S., Andersen-Nissen, E., Mauck, W.M., Zheng, S., Butler, A., Lee, M.J., Wilk, A.J., Darby, C., Zager, M., et al. (2021). Integrated analysis of multimodal single-cell data. Cell 184, 3573–3587.e29. 10.1016/j.cell.2021.04.048.

122. McInnes, L., Healy, J., Saul, N., and Großberger, L. (2018). UMAP: Uniform Manifold Approximation and Projection. Journal of Open Source Software 3, 861. 10.21105/joss.00861.

123. Hao, Y., Stuart, T., Kowalski, M.H., Choudhary, S., Hoffman, P., Hartman, A., Srivastava, A., Molla, G., Madad, S., Fernandez-Granda, C., et al. (2024). Dictionary learning for integrative, multimodal and scalable single-cell analysis. Nat Biotechnol 42, 293–304. 10.1038/s41587-023-01767-y.

124. Ianevski, A., Giri, A.K., and Aittokallio, T. (2022). Fully-automated and ultra-fast cell-type identification using specific marker combinations from single-cell transcriptomic data. Nat Commun 13, 1246. 10.1038/s41467-022-28803-w.

125. Aibar, S., González-Blas, C.B., Moerman, T., Huynh-Thu, V.A., Imrichova, H., Hulselmans, G., Rambow, F., Marine, J.-C., Geurts, P., Aerts, J., et al. (2017). SCENIC: Single-cell regulatory network inference and clustering. Nature methods 14, 1083. 10.1038/nmeth.4463.

126. Jin, S., Guerrero-Juarez, C.F., Zhang, L., Chang, I., Ramos, R., Kuan, C.-H., Myung, P., Plikus, M.V., and Nie, Q. (2021). Inference and analysis of cell-cell communication using CellChat. Nat Commun 12, 1088. 10.1038/s41467-021-21246-9.

